# HIV-1 provirus transcription and translation in macrophages differs from pre-integrated cDNA complexes and requires E2F transcriptional programs

**DOI:** 10.1101/2021.04.21.440655

**Authors:** Albebson L. Lim, Philip Moos, Christopher D. Pond, Erica C. Larson, Laura J. Martins, Matthew A Szaniawski, Vicente Planelles, Louis R. Barrows

## Abstract

HIV-1 cDNA pre-integration complexes have been shown to persist for weeks in macrophages and to be transcriptionally active. Early and late gene transcripts are produced, along with some viral proteins, yet whole virus is not. While previous work has focused on the transcription and translation of HIV-1 genes; our understanding of cellular milieu that accompanies viral production is incomplete. We have used an *in vitro* system to model HIV-1 infection of macrophages, and single cell RNA sequencing (scRNA-seq) to compare the transcriptomes of uninfected cells, cells harboring pre-integration HIV-1 complexes (PIC) and those containing integrated provirus and actively making late HIV proteins. These are also compared to control cells, not exposed to virus.

Several observations provide new perspective on the effects of HIV-1 transcription from pre-integrated cDNA versus from integrated provirus. First, HIV-1 transcript levels do not necessarily correlate with virus production, cells harboring PIC cDNA have transcript loads comparable to cells transcribing from provirus and making p24, mCherry, and vpu proteins. Second, all HIV-1 transcripts are easily detectable in abundance from PIC cDNA transcription, as is the case with cells transcribing from provirus, although the frequency of PIC cells with detectable gag-pol, tat, env, and nef transcripts is higher than the corresponding frequencies observed for “Provirus cells”. Third, the background transcriptomes of cells harboring pre-integrated HIV-1 cDNA are not otherwise detectably altered from cells not containing any HIV- 1 transcript. Fourth, integration and production of p24, mCherry, and Vpu proteins is accompanied by a switch from transcriptomes characterized by NFkB and AP-1 promoted transcription to a transcriptome characterized by E2F family transcription products. While some of these observations may seem heretical, single cell analysis provides a more nuanced understanding of PIC cDNA transcription and the transcriptomic changes that support HIV-1 protein production from integrated provirus.

**Author Summary:** Single cell analysis is able to distinguish between HIV-1 infected macrophage cells that are transcribing pre-integrated HIV-1 cDNA and those transcribing HIV-1 provirus. Only cells transcribing HIV-1 provirus are making p24, marker mCherry and Vpu proteins, which corresponds with a change in the host cell’s background transcriptome from one expressing viral restriction and immunological response genes to one that is expressing genes associated with cell replication and oxidative phosphorylation.

## Introduction

The major barrier to curing HIV-1 in patients is a small reservoir of cells that are latently infected and impervious to immune recognition and clearance [1–3]. The study of HIV-1 latency is complicated by the fact that latently infected cells *in vivo* are extremely rare. It is a drawback that many studies of latency have relied on bulk sequencing endpoints. Under these conditions, the specific parameters defining the latently infected cell are diluted in the context of a vast heterogeneous population. In addition, multiple mechanisms can result in latency reversal and therefore one latently infected cell may differ from the next (reviewed in ref. [4]). Thus, averaging data from a heterogeneous cell population, such as data obtained by bulk sequencing studies, leads to confusion rather than clarification.

Following infection and reverse transcription, the pre-integration cDNA complex (PIC), in which the HIV-1 genome may take linear or circular forms, serves as a template for transcription [5–7]. In dividing T cells, the PIC is short-lived, and the transient transcription of genes from the PIC is considered irrelevant [6]. However, in quiescent T cells the PIC has been shown to be longer lived and even results in sufficient transcription for virion production in response to latency reversal agents [8]. In macrophages the dynamics of PIC integration are different [9–12]. Macrophages have been shown to harbor transcriptionally active pre-integrated HIV-1 cDNA for months [10, 11], however this PIC mRNA is not thought to result in the production of virus [12].

We have recently used single cell RNA sequencing (scRNA-seq) techniques [13, 14] to show that in an activated monocytic cell line, most of the cells in a culture infected with HIV-1 are in fact producing HIV transcripts, while only a minority are producing Gag/p24 and other proteins. Levels of HIV-1 transcript do not correlate with virus production, since many of the cells harboring PIC complexes have transcript loads comparable to cells making p24. This means that a high load of viral transcript is not a sufficient switch to reverse latency. Overall, within the limitations of 10X Genomics technology [15], transcripts for gag-pol, tat, env, and nef are found in higher numbers of PIC cells, compared to cells transcribing from provirus. However, the loads of these transcripts detected in PIC and provirus cells is similar. Transcripts for gag, vif, vpr, rev, vpu, and the marker gene mCherry are detectable in relatively equivalent percentages of cells transcribing from provirus or PIC, and at similar levels per cell. In no case did “Provirus” cells (those transcribing from integrated HIV-1 cDNA and producing gag proteins) appear to be producing any of the detected HIV-1 transcripts at a higher prevalence, or at higher concentrations per cell.

Quite notably, the background transcriptomes of cells harboring HIV-1 PIC are not detectably altered by PIC transcription. Unsupervised clustering shows cells containing PIC transcripts to be distributed equally throughout the PIC/Bystander cell cluster. “Bystander” cells are defined as those cells not containing any detectable HIV-1 transcript. Thus, cells harboring HIV-1 PIC appear “oblivious” to the presence of HIV-1 gene transcription, even though some have been reported to be producing detectable levels of Nef, other HIV-1 proteins and chemokines [9, 10, 15–19].

Transcription Factor Targeting analysis [20–22] shows that NFkB and AP-1 transcription products are predominant in the transcriptomes of PIC and Bystander cells. In contrast, Provirus cluster cells, on average, contain higher total amounts of viral transcripts and their transcriptomes predominantly exhibit E2F promoter family transcription products. These cells make detectable amounts of p24, mCherry and Vpu proteins. This seems counter-intuitive because NFkB and AP-1 are transcription sites in the HIV-1 LTR that promote transcription from the provirus [23, 24]. In addition, E2F has been reported to suppress HIV-1 transcription [25, 26]. Nevertheless, in our model it is clear that when cells are making late HIV-1 gene proteins, the transcriptomes exhibit activation of E2F regulated transcripts, while NFkB and AP- 1 regulated genes are relatively suppressed. Western blotting data agrees with the transcription factor analysis in that Rb phosphorylation is increased in the Provirus cluster cells, which correlates with increased E2F activation.

Western blotting also shows that cells transcribing from integrated provirus are making other HIV-1 proteins. Synthesis of p24 and Gag protein was detected in these cell preparations, as was mCherry, in positive correlation with flow cytometry results. Vpu protein was also detected only in protein preparations containing Provirus cluster cells.

E2F domination of the transcriptome is accompanied by activation of regulomes associated with cell division and RNA processing [27, 28]. The idea that the general cell transcriptome must transition to support virus production, indicated by E2F promoter family signaling, is supported by an experiment that shows that cells already making virus are 2 to 5 times more likely to make virus on a second round of infection than PIC or Bystander cells in the same culture. This was also observed in primary macrophage and T cell cultures, suggesting that a cell’s transcriptional background, in general, is a key factor in determining if virus will be produced or not.

## Results

### DHIV3 infection of activated THP-1 macrpohages yields cells with transcriptomes containing DHIV3 mRNA but otherwise identical to uninfected cells

We used a VSV-G-pseudotyped DHIV3 virus that expressed mCherry to promote consistent levels of viral entry in PMA-activated THP-1 cells and to allow for easier interpretation of post-cell entry events [29, 30] (Fig. S-1). Following infection, flow cytometry data clearly revealed two clusters of THP-1 cells, mCherry positive cells (usually from 2 to 20 percent of the total population, depending on DHIV3 titer) and mCherry negative cells (Fig. 1).

**Figure 1.**
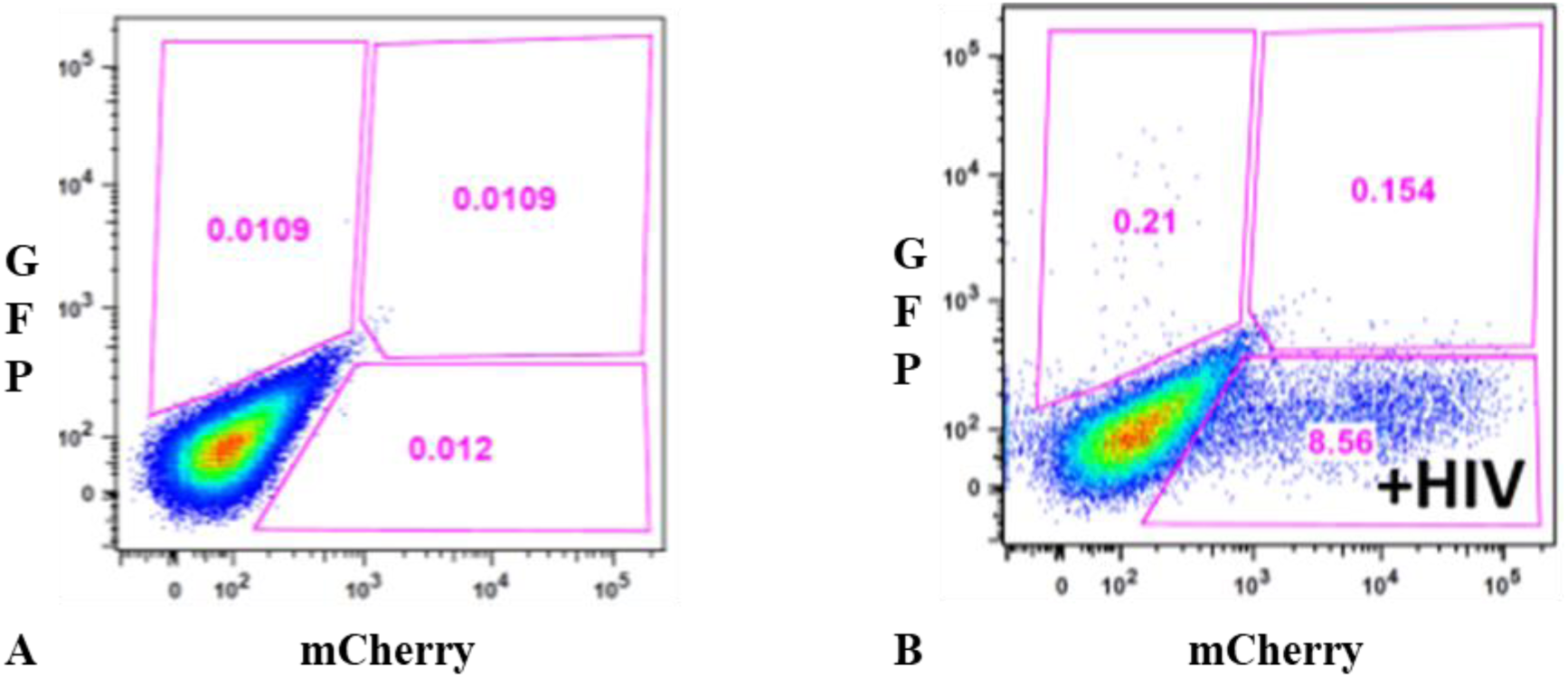
Flow cytometry analysis of DHIV3-mCherry infected THP-1 cells. Panel **A**) mock infection of PMA activated THP-1 cells. Panel **B**) PMA activated THP-1 cells infected with DHIV3-mCherry. Absicca, mCherry (Texas Red) emission. Ordinate, GFP (FITC) emission. mCherry positive cells equal approximately 8.5% of the total viable cell population.

Quantifying DHIV3 infection by scRNA-seq closely agreed with the flow cytometry data in that two clusters of cells were clearly identified, based on an individual cell’s background transcriptome, in percentages that agreed with flow cytometry of replicate cultures (Fig. 2). Data analysis is described in detail in the Methods. Briefly, raw fastq files were generated, aligned to a custom reference genome (GRCh38 augmented with mCherry and HIV genes) and per cell gene counts generated with the 10X Cell Ranger pipeline. Following basic QC and filtering as suggested by Seurat, we generated both UMAP and t-SNE clustering projections [15]. Library construction targeted 5,000 cells and routinely yielded greater than 4,000 cells following Seurat analysis and quality control. Library construction and sequencing experiments were performed with technical repeats. The technical repeats were found to be statistically identical and were therefore combined (Fig. S-2).

**Figure 2.**
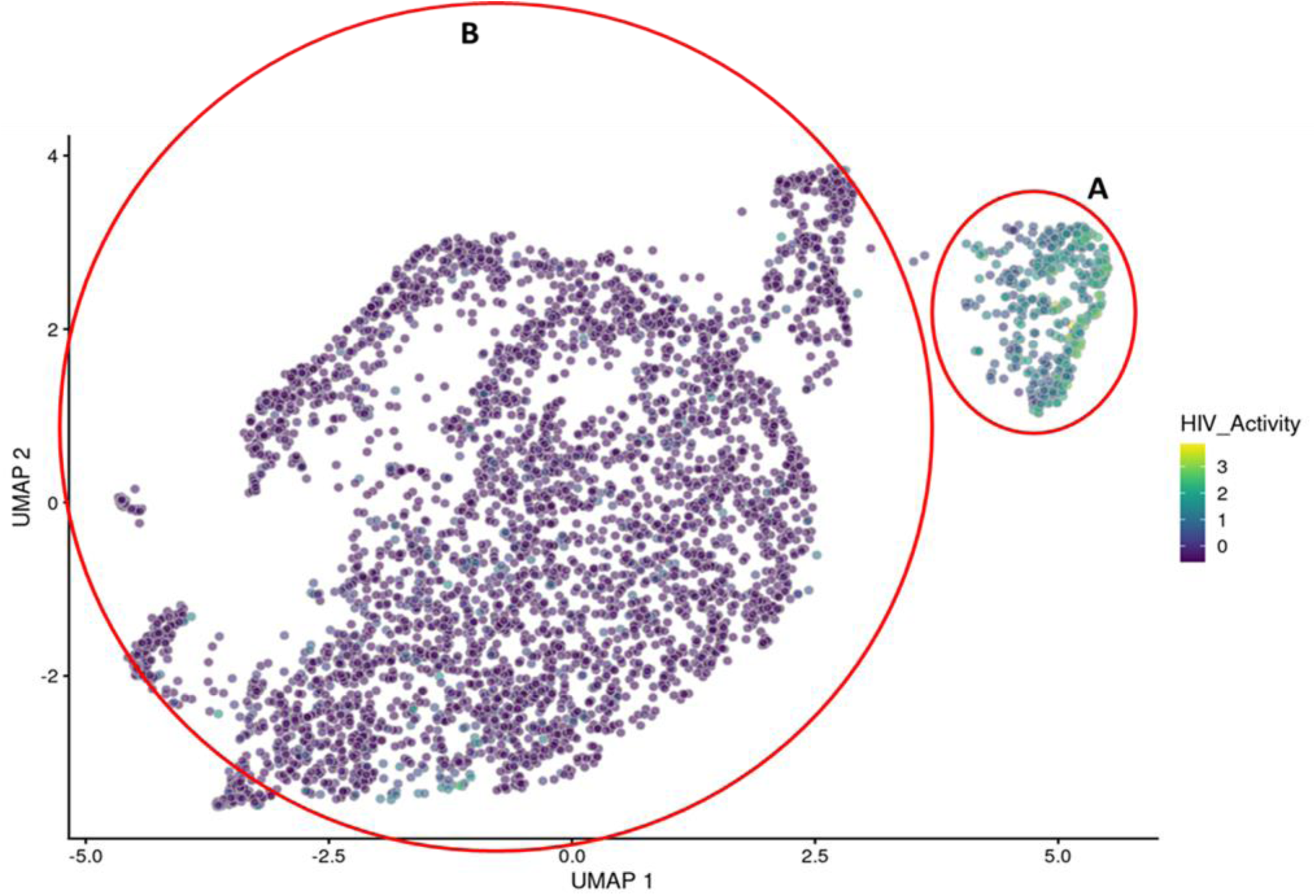
UMAP projection of scRNA seq data from experiment HIVreplicate1. Greater than 14,000 different cellular genes were detected in this analysis, including the 9 viral genes and mCherry message originating from DHIV3-mCherry. A semi-supervised two cluster model was adopted, the smaller “Provirus cluster” (cluster A) was 8.1% of the total cell population, approximately equivalent to the percentage of mCherry positive cells from Fig. 1. The two semi- supervised clusters are circled in red. PIC/Bystander cluster is indicated as cluster B. The HIV activity scale presents the Seurat module score that is described in methods. Input data in this analysis included 33,819 PCA entries. The number of cells with detected genes following Seurat QC in the PIC/Bystander cluster was 12% of the total population. Bar codes of the same cells tracked to the Provirus clusters, regardless of whether the clusters were generated using UMAP or Seurat-tSne tools (Fig. S-3).

Following Seurat analysis the number of mean reads per cell was approximately 35,000, with a median of more than 2,500 genes detected per transcriptome, greater than 15,000 different gene transcripts detected in the overall library. Control cultures yielded slightly larger numbers of cells captured, with greater than 6,000 cells each and with more than 3,000 gene detected per cell. All experiments were conducted with parallel cultures that were analyzed by flow cytometry. In the HIVreplicate1 experiment, flow cytometry indicated 8.6 percent mCherry positive cells (Fig.1).

Figure 2 shows the scRNA-seq UMAP analyses of HIVreplicate1 (t-Sne projection shown in Fig. S-3). When we quantified HIV-1 transcripts in individual cells, we found that cells in the smaller, “Provirus” cluster (defined as what we now understand to be cells transcribing provirus template) were characterized by transcriptomes generally containing higher minimum levels of HIV-1 transcripts. This “Provirus” cluster accounted for approximately 8.1 % of the total cell population, and closely matched the percentage of cells identified as mCherry positive by flow cytometry in the duplicate culture, shown above. In the second “PIC/Bystander” cluster of mCherry negative cells, we found that many of the macrophages (more than 50%) were actually producing HIV-1 transcript (Fig. S-3 plots cells with any detected HIV-1 transcript). We now understand that these cells are “PIC” cells, defined as cells containing HIV-1 mRNA transcribed from pre-integrated cDNA complexes. Some PIC cell transcriptomes appeared to contain HIV-1 transcript loads as high as many cells detected in the Provirus cluster, but most expressed lower amounts (Fig. 2; S-3). Remarkably, these PIC cells had no other statistically significant changes to their transcriptomes that would differentiate them from the truly uninfected “Bystander” cells (defined as cells in the PIC/Bystander cluster not containing detectable HIV-1 mRNA). Thus, PIC and Bystander cells made up the “PIC/Bystander” cluster.

We used a Feature map to plot the influence of cell cycle, number of genes detected or mitochondrial transcript number on the distribution of cells containing HIV-1 transcripts (Fig 3 B-D). None of these factors had any significant influence of the distribution of PIC cells throughout the PIC/Bystander cluster. We then used unsupervised clustering and violin plots of HIV transcript load to examine the distribution of cells containing HIV-1 transcripts (Fig. 4) throughout the Provirus and PIC/Bystander clusters. Using K-nearest neighbors clustering, at K10, 2 clusters of cells (clusters 6 and 8) were identified that accounted for most of the Provirus cluster cells (372 of the 380 cells), as determined by semi supervised clustering shown above (Fig. 2). Therefore, the combined transcriptomes of cells in clusters 6 and 8 were compared to the combined transcriptomes of cells in clusters 1-5, 7 and 9-10, and used to generate the DGE analyses shown below. Unsupervised clustering using a range of designated K values from 10 to 130 is shown in Figure S-4.

**Figure 3.**
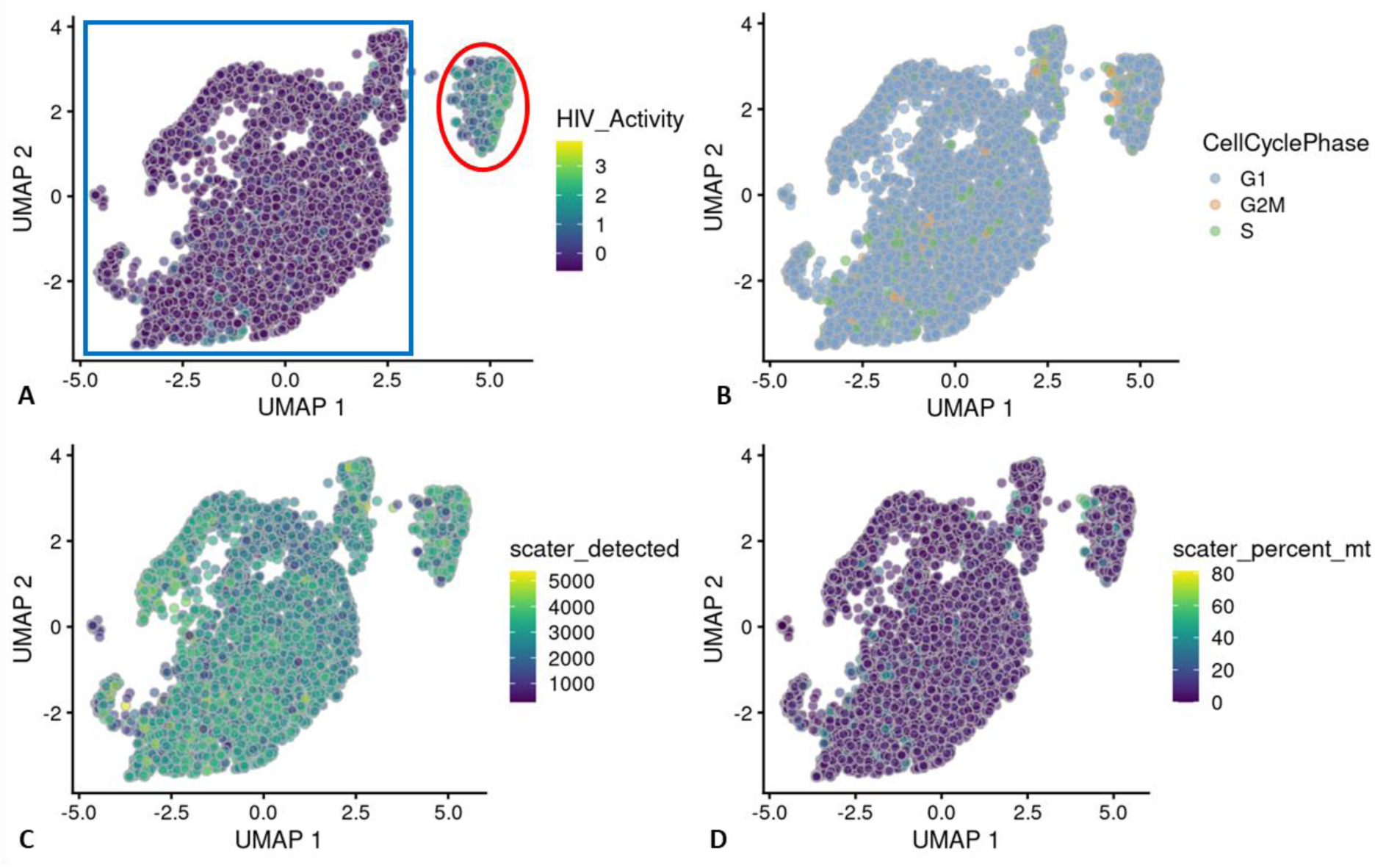
UMAP Feature plot of experiment HIVreplicate1. Panel **A)** Distribution of HIV-1 transcript positive cells shown in Fig. 2. Provirus cluster circled in red, PIC/Bystander cluster in the blue square. Panels **B**), **C**) and **D**), respectively, show no influence of cell cycle, number of genes detected per cell, or percent mitochondrial transcripts (positively correlated with cell stress) on the distribution of PIC cells (HIV-1 transcript containing cells) throughout the PIC/Bystander cluster.

**Figure. 4.**
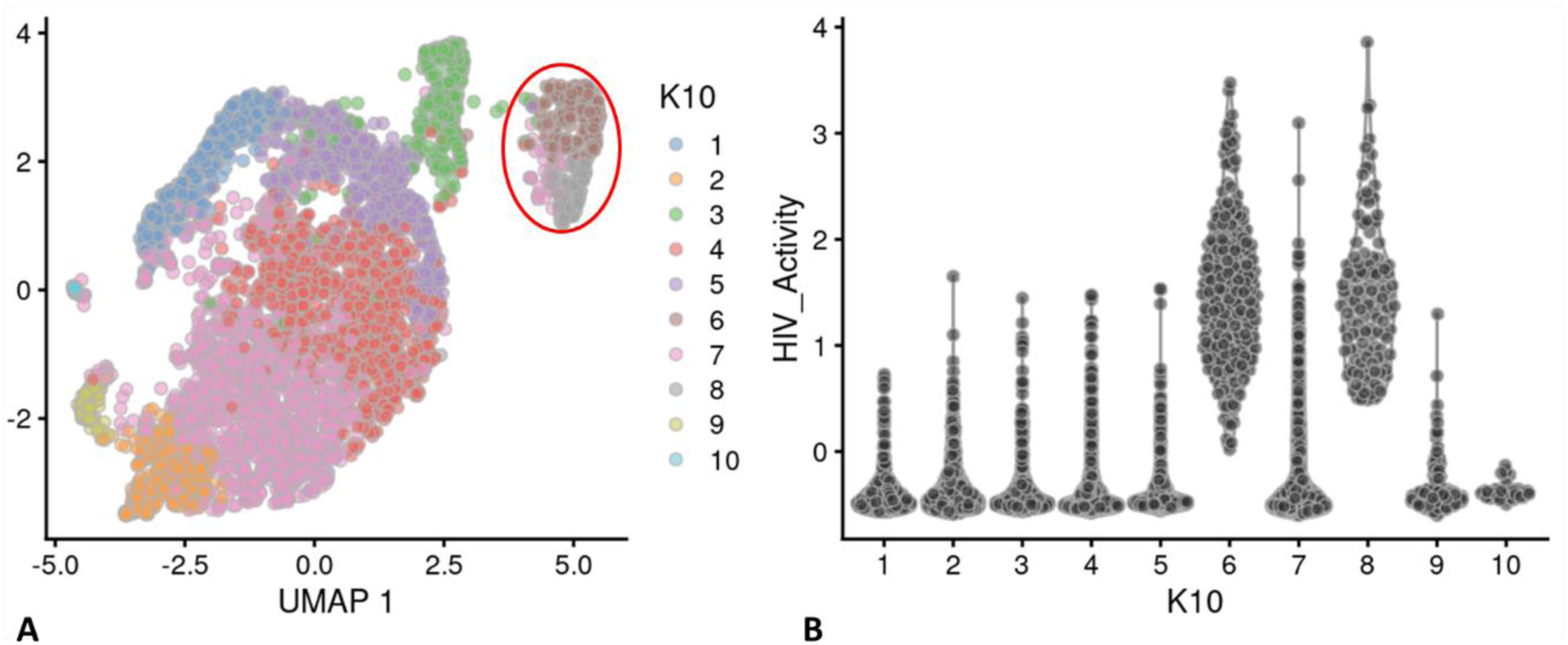
Unsupervised clustering of UMAP shown in. Fig 2. Panel **A**) shows unsupervised clustering obtained at K equals 10. Panel **B**) Violin plot of HIV-1 transcripts/cell in the 10 clusters identified at K10 (Scran’s buildSSNGraph using the PCA as input). PIC cells with detectable HIV-1 transcripts, were distributed throughout clusters 1-5, 7 and 9-10. Clusters 6 and 8 contained 372 of the 381 cells included in the semi-supervised Provirus cluster (circled in red). Stipulation of lower K values means that during analysis any one given cell is clustered with a smaller number of cells with similar transcriptomes.

Clusters 6 and 8 (Fig. 4B) were characterized as containing few cells with low HIV transcript load. Thus, cells in what we define as the Provirus cluster differ significantly from PIC cells in terms of average minimal DHIV3 transcript load. They also differ significantly in background transcriptome, as shown below. The multiple clusters generated in this unsupervised clustering are likely stochastic. Most importantly, they were not influenced by the presence or absence of PIC cells (Fig. 4B). PIC cells are distributed throughout all the PIC/Bystander clusters, regardless of the specified K value. So, for our purposes, the two cluster model shown in Figure 2 was considered to accurately represent the interpretation gained from flow cytometry (Fig. 1), namely: either the cells were making mCherry protein, or they were not.

Inferring from our flow cytometry percentages, we hypothesized that only cells in the Provirus cluster were making fluorescing mCherry protein. To confirm that cells making mCherry were also making late viral proteins, the production of p24 capsid protein was quantified using flow cytometry and monoclonal mouse IgG-AG3.0 anti-Gag p24 antibody.

Cells were fixed in formaldehyde, and secondary anti-mouse Alexa Fluor antibody allowed detection during flow cytometry. When cells were analyzed using FACS Canto, only cells making mCherry were found to stain for p24 (Fig. 5). This suggests that the presumed PIC cells are not making late viral proteins in detectable amounts, while only Provirus cells appear to be making late viral proteins.

**Figure 5.**
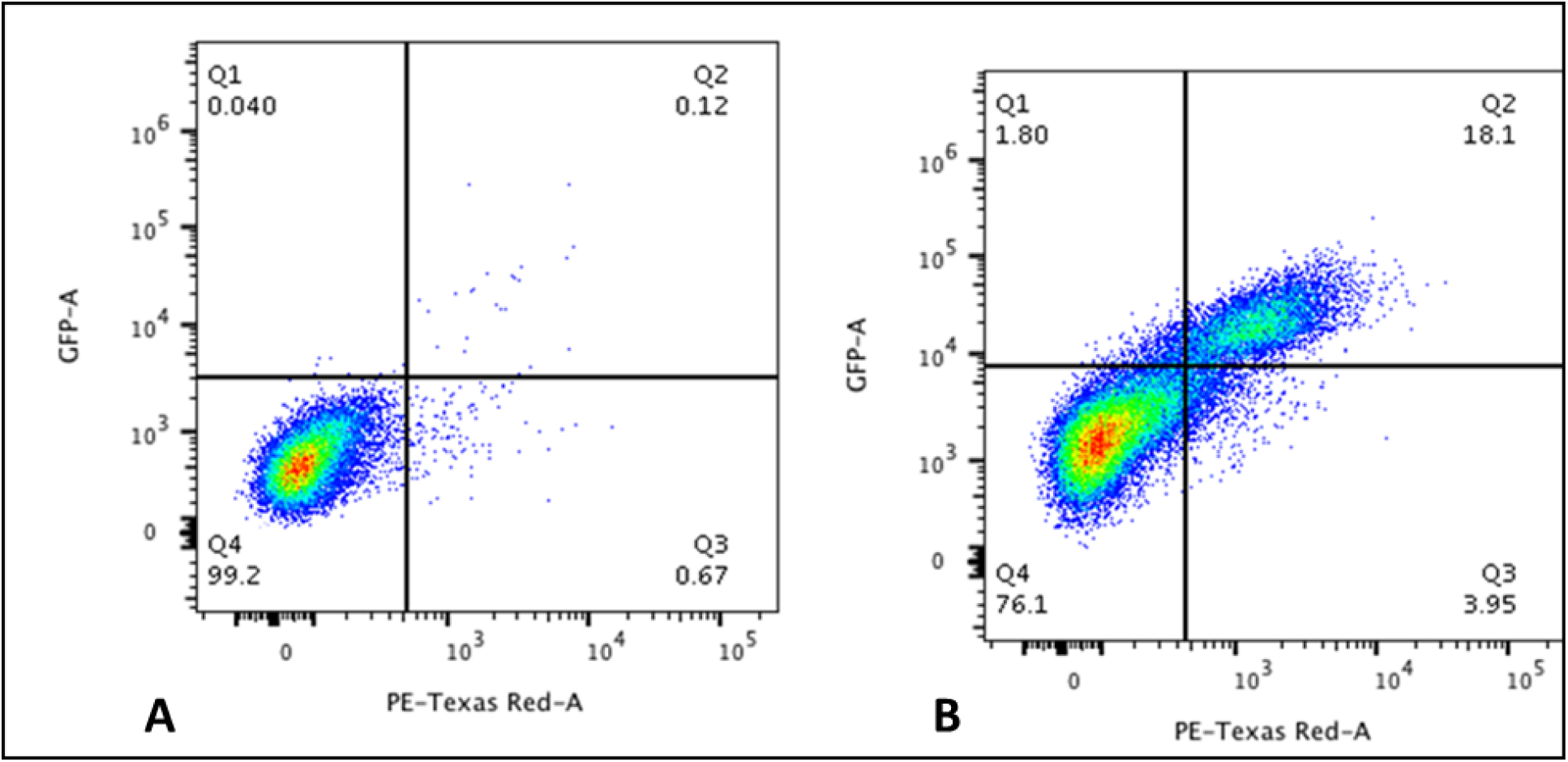
Flow cytometry analysis of DHIV3-mCherry infected THP-1 cells using p24 antibody. Panel **A**) mock infection. Panel **B**) mCherry expression was positively correlated with p24/Gag antigen detection by flow cytometry. Abscissa mCherry (Texas Red) emission. Ordinate, GFP (FITC) emission. mCherry/p24 positive cells equal approximately 18% of the total viable cell population in this experiment.

### The DHIV3 mRNA in “PIC” cells is due to PIC transcription

Integrase inhibitors have proven to be useful tools in the study of PIC transcription [16]. We used 25nM MK-2048 [31] for our experiments to confirm the “Provirus” versus “PIC” status of our cell clusters. When integrase inhibitor treated cultures were analyzed by flow cytometry, mCherry producing cells were routinely reduced from an average of about 12% to less than 2% of the total (Fig. 6). The use of the viability stain assured that this was not due to death of infected and drug-treated cells. We then analyzed protein preparations from parallel cultures of these cells by Western blotting. For the Western analysis, we initially purified populations of mCherry positive and mCherry negative cells using FACS analysis, collecting cells in the viable windows shown in Figure 6. However, because of the laborious difficulty in obtaining sufficient quantities of protein from the sorted cells, we resorted to comparing whole cultures of cells obtained either in the presence or in the absence of integrase inhibitor. We achieved >80% inhibition of mCherry production by integrase inhibitor treatment in these experiments. However, this means that in the integrase inhibitor treated/DHIV3 infected cell protein preparations there probably remained proteins from some mCherry positive cells, albeit in amounts greatly reduced from amounts present in the protein preparations from the corresponding DHIV3 infected cultures without integrase inhibitor treatment. Furthermore, in the DHIV3 infected cell preparations in the absence of integrase inhibitor, the majority of protein (>80%) still came from the mCherry negative cells. Both these protein preparations were compared to an equivalent amount of protein from Control cells. Control cell protein preparations were from PMA activated THP-1 cells not exposed to DHIV3.

**Figure 6.**
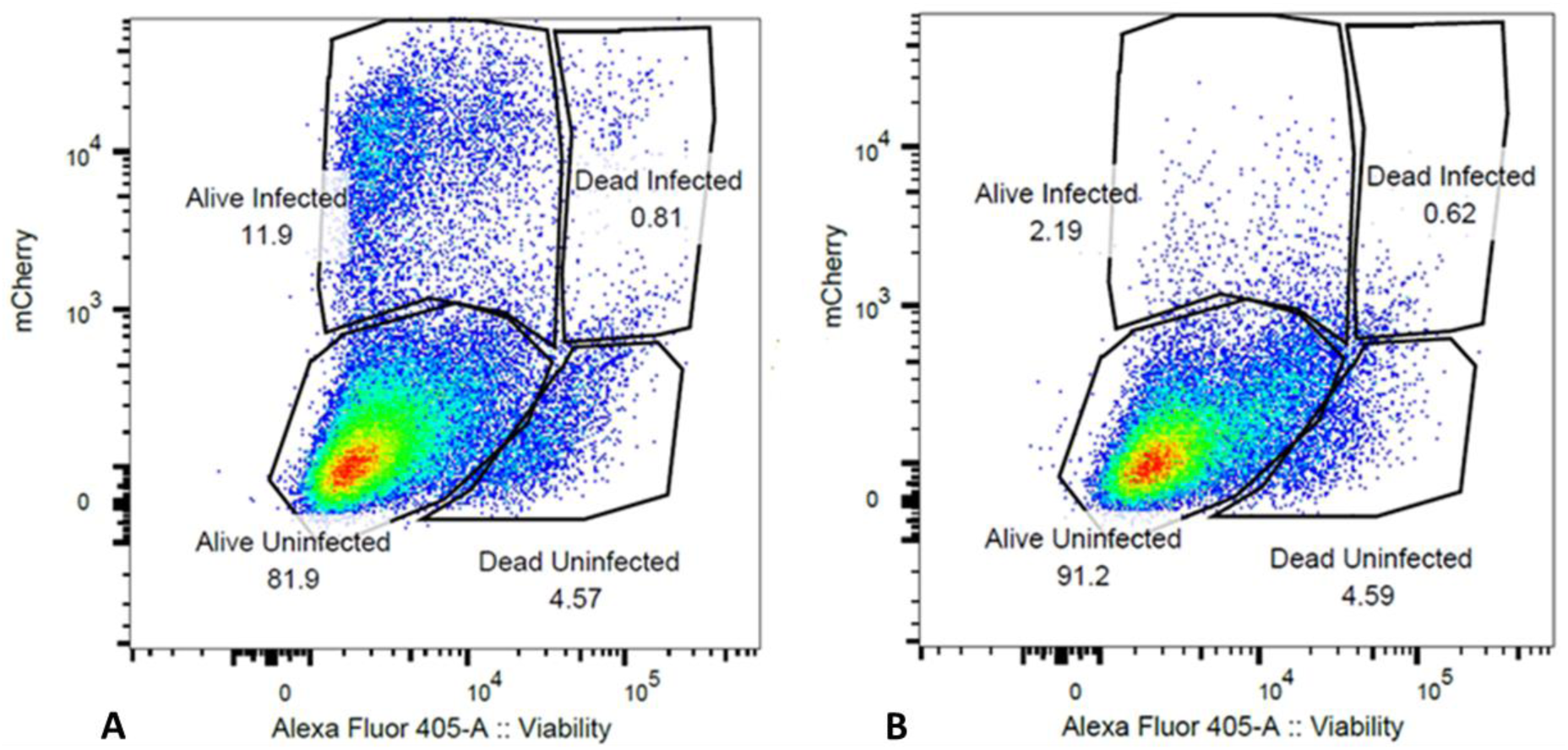
Integrase inhibitor treatment selectively reduces mCherry positive cells. Panel **A**) Flow cytometry analysis of DHIV3-mCherry infected THP-1 cells, versus viability stain. Abscissa shows viability stain intensity, ordinate shows mCherry intensity. Infected (Active), mCherry-producing cells account for approximately 12 % of the cell population. Panel **B**) Same as A except with the addition of 25nM MK-2048 integrase inhibitor at the time of infection. Integrase inhibitor effectively reduces the number of mCherry producing cells, without decreasing cell viability.

We found protein preparations from DHIV3-mCherry cultures in the absence of MK- 2048 to contain mCherry (Fig. 7 A). In addition, these same protein preparations contained p24 and p24 precursor proteins (Fig. 7 B). As expected, p24 protein was also detectable in protein from the integrase inhibitor treated culture. In the literature, this observation has been attributed to residual p24 protein lingering from the initial infection [10]. Fortunately, the p24 antibody we used is known to bind both p24 and Gag precursor proteins [10, 32]. We found both p24 and precursor p24 proteins only in the protein samples from cells not treated with integrase inhibitor. We obtained additional support for this interpretation by taking a 48 hr time point (Fig. 7D). The extended time point showed p24 precursor proteins only appearing in the integrase inhibitor cultures 48 hours after treatment. In contrast, p24 and Gag precursor proteins were increasingly detectable in cell proteins from integrase competent infections. Thus, p24 detected in DHIV3 infected cells was likely due to residual p24 from the original infection.

**Figure 7.**
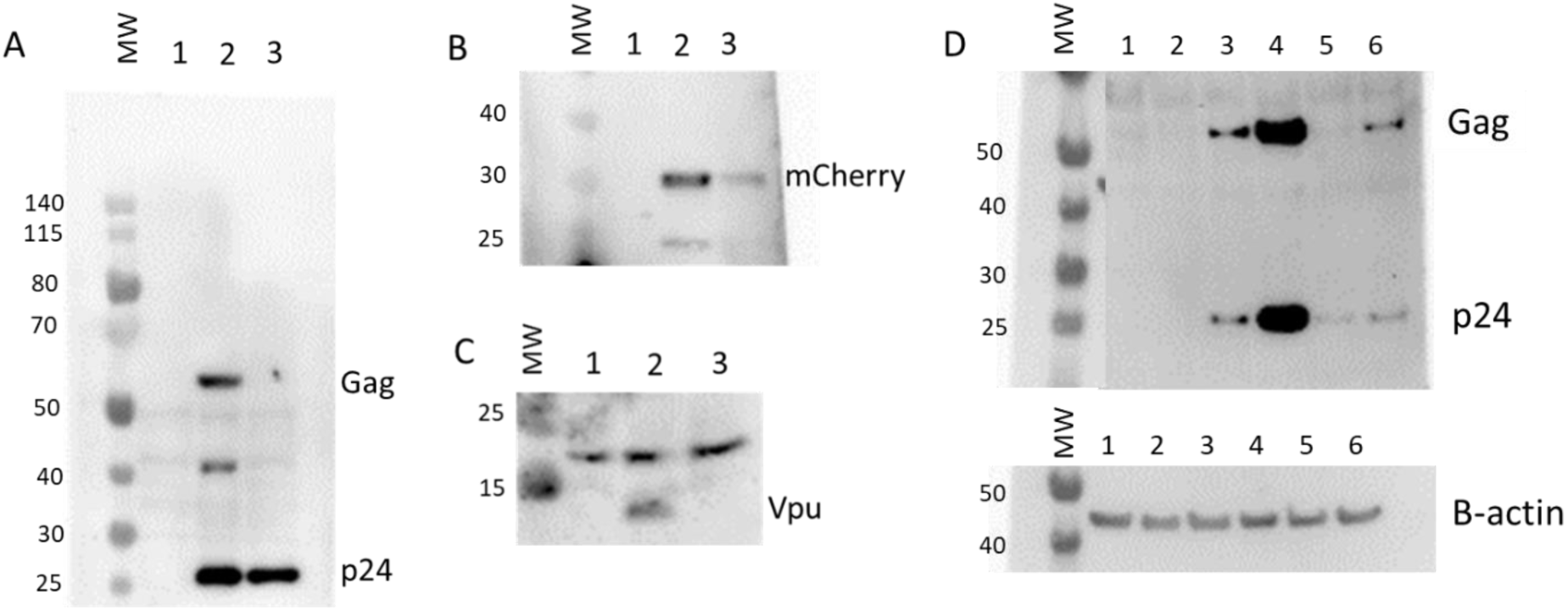
Effect of integrase inhibitor on mCherry, p24, Gag, and Vpu protein production in cultures containing DHIV3-mCherry infected cells. MW, molecular weight markers. Lane 1, Control cell protein; Lane 2, protein from DHIV3 infected culture; Lane 3, protein from DHIV3 infected cultures treated with integrase inhibitor (25nM MK-2048) as shown above in Figure 6. **A)** Lane 2, p24, and Gag precursor proteins visualized with p24 antibody used above in (Fig. 6), and HRP linked anti-mouse secondary antibody. The p24 band in lane 3 is residual from infection as reported in the literature [10]. The presence of precursor proteins in lane 2 shows Provirus synthesis in cultures containing Provirus cells. **B)** Antibody used was goat anti-mCherry, developed with HRP linked anti-goat secondary. mCherry protein clearly visible in preparations containing Provirus cell protein. **C)** Lane 2, Vpu detected with rabbit antibody, visualized using HRP linked anti-rabbit secondary. The resolution of the image slightly compromised due to the small size of Vpu protein. **D)** Lanes 1 and 2, Control cell protein at 24 and 48 hrs respectively; lanes 3 and 4, protein from DHIV3 infected culture at 24 and 48 hrs respectively; lanes 5 and 6, protein from DHIV3 infected cultures treated with integrase inhibitor (as above) at 24 and 48 hrs respectively. At 24 hrs post-infection, we only found both p24 and precursor Gag proteins in the protein samples from DHIV3 infected cells in the absence of integrase inhibitor. At 48 hrs post-infection, in the absence of integrase inhibitor, the amounts of detectable p24 and Gag proteins were dramatically increased from levels at 24 hrs post- infection. As seen initially (Panel A), some p24 protein was detectable in integrase inhibitor treated cultures at 24 hrs post-infection, however Gag is not detectable at this time. At 48 hrs post-infection in the integrase inhibitor treated-cultures, some Gag protein does becomes detectable, reflecting production in cells that escaped complete integrase inhibition. This is in agreement with our flow cytometry analysis that showed suppressed, but still detectable numbers of mCherry positive cells in the integrase inhibitor treated cultures. The Gag precursor proteins only appear in the integrase inhibitor treated culture proteins 48 hrs after treatment. All antibodies, sources, and dilutions are provided in Methods.

Further confirmation of the PIC versus Provirus status of our semi supervised cluster cell populations was obtained using real-time polymerase chain reaction (real-time PCR) analysis [33, 34]. DNA from the respective cell treatment groups described above (Control, DHIV3 infected, DHIV3 infected plus integrase inhibitor) was isolated and analyzed by PCR using multiple sets of primers. For detection of integrated proviral DNA a set of primers [34] was used to amplify from random nascent human genomic Alu sequences to an internal HIV LTR sequence. This initial amplification step was followed by a second PCR amplification step using nested primers, which would only amplify discrete DNA products that contain integrated provirus [34]. Evidence for integrated provirus was detected in much higher abundance from DNA of DHIV3 infected cells in the absence of integrase inhibitor (Fig S-5), although small amounts of integrated provirus DNA was detected in DHIV3-mCherry infected and integrase inhibitor treated cell culture DNA, when using higher amounts (200 ng) of starting DNA. This is in agreement with Flow cytometry results (Fig. 6) and Western analysis (Fig. 7) indicating small amounts of proviral DHIV3-mCherry DNA in our integrase inhibitor treated cultures.

To demonstrate unequivocally that our PIC cluster cells do indeed contain PIC HIV cDNA, we used the primers of Brussels and Sonigo [33], which are internal to the HIV-1 LTR sequence. These primers were oriented in a way so as to detect only circular 2-LTR PIC HIV-1 DNA by bridging the 2-LTR junction [33, 34]. We found that we required 2 rounds of PCR amplification to obtain the predicted 2-LTR amplicon, suggesting that this particular PIC species is in low abundance in our model cells. We tested this conclusion using bracketing primers (see Methods) to generate a product to contain the predicted amplicon of Brussels and Sonigo [33], and then followed with a round of amplification using the previous primers to generate a nested product. This approach also generated the predicted amplicon product from DHIV3-mCherry infected culture DNA, whether in the presence of integrase inhibitor or not. The predicted 2-LTR PIC cDNA PCR product was not detected in Control cell DNA.

To confirm that total PIC cDNA is in relatively high abundance in our DHIV3-mCherry infected cultures, we adapted previous primers to amplify total DHIV3 LTR DNA. With these primers, similarly high levels of total PIC cDNA was detectable in DHIV3 infected cells, whether in the presence of integrase inhibitor or not (Fig S-5).

Taken together, the Western blot and real-time PCR data confirm our flow cytometry observations that mCherry producing cells were also producing p24. The fact that the mCherry cells are selectively suppressed by integrase inhibitor treatment leads to the conclusion that only Provirus cluster cells are making p24 or mCherry proteins from the DHIV3 transcripts. Conversely, transcripts detected in PIC cells, due to PIC transcription, and do not lead to detectable mCherry or p24 synthesis.

We tried antibodies for all the other major HIV-1 proteins to test the correlation of protein expression with the detection of transcript. One antibody that yielded a clear result was the polyclonal antibody against Vpu. In this case, we see a result very similar to that obtained with mCherry and p24, in that protein was clearly detected in samples from infected cell cultures that were not treated with integrase inhibitor (Fig. 7). Vpu was not detectable in protein from the integrase inhibitor treated cells. It is an interesting side-note that Vpu has been associated with inhibition of NFkB promoted transcription [35, 36].

### scRNA-seq analysis of integrase inhibitor treated DHIV3 infected cultures

DHIV3 infected THP-1 cells were treated with 25nM MK-2048 at the time of infection, using our established protocol, and analyzed by scRNA-seq. In this experiment, the integrase inhibitor blocked ∼87.5% of mCherry production by flow cytometry analysis of a parallel culture (Fig. 6). The UMAP image of DHIV3 transcript features is shown in Figure 8A. Transcriptome clustering using varying nearest neighbor K-values (K10, K50, K90, and K130) yielded 3 to 7 clusters, depending on the K-value applied (S-6). Regardless of the specified K-value, we did not detect a Provirus cluster as was observed with integrase competent infections (Fig. 8C). Again, the distribution of HIV-1 transcript containing cells throughout the semi supervised integrase inhibitor single cell cluster was not effected by cell cycle, percent mitochondrial gene expression or numbers of genes detected per cell (Fig. S-7). This result confirms that integrase inhibitor-treatment selectively suppresses cells in the Provirus cluster, agreeing with results obtained by Western blot and PCR analysis, and confirms that the DHIV3 mRNA detected in the PIC cluster cells is due to transcription of PIC complexes.

**Figure 8.**
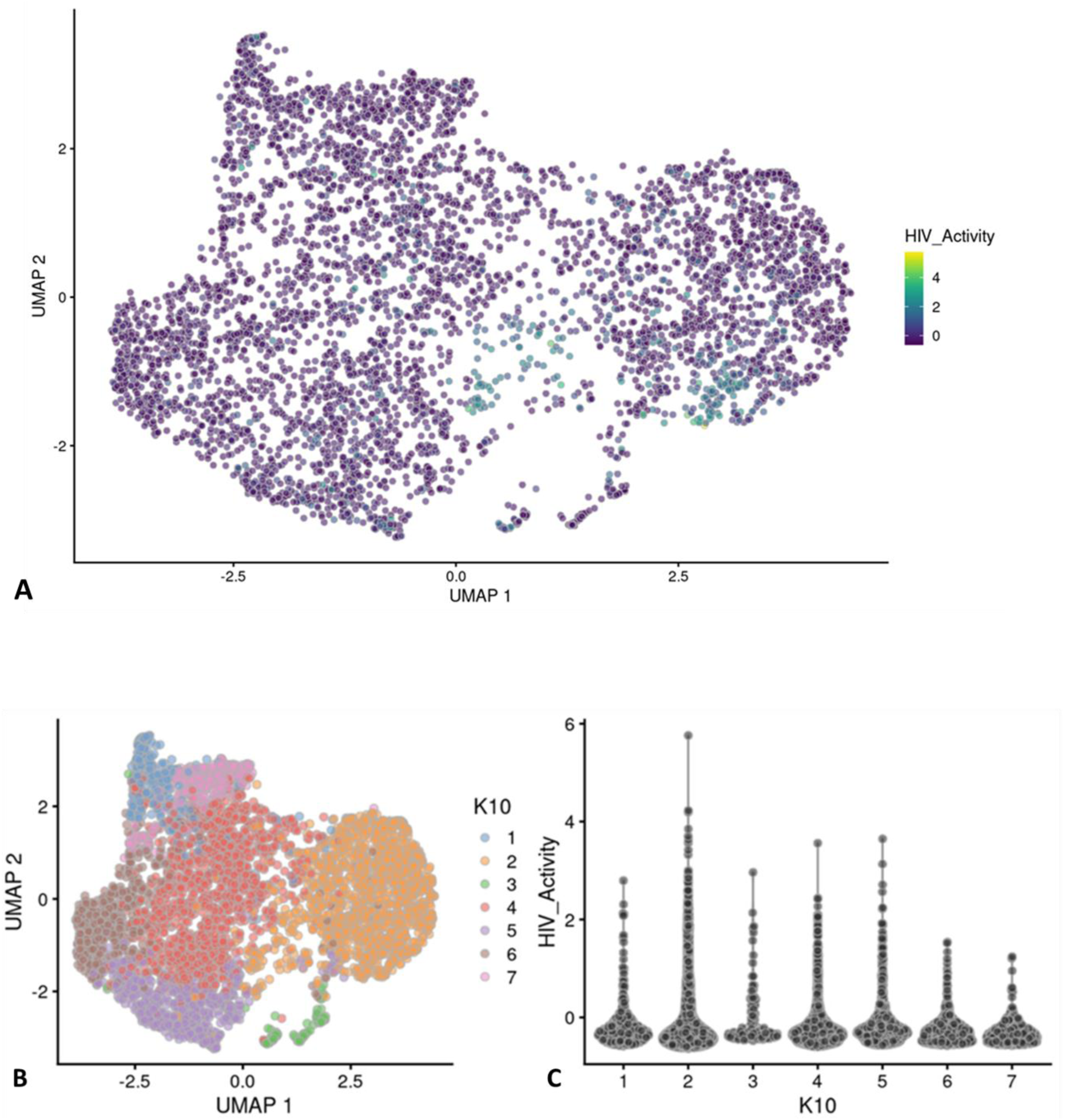
UMAP analysis of integrase inhibitor treated DHIV3-mCherry infected THP-1 cells. The experiment performed as shown in Figure 6 B, with 25nM MK-2048 added at the time of DHIV3 addition. Data were analyzed identically to data shown in Figure 2. Panel **A)** Feature plot showing the distribution of cells containing HIV-1 transcript, generated as described. Panel **B)** K10 unsupervised clustering generated 7 clusters (Scran’s buildSSNGraph using the PCA as input). HIV-1 transcripts were distributed equally throughout all of them. No cluster corresponding to the “Provirus” cluster detected in Figure 4 was detected, regardless of K value used (see Fig. S-6). These data agree with the concept that integrase inhibitors selectively target and reduce the number of Provirus cluster cells.

### Hallmark and REACTOME analyses indicate mitogenic associated pathways are up-regulated in Provirus cluster cells whereas viral restriction and interferon associated pathways are upregulated in PIC cluster cells

Differential gene expression (DGE) in Provirus versus PIC/Bystander cluster transcriptomes was analyzed using GSEA with Hallmark or REACTOME gene lists (Appendix I), the pairwise TTests function from Scran was used to determine the statistically significant differentially expressed genes between groups. Table I presents the statistically significant Hallmark results. By comparing the DGE between the Provirus and PIC/Bystander cell clusters using Hallmark and REACTOME tools, it was clear that Provirus cluster cells were differentiated from PIC/Bystander cluster cells in several key ways. In general, the transcriptomes of cells in the Provirus cluster were characterized by gene transcripts associated with cell replication, whereas the transcriptomes of cells in the PIC/Bystander cluster were characterized by pathways associated with immune-response and interferon signaling. The analyses clearly showed upregulation of cell replication, oxidative phosphorylation, protein synthesis and E2F family targeted pathways in the Provirus cluster cells. On the other hand, NFkB, AP1, interferon responsive and immune response pathways relatively upregulated in the PIC/Bystander cluster cell transcriptomes. Intuitively, this makes sense, but it runs contrary to established literature that associates E2F transcription factors with decreased viral production [24, 25]. This finding would not be obvious without the use of single cell analysis. In the absence of single cell analysis the Provirus cluster’s differentially expressed gene transcripts would have been swamped out by the 90% of mRNA obtained from PIC and Bystander cells.

**Table I.**
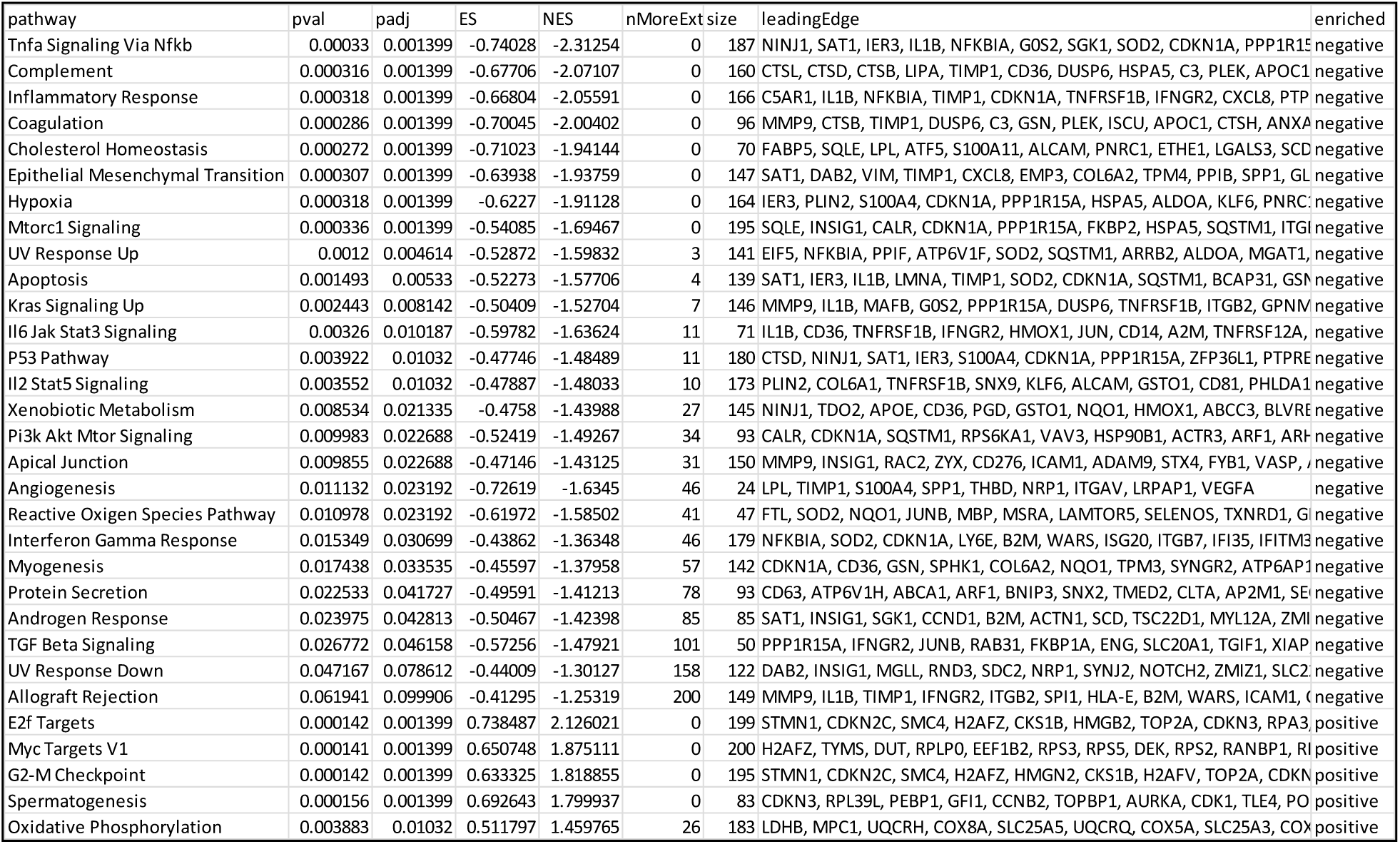
Hallmark analysis of gene pathways up or down-regulated detected in Provirus versus PIC/Bystander cluster cells (respectively). GSEA Hallmark analysis (fgsea R package) of metabolic pathways negatively or positively regulated (p<0.1) in the Provirus cluster transcriptome versus the PIC/Bystander cluster transcriptome. Pathways up-regulated in Provirus cells include E2F, Myc trargets, G2-M checkpoint, spermatogenesis, and oxidative phosphorylation. Down-regulated pathways identified are more numerous, but notably included TNFα signaling via NFkB, inflammatory response genes, apoptosis, and interferon γ response. The pairwiseTTests function from Scran was used determine the significance of genes between groups. The significant DGE subsets were used for all comparisons.

Using the UMAP Feature plots shown in Figure 9, we visualized the distribution of cells containing some of the most statistically significant differentially expressed transcripts in our Provirus cell transcriptomes versus the PIC/Bystander cell transcriptomes. The GSEA lists of the differentially expressed genes are provided in supplementary Appendix I. This leads us to hypothesize that the cell’s transcriptome background is what determines if virus transcript will lead to virus protein production, not the differential transcription of HIV-1 genes. This hypothesis was supported by Transcription Factor Targeting and Western blot analyses (below).

**Figure 9.**
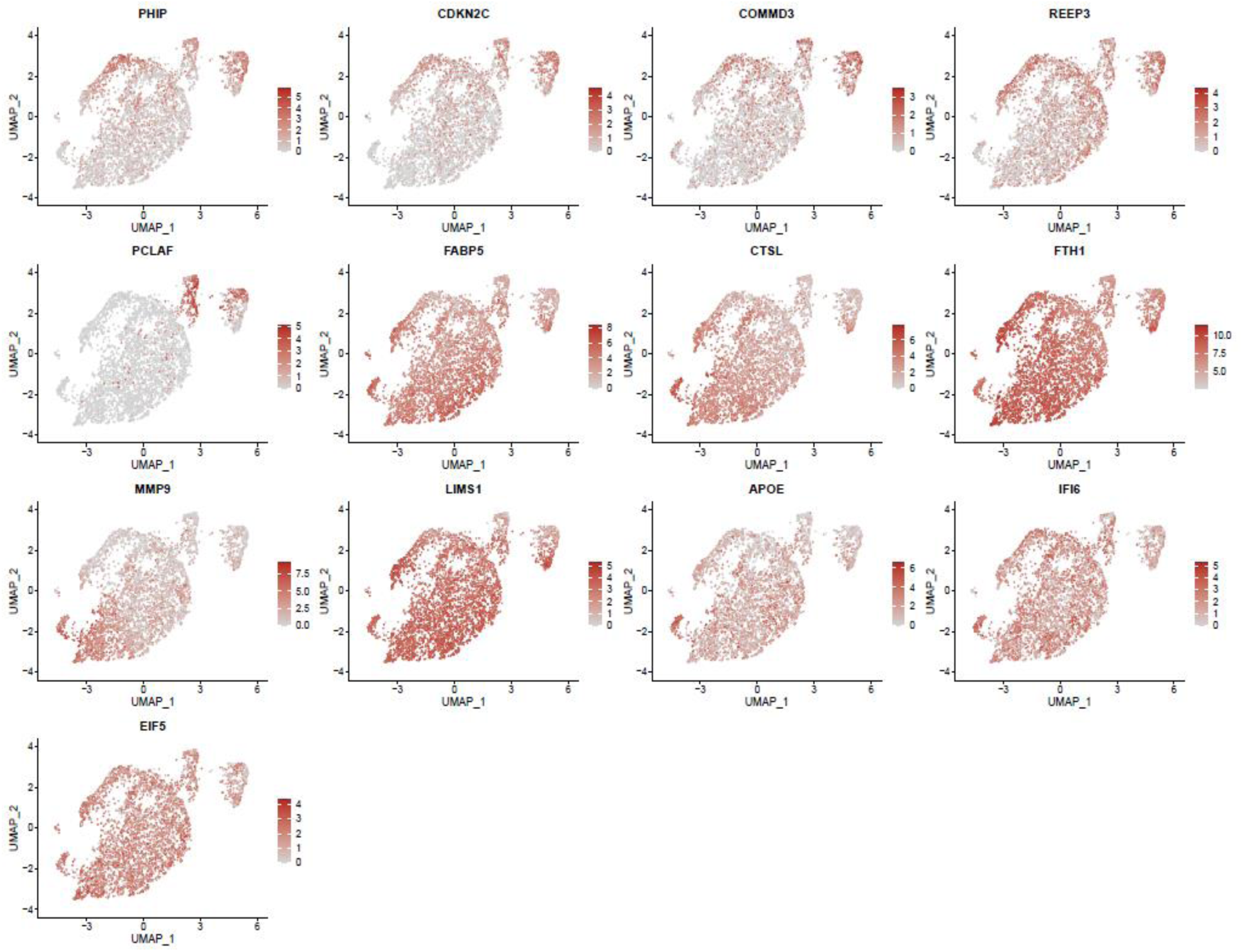
Distribution of gene transcripts exhibiting high levels of differential expression between Provirus and PIC/Bystander clusters. Feature plot showing the distribution of cells from Figure 2, containing transcripts of 10 of the 20 most highly differentially expressed transcripts in Provirus versus PIC/Bystander GSEA data sets. APOE, IFI6, and EIF5 were also included because they were highly expressed in the PIC/Bystander cluster. As above, these UMAP projections were made with Seurat’s FeaturePlot function. They are colored by expression of individual genes (normalized log2 values). Highly expressed genes in Provirus cluster cells include PHIP (Pleckstrin Homology Domain Interacting Protein), CDKN2C (Cyclin Dependent Kinase Inhibitor 2C), COMMD3 (COMM Domain Containing Protein 3), REEP3 (Receptor Accessory Protein 3), and PCLAF (PCNA Clamp Associated Factor). Highly expressed transcripts detected in the PIC/Bystander cell transcriptome include FABP5 (Fatty Acid Binding Protein 5), CTSL (Cathepsin L), FTH1 (Ferritin Heavy Chain 1), MMP9 (Matrix Metallopeptidase 9), LIMS1 (LIM Zinc Finger Domain Containing 1), APOE (Apolipoprotein E), IFI6 (Interferon Alpha Inducible Protein 6), and EIF5 (Eukaryotic Translation Initiation Factor 5).

### PIC and Provirus Cells express all DHIV3 genes at equivalent levels, although higher numbers of PIC cells detectably express some transcripts

We were interested to know if early HIV-1 gene transcripts (tat, nef, or rev) predominated in PIC cell transcriptomes, versus later transcripts in the Provirus cells. We used Feature plots to determine the distribution of selected HIV gene transcripts expressed in individual cells. It was found that cells expressing the respective early or later gene transcripts were distributed throughout the image (Fig. 10 A).

**Figure 10.**
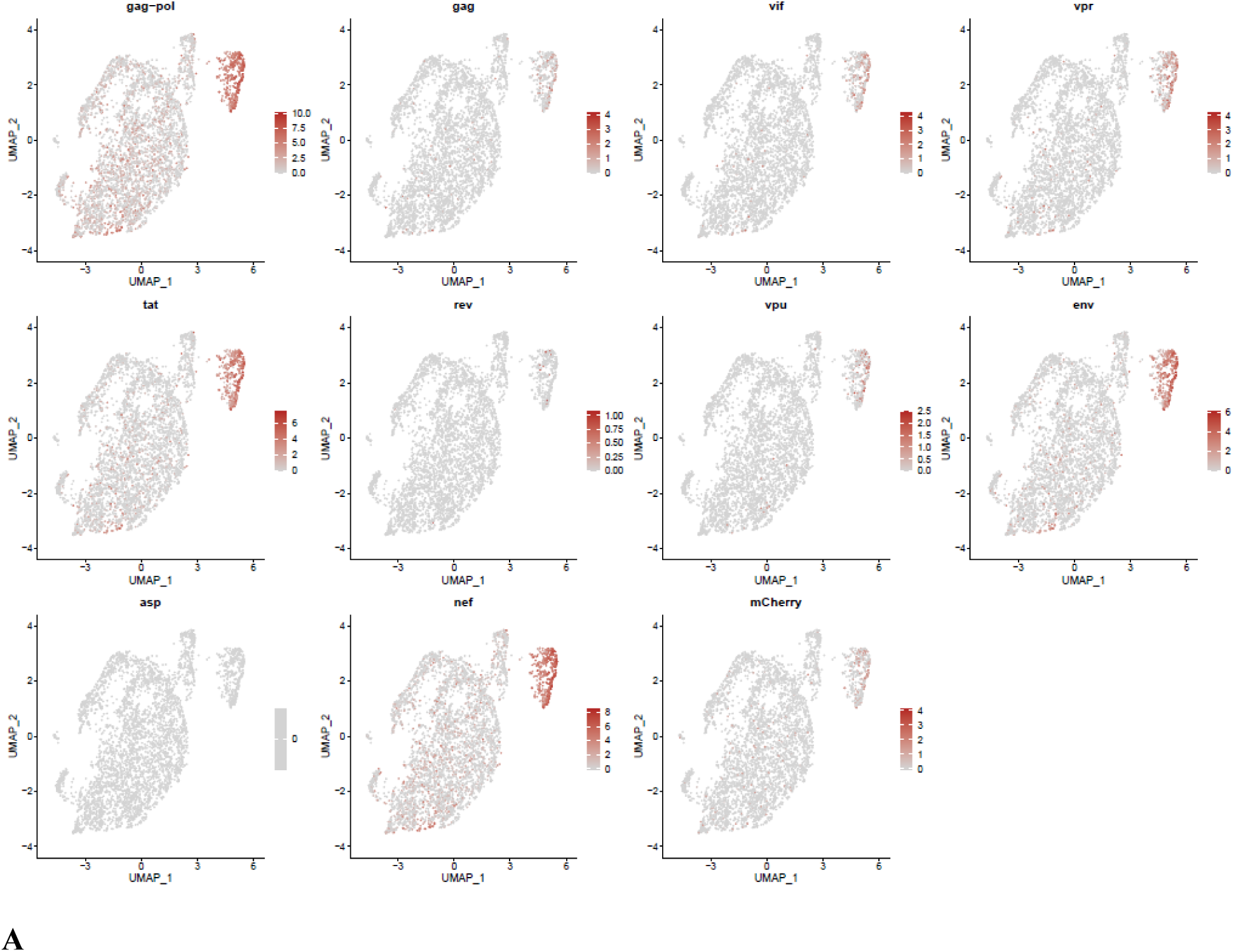
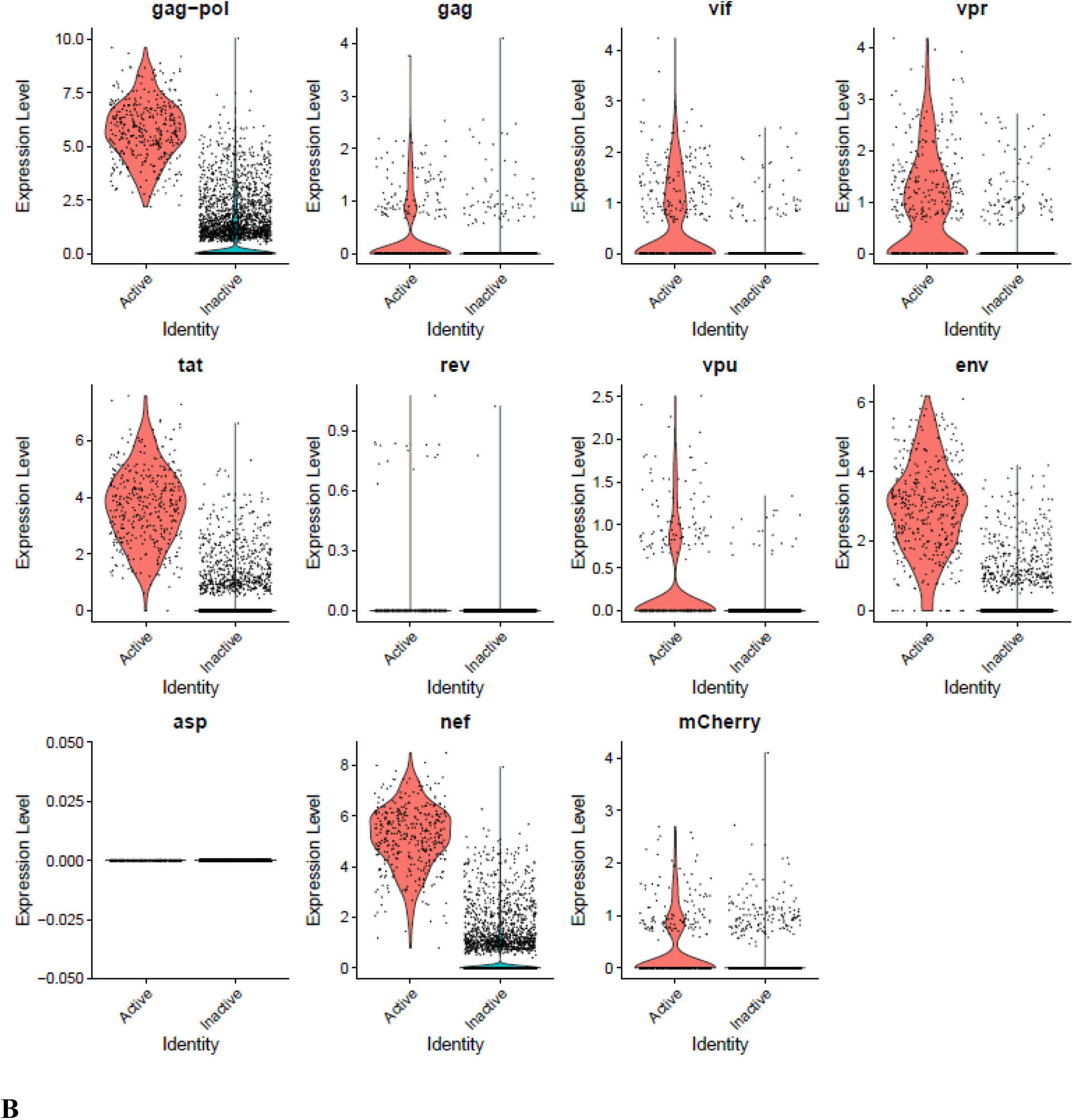
The Distribution of HIV-1 transcripts throughout Provirus and PIC/Bystander clusters. Panel **A**) Feature plot showing the distribution of cells from UMAP in Figure 2 that contain detectable DHIV3-mCherry transcripts. As described above, these UMAP projections were made with Seurat’s FeaturePlot function. They are colored by expression of individual genes (UMAP projection colored by walktrap, normalized log2 values). ASP is a negative control, bacterial gene transcript sequence. **B**) Violin plots of DHIV3-mCherry transcript/cell in cells from the Provirus and PIC clusters showing transcript level and cell number. The provirus cluster contained transcriptomes of 371 cells, the number of PIC cells in the PIC/Bystander cluster was 569 cells, thus the Provirus/PIC cell number ratio was 0.65. The plots were made with Seurat’s VlnPot function. They show normalized log2 transcript levels. Two patterns of transcript distribution are evident. The first pattern is seen with gag-pol, tat, env, and nef, in which relatively high numbers of cells in the PIC/Bystander cluster express the transcripts, with the transcripts being detected in fewer numbers of Provirus cluster cells. The second pattern is seen with gag, vif, vpr, rev, vpu, and mCherry, in which relatively equal absolute numbers of cells in the Provirus and PIC/Bystander clusters are detected with the transcript sequences, remembering that there are more PIC cells than Provirus cells. The relative transcript loads per PIC cell versus the Provirus cells overlap. Negative control sequence (asp) shows no distribution.

We then used violin plots to compare the load of individual DHIV3 gene transcripts in Provirus cluster cells to PIC cells (Fig. 10 B). The Provirus cluster (from Fig. 4) contained transcriptomes of 372 cells, the PIC/Bystander cluster contained 569 cells, thus the ratio of Provirus to PIC cells was ∼0.65. This visualization was much more informative and yielded a more nuanced understanding (Fig. 10 B). Some transcripts, such as gag-pol, tat, env and nef were detected in far more cells in the PIC cluster, albeit often at the lower level of detectable loads per cell. In contrast, gag, vif, vpr, rev, vpu and mCherry, were clearly detectable, but in lower numbers of Provirus and PIC cells. Cells containing these transcripts were comparably prevalent in the two groups. All transcripts were easily detectable in both Provirus cells and PIC cells. Furthermore, there is a clear overlap in the levels of HIV-1 gene transcripts detected in Provirus and PIC cells.

During transcription of pro-virus, HIV-1 does not produce all transcripts in equal number or at the same time [37, 38]. The specific processed gene transcripts produced initially in viral replication differ from transcripts produced later on. Furthermore, 10X Genomics scRNA-seq library production is known for significant numbers of dropouts, and cDNA copying of various gene transcripts during library construction varies in efficiency [15, 39, 40]. Furthermore, in using poly-T primers in the cDNA library construction, the 10X process introduces a 3’ bias toward the detection of given sequences in a transcript [15]. Thus, it is not possible to make quantitative comparisons between the different transcripts using this approach. Nevertheless, the overarching take-away from this single cell analysis is that cells making fully spliced transcripts such as nef, tat, and nev are equally prevalent with cells making gag-pol transcripts (Fig. 10). Furthermore, there appears to be two patterns of transcription from PIC. One pattern, observed with gag-pol, tat, env, and nef, is characterized by gene transcripts being more frequently detectable in PIC cells than Provirus cells. The other pattern, observed with gag, vif, vpr, rev, vpu, and mCherry, suggests relative equal frequency of transcription in Provirus and PIC cells.

### Biological repeat experiments confirm observations

To confirm the conclusions obtained from the analyses presented above, an independent biological repeat experiment was conducted. The repeat experiments captured over 4,500 cells, with an average greater than 3,000 genes per transcriptome. The control cell cultures again yielded a slightly larger number of cells captured with more than 3,000 genes transcripts per cell. The biological repeat experiments, (HIVreplicate2 and WT2) were also conducted with parallel cultures that were analyzed by flow cytometry. Flow cytometry indicated 2.6 percent mCherry positive cells in the HIVreplicate2 experiment. Figure S-8 shows a UMAP analysis of the biological repeat experiments HIVreplicate1 and HIVreplicate2. Figure S-9 shows unsupervised clustering of HIVreplicate2 using K nearest neighbor values from 10 to 130. Figure S-10 shows distribution of respective DHIV3-mCherry transcripts throughout HIVreplicate2 UMAP.

Several comparisons were made to confirm that transcriptomes from Provirus and PIC/Bystander clusters in the repeat experiments were identical. In the first comparison, differentially expressed genes (positive or negative) were identified between the Provirus and PIC/Bystander clusters in the respective biological repeats. The log2 fold changes from these gene sets were then compared to test if the differences between the transcriptomes of Provirus and PIC/Bystander cells in the repeat experiments was consistent (Fig. S-11). Because there almost 4 times as many Provirus cluster cells in the HIVreplicate1 experiment, there were more significantly differentially expressed genes in the HIVreplicate1 case, compared to the HIVreplicate2 case. Nevertheless, two the gene sets were positively correlated. Correspondingly, GSEA with Hallmark and REACTOME gene sets identified many of the same pathways as differentially regulated in the two biological repeats (Appendix I).

To obtain additional statistical certainty for our conclusion that results obtained for HIVreplicate1 were repeated in HIVreplicate2, we compared log2 fold change values from the Provirus or PIC/Bystander clusters in HIV infected cells to the log2 fold change values from the Control (WT) PMA-treated THP-1 cultures. This 8-way comparison (shown in Fig. 11) provided statistical certainty that DGE sets from the Provirus cluster and PIC/Bystander cluster gene sets from the biological replicates were not different. The replicate Provirus and PIC/Bystander gene sets have a generally strong concordance between themselves and there is a modest to strong non-zero mean trend in logFC among genes that changed in at least one of the contrasts among replicates (FDR 5%). In every comparison, a significant positive correlation was obtained from the commonly detected, significantly differentially expressed genes of Provirus or PIC/Bystander clusters in the two biological repeats when compared to the Control samples. The weakest correlations were observed in comparing PIC/Bystander to Control cell DEGs, especially Control experiment 2 (probably because there is more commonality between genes detected in PIC/Bystander cells and Control cells than there is between Provirus cluster cells and Control cells), but still the data between HIVreplicate1 and HIVreplicate2 were concordant. Therefore, the transcriptome data obtained from the 2 biological repeat experiments were not different. In other words, statistically identical representative transcriptomes for Provirus and PIC/Bystander clusters were obtained in independent biological repeat experiments.

**Figure 11.**
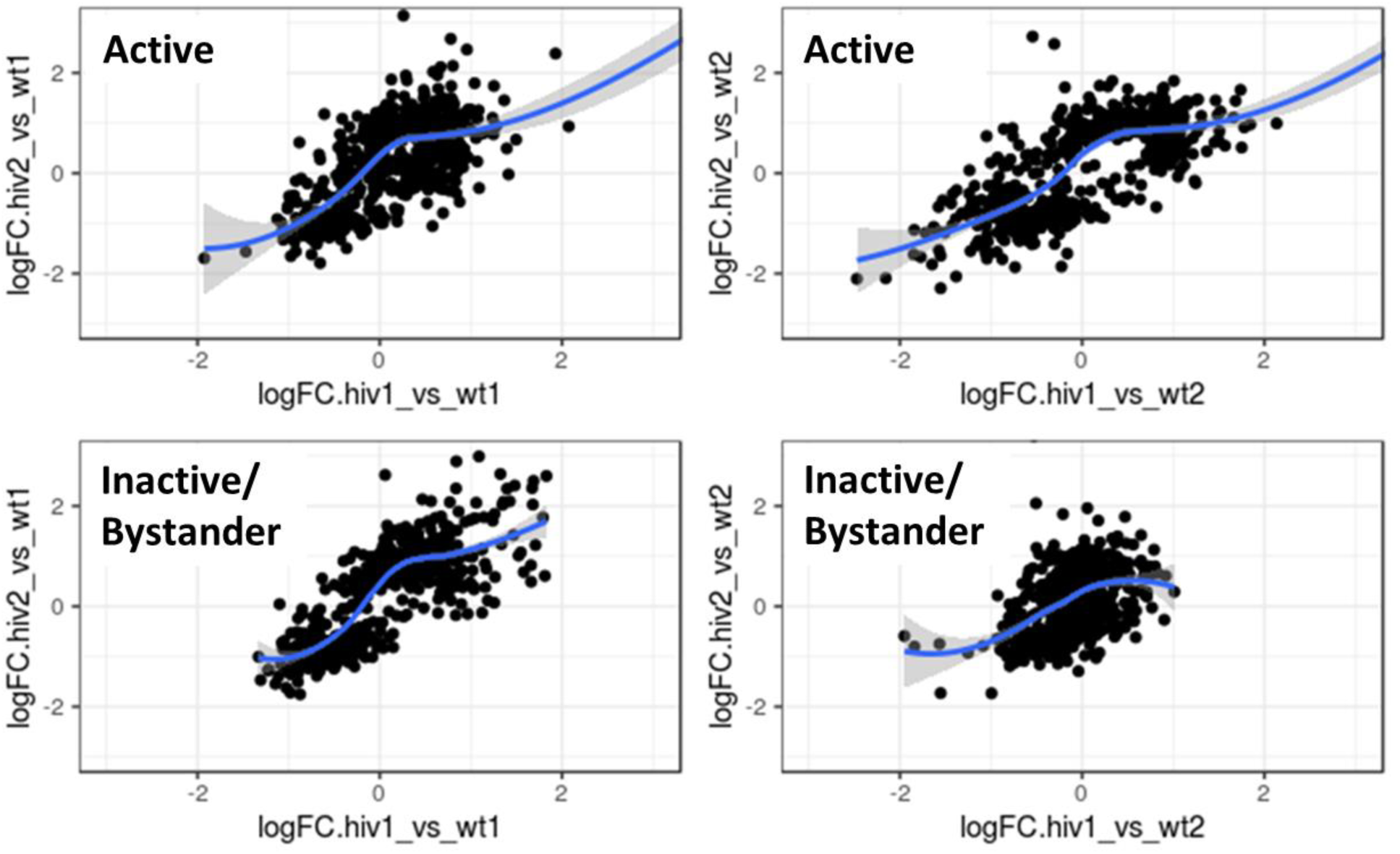
Differential gene expression comparison of Provirus and PIC/Bystander cluster gene transcripts versus 2 independent biological repeat control (wt) experiments. In every comparison, a significant positive correlation was obtained from the common detected differentially expressed genes of Provirus or PIC/Bystander clusters in the two biological repeats when compared to the Control samples. Consistent positive correlation in this 8-way comparison confirmed statistical identity between biological repeat experiments. The trend line in the plot is the result of the function: stats::loess (R Package Documentation) [47], using default parameters. The fitted curves are shown with 95% confidence band.

As was the case for HIVrepeat1, a clear Provirus cluster was detectable in the HIVrepeat2 (Fig. 12, S-8). However, because the level of Provirus infection in HIVrepeat2 was lower than in HIVrepeat1, the frequency of DHIV3 transcript detection in the Provirus cluster cells was proportionately lower, while the absolute number of detectable PIC cells was relatively higher. As was observed for HIVrepeat1, PIC cells with detectable HIV-1 transcripts, were randomly distributed throughout the Bystander cluster. Unsupervised clustering (Fig. 12, S-9) generated 10 clusters at a K nearest neighbor values of 10. Clusters 1, 2 and 4-10 contained most of the cells in the semi supervised PIC/Bystander from Figure 12. Cluster 3 contained 135 of the 227 cells in the semi supervised Provirus cluster (circled in red, Fig. 12), and was compared to the combined transcriptomes of the remaining clusters to generate the violin plots in Figure 13. The patterns of transcription observed in HIVrepeat1 were confirmed in this experiment. The transcripts of gag, vif, vpr, vpu, and mCherry were detectable in PIC cells at frequencies and levels of expression similar to those observed in the Provirus cells. In contrast, even correcting for the Provirus/PIC ratio of 0.17, the transcripts of gag-pol, tat, env, and nef were again detectable in proportionally higher number of PIC cells. Again, there was clear overlap in the loads of HIV-1 gene transcripts detectable in Provirus and PIC cells. No cells expressing rev were detected in the Provirus cluster in this repeat, due either to the low efficiency of detecting this transcript in the 10X system, or because rev is expressed only at low levels in relatively few cells, or both.

**Figure. 12.**
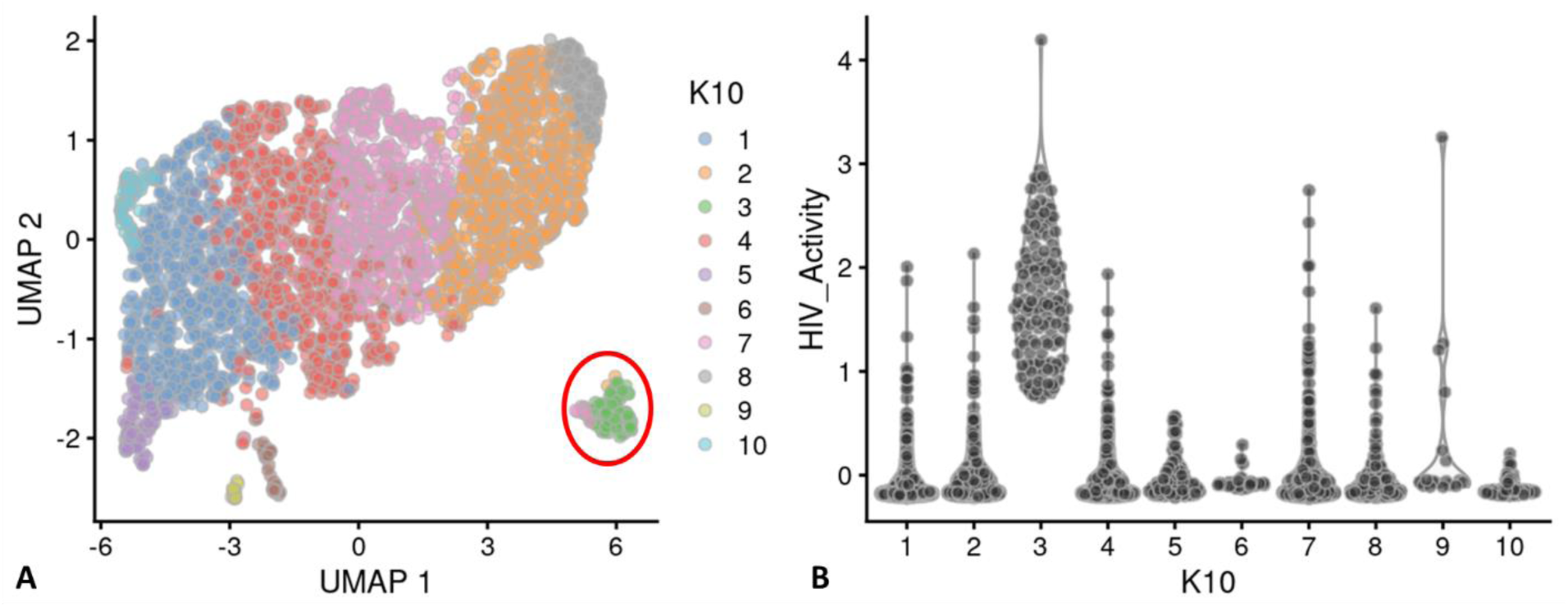
Unsupervised clustering of HIVrepeat2. Panel **A**) shows unsupervised clustering obtained at K equals 10. Panel **B**) Violin plot of HIV-1 transcripts/cell in the 10 clusters identified at K10 (Scran’s buildSSNGraph using the PCA as input). PIC cells with detectable HIV-1 transcripts, were distributed throughout clusters 1, 2 and 4-10. Cluster 3 contained 135 of the 227 cells in the semi supervised Provirus cluster (circled in red).

**Figure 13.**
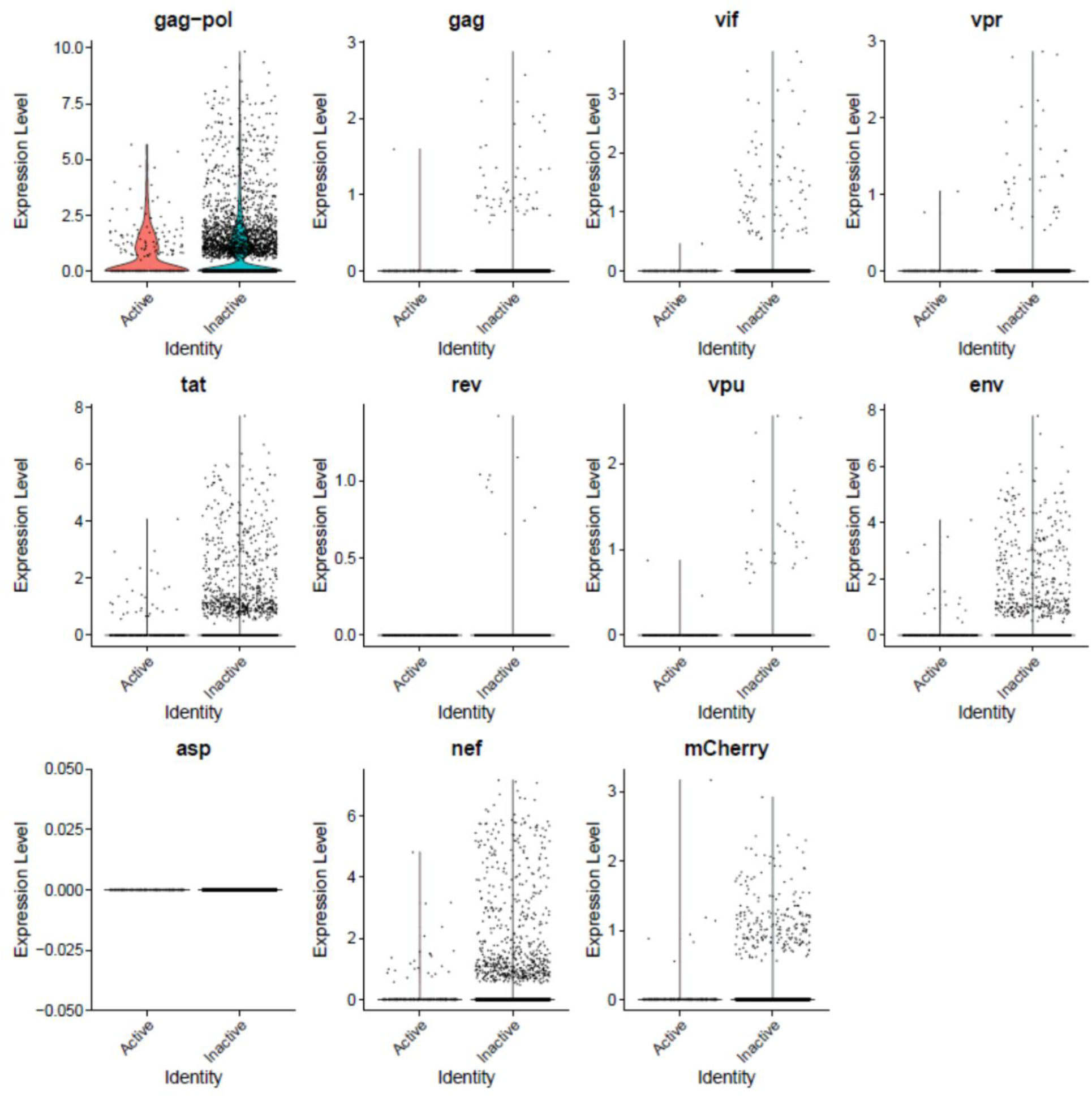
The Distribution of HIV-1 transcripts throughout Provirus and PIC/Bystander clusters in HIVrepeat2. Violin plots of DHIV3-mCherry transcript/cell in cells from the Provirus and PIC clusters showing transcript level and cell number. As described above, these were made with Seurat’s VlnPot function. They show normalized log2 transcript levels. The two patterns of transcript distribution observed in HIVrepeat1 are evident. The first pattern is seen with gag-pol, tat, env, and nef, in which high numbers of cells in the PIC/Bystander cluster detecably express the transcripts. The second pattern is seen with gag, vif, vpr, vpu, and mCherry, in which fewer Provirus or PIC cluster cells are detected expressing the transcripts, but those cells expressing the transcripts are doing so at slightly higher average levels of transcripts per cell. It is difficult to compare transcript loads in the Provirus cluster cells to the results in HIVrepeat1 (Fig. 10) due to the lower number of Provirus cells detected in this HIVrepeat2 experiment. In this experiment, the ratio of Provirus cells to PIC cells was 0.17. Nevertheless, the relative patterns observed in HIVrepeat1 are observed here. Following Seurat QC, no Provirus cells expressing rev were detected. Negative control sequence (asp) shows no distribution.

### Psupertime analysis indicates progression of cluster transcriptomes from Control to PIC/Bystander to Provirus

To understand the transcriptome transitions needed to move from unexposed and uninfected “Control” cells to PIC/Bystander cells, and on to Provirus cluster cells, we performed a psupertime analysis [41–43] of the respective cell cluster transcriptomes. Psupertime is a supervised pseudotime [41] technique. It explicitly uses sequential condition labels as input. Psupertime is based on penalized ordinal logistic regression that places the cells in the ordering specified by the sequence of labels. This allows for targeted characterization of processes in single cell RNA-seq data.

One thousand cells were randomly selected from each transcriptome cluster (Control, PIC/Bystander and Provirus) and their transcriptomes combined for psupertime analysis. Imposition of Cluster identity yielded the image shown in Figure 14. The psupertime-type analysis showed closer similarity between Control and PIC/Bystander transcriptomes than between Control and Provirus transcriptomes, and closer similarity between PIC/Bystander and Provirus transcriptomes than between Control and Provirus. The GSEA list of the DGEs that contributed to this faux-progression from Control to PIC/Bystander to Provirus are presented in Appendix II.

**Figure 14.**
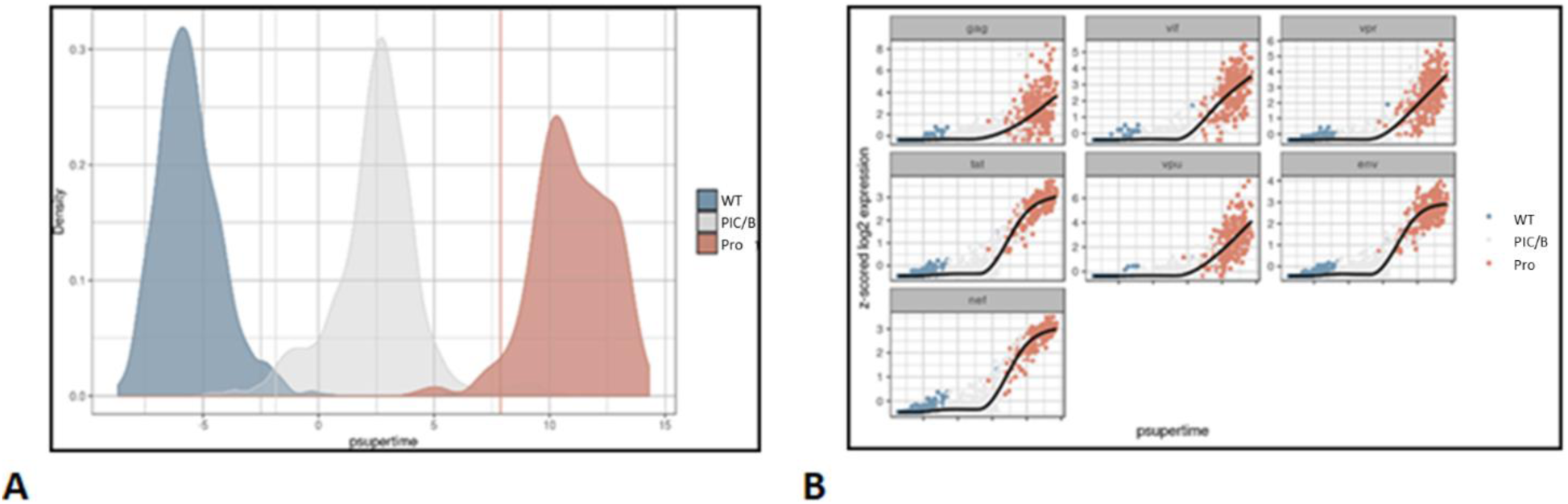
Psupertime analysis of Control, PIC/Bystander, and Provirus cell transcriptomes. Psupertime analysis is a supervised pseudotime approach that explicitly uses the sequential labels as input. It uses a regression-based model that acknowledges the cell labels to identify genes relevant to the process. Panel **A**) one thousand Control (WT), PIC/Bystander (PIC/B), and Provirus (Pro) cell transcriptomes were randomly selected and analyzed. Imposition of identity revealed a pseudo-evolution of Control to PIC/Bystander to Provirus cell transcriptomes. Panel **B**) distribution of HIV-1 transcripts through these clusters agrees with results shown in Figure 5, showing no bias toward early or later gene transcripts.

When we examined the expression of DHIV3 transcripts through the psupertime progression, the analysis showed no obvious preference for early gene transcription in cells belonging to the PIC/Bystander versus the Provirus clusters (Fig. 14B).

When questioning which transcription factors were regulating the transcriptome transitions, we searched the contributory psupertime DGE transcripts for transcription factors. Many transcription factors that differed in expression in the contrasted transcriptomes were identified, Appendix II. However, this yielded a complicated picture and did not clarify which transcription factors might be most important for controlling the transcriptome transitions from Control to PIC/Bystander to Provirus clusters. However, because the activity of most transcription factors is regulated by activation of proteins already present within the cell, and not at the transcription level, we speculated that Transcription Factor Targeting analysis might be more informative as to which transcription factors were key to cluster transitions.

### E2F, NF-kB and AP1 control phenotype transitions between PIC/Bystander cells and Provirus cells

We used Transcription Factor Targeting analysis to identify transcription factors that controlled DGEs in our Control, PIC/Bystander and provirus clusters. This analysis was consistent with the aforementioned Hallmark and REACTOME analyses. The E2F family of transcription factors predominate in regulating the Provirus cluster DGEs (Table II). Twenty out of the 29 possible promoter-associated transcription factor interactions positively associated with the transition from the PIC/Bystander to the Provirus transcriptome identified with the E2F transcription factor family (Table 2). Thus, E2F is clearly associated with pathways that determine the phenotype of cells in the Provirus cluster. Conversely, 19 possible promoter-associated transcription factor interactions were negatively associated with DGEs reflecting the transition from PIC/Bystander to Provirus cells. Of these 19 possible promoter complexes, 8 different promoter interactions were identified to be associated with NFkB and AP1 transcription (Table 2) suggesting that downregulation of NFkB and AP1 also plays a key role in shaping the Provirus cluster cell transcriptome.

**Table II.**
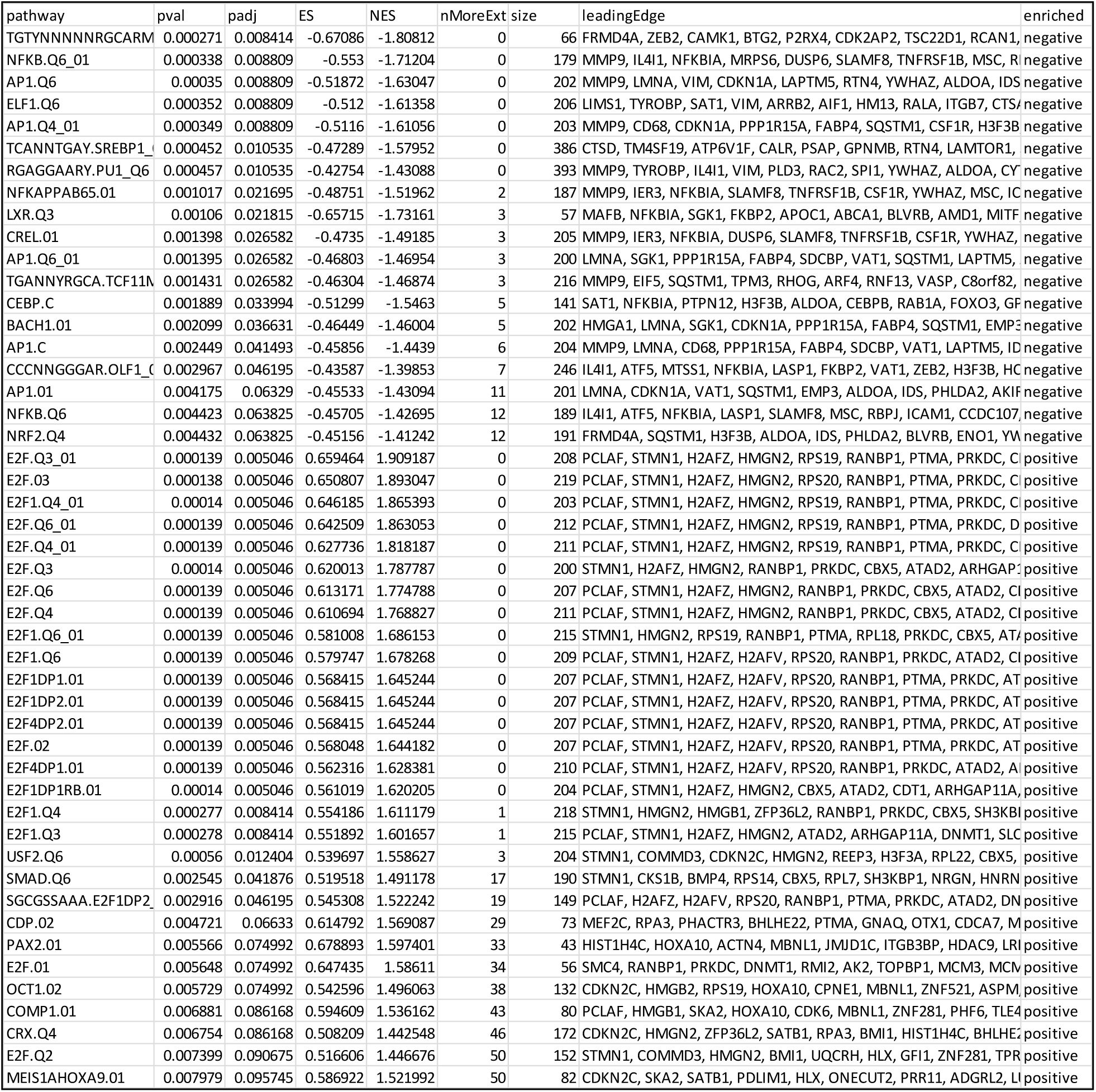
Transcription Factor Targeting analysis of DGE contrasting PIC/Bystander and Provirus cells. TFT analysis (GSEA with the fgsea R package and the C3 collection from msig) suggests that at least 3 transcription factor families control the transition from PIC/Bystander transcriptomes to Provirus cluster transcriptomes. These are E2F, NFkB, and AP1 family promoter proteins. In particular, increased E2F regulated transcription appears to correspond with the transition to the production of viral proteins. The pseudo-transition from Control to PIC/Bystander is characterized by down-regulation of E2F family regulated transcripts and up-regulation of NFkB and AP1 regulated transcripts Appendix III. In comparing Provirus to PIC/Bystander transcriptomes, E2F family promoted transcripts are up-regulated, while NFkB and AP1 transcription products are down-regulated. Comparing Provirus to Control transcriptomes shows that overall Provirus cells have increased E2F regulated transcripts and decreased NFkB transcripts (with no significant change detected in AP1 regulation).

AP-1 and NFkB appear to play roles in maintenance of the PIC/Bystander cell transcriptome as well. Correspondingly, Transcription Factor Targeting identified differences between Control and PIC/Bystander cell transcriptomes. Simple exposure of activated THP-1 cells to DHIV3 was sufficient to decrease E2F signaling in PIC/Bystander cells, compared to Control cells, and to increase AP-1 and NFkB signaling (Appendix III). We presume this effect on the PIC/Bystander cells is through PAMP and/or interferon signaling. Again, these results are consistent with the results obtained with Hallmark and REACTOME analysis of the DGEs.

### Western Blot analysis of Retinoblastoma protein phosphorylation from Control, PIC/Bystander and Provirus cells

To confirm a role for E2F and NFkB in regulating the transcriptomes of Provirus and PIC/Bystander clusters (respectively), we sorted Provirus (mCherry positive cells) from PIC/Bystander cells using the FACS Canto. Rb phosphorylation is associated with activation of E2F promoter family transcription. We hypothesized Provirus cluster cells would exhibit retinoblastoma (Rb) phosphorylation consistent with E2F activation [27, 42]. We used anti-T821 Phospho-Rb antibody. Phosphorylation of Rb at threonine-821 (T821) blocks pocket protein binding, including E2F family proteins, and activates E2F family promoter gene transcription [44]. Isolated Provirus cells had the greatest phosphorylation of Rb (pRb), compared to Control and PIC/Bystander cells (Fig. 15). Interestingly, phosphorylation of Rb in PIC/Bystander cells was actually lower than that detected in Control cells (Fig. 15). We also probed these blots for mCherry protein as was shown in Figure 7. In this case, with cells being sorted before protein preparation, mCherry protein detection was detectable and only in the Provirus cluster cells.

**Figure 15.**
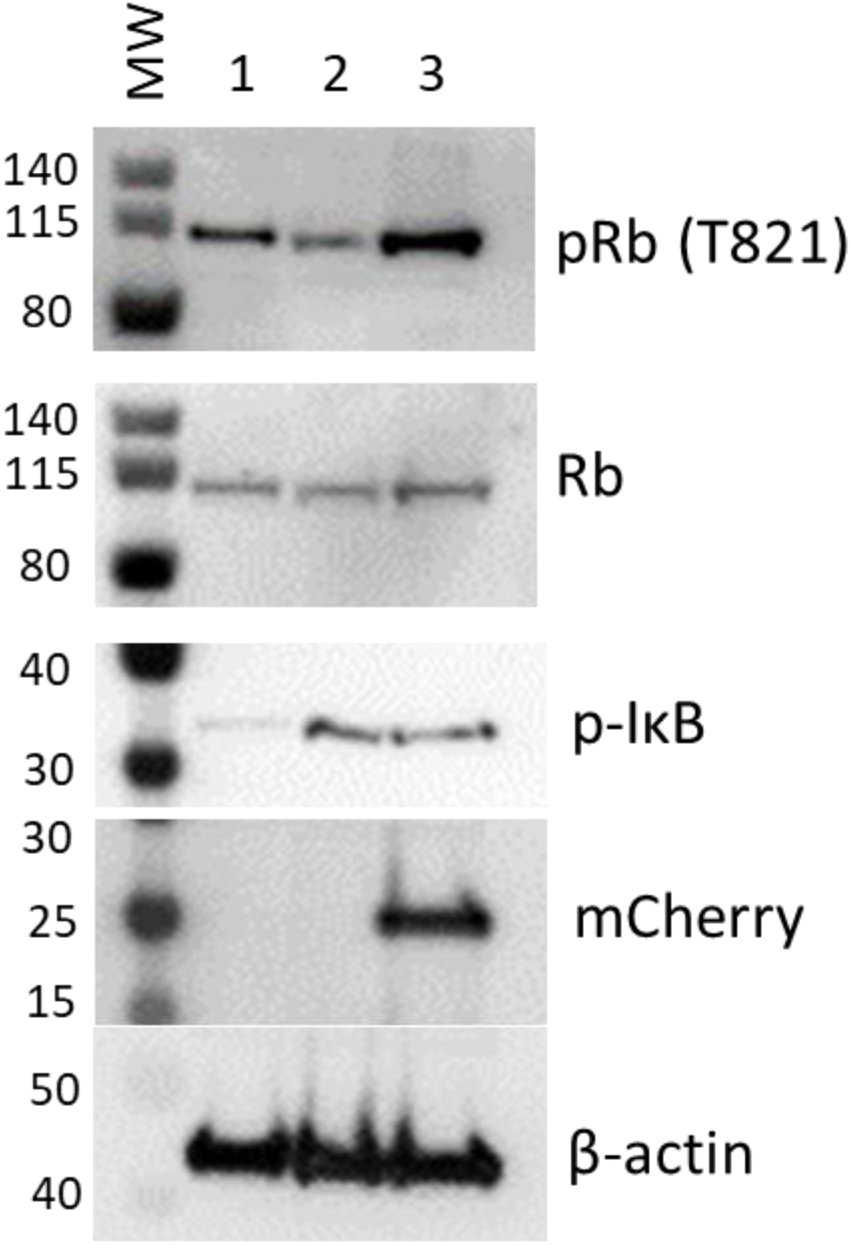
Western blot analysis for phospho- Rb or IkB in protein from mCherry negative versus mCherry positive cells. Cells infected with DHIV3-mCherry were purified by FACS sorting based on their expression of mCherry fluorescence. Lane 1) Protein from Control cells; Lane 2) Protein from PIC/Bystander cells; Lane 3) Protein from Provirus cells. Phospho- Rb (Phospho-T821 Rb antibody) was used to quantify Rb pocket phosphorylation, anti-Rb control antibody was used to quantify Rb protein levels relative to actin (visualized with beta- actin antibody). mCherry protein confirmed with anti-mCherry antibody used in Figure 7. PIC/Bystander cells show the lowest level of Rb phosphorylation, Provirus show the highest, in close agreement with Transcription Factor Targeting results. Panel B, Lane 1) Protein from Control cells; Lane 2) Protein from PIC/Bystander cells; Lane 3) Protein from Provirus cells. Phospho-IkB S32 antibody was used to quantify activated IkB. Control cells show the lowest level of IkB phosphorylation, no difference was detectable between Provirus and PIC Cluster cells.

These findings are consistent with the Transcription Factor Targeting analysis, which showed that E2F promoter family transcription was predominant in Provirus cluster cells. It also agreed with the Transcription Factor Targeting analysis in that PIC/Bystander cells exhibited reduced Rb phosphorylation, and presumably E2F driven transcription, when compared to either Provirus cluster of Control cells. We did not find a difference in NFkB deactivation in the Provirus cells as would be implied by phospo-IkB S32, (Fig. 15), and hypothesize that other mechanisms must account for the relative decrease in NFkB driven transcription in Provirus cluster cells.

### Cells transcribing from provirus are more likely to produce viral proteins upon second infection than PIC or Bystander cells

If Provirus cluster cells have already committed to the production of virus, represented by the switch of their background transcriptome to favor E2F transcription factor interactions, we hypothesized that they should be more efficient at producing virus upon second infection. We sequentially infected of activated THP-1 cells with DHIV3-mCherry followed at 24 hr by an infection with DHIV3-GFP (DHIV3-mCherry infection at 0 hr and DHIV3-GFP infection at 24 hr). We found a higher percentage of cells positive for mCherry and GFP after 48 hr compared to GFP alone (Fig. 16). At his time point, which was 24 hours after DHIV3-GFP infection, about half the mCherry positive cells were also GFP positive. Whereas, less than one quarter of the mCherry negative cells were expressing GFP protein. This trend continued out to 72 hours post DHIV3-mCherry infection, 48 hours after DHIV3-GFP addition, where about 60% of the mCherry cells were also GFP positive, compared to about 40% GFP positive in mCherry negative cells. In repeat experiments and in experiments using primary macrophage and T-cell cultures (Fig. S-12) the results were the same. Provirus cluster, mCherry positive cells, were 2X to 5X more likely to make DHIV3-GFP protein upon second infection than PIC/Bystander cluster cells. Thus, cells already committed making virus were more likely to make virus from a second infection than PIC/Bystander cells on first infection.

**Figure 16.**
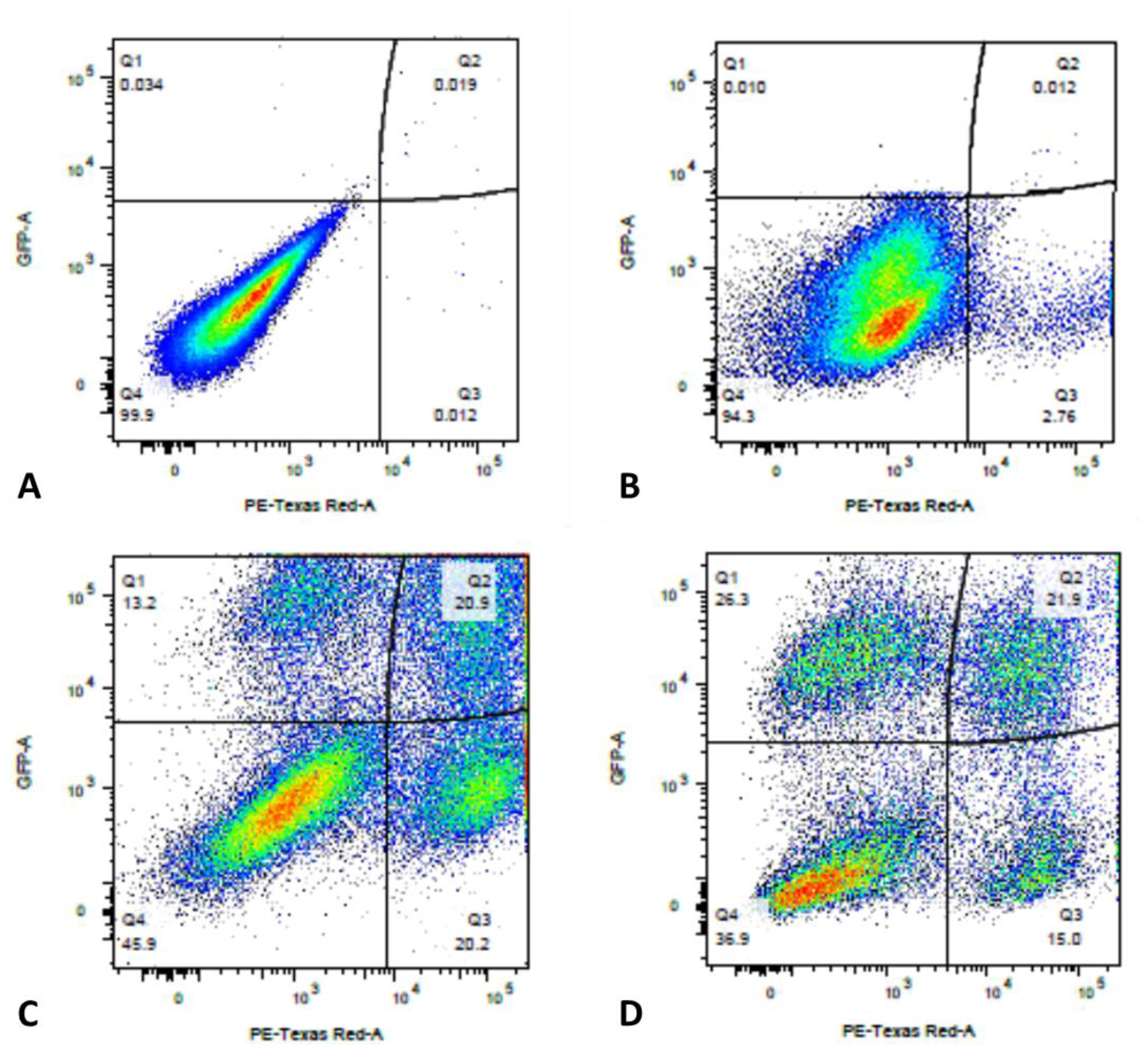
Sequential infection of THP-1 cells with DHIV3-mCherry followed 24 hrs later with GFP DHIV3. Abscissa, mCherry signal, Ordinate, GFP signal. Provirus cluster, mCherry positive, cells were 2 to 5 times more likely to make HIV-1 encoded GFP protein upon second infection than PIC/Bystander cells upon second infection. Panel **A**) time equal 0 hrs; addition of DHIV3-mCherry. Panel **B**) time equal 24 hrs; addition of DHIV3-GFP. Panel **C**) time equals 48 hrs after DHIV3-mCherry addition, 24 hrs after DHIV3-GFP addition. Panel **D**) time equals 72 hrs after DHIV3-mCherry addition, 48 hrs after DHIV3-GFP addition. Percentage of mCherry cells also producing GFP, compared to cells producing mCherry only, is always 2 to 5 times higher than the percentage of cells making only GFP, compared to those cells not producing either mCherry or GFP.

## Discussion

Early research into the HIV-1 life cycle identified transcription of HIV-1 PIC cDNA in T-cells and macrophages [5–12]. The unintegrated viral DNA can take several forms, linear and the 1-LTR- and 2-LTR circles [7, 45], with much of the transcription thought to emanate from 1- LTR circles. In macrophages it has been shown that HIV-1 PIC cDNA can persist and be actively transcribed for months. However, it is generally agreed that PIC HIV-1 transcription in macrophages does not routinely produce infectious virus [12]. The use of single cell techniques has enabled us to quantify both the numbers of cells expressing given HIV transcripts in a mixed culture, and also the relative transcript loads of each HIV-1 gene in each infected cell [13–15]. It also allows us to put the cells containing viral transcripts into the context of their background transcriptomes. This provides the opportunity to compare cells producing late viral proteins to those producing transcript but not late viral proteins in order to better understand the cellular metabolic background necessary for virus production.

We show that scRNA-seq can differentiate between macrophage cells that transcribe from PIC HIV-1 cDNA and macrophage cells transcribing from integrated HIV-1 provirus. In proving this observation, we discovered that synthesis of many HIV-1 proteins is only detectable in provirus cells, even though HIV-1 RNA transcripts can be detected equally in cells transcribing from provirus or PIC HIV-1 cDNA. This is consistent with the literature that finds virus production only from provirus [12]. Single cell RNA sequencing can distinguish between the two cell types because the background transcriptomes vary dramatically. PIC cell transcriptomes are characterized by NFkB and AP-1 promoted transcription, while transcriptomes of cells transcribing from provirus are characterized by E2F family transcription products. We also find that the transcriptomes of PIC cells and Bystander cells are identical, suggesting that PIC cells are “oblivious” to the fact that they are making HIV-1 transcripts. Furthermore, the presence of pathogen alters the transcriptome of the uninfected Bystander cells, so that they are clearly distinguishable from true Control cells (cells not exposed to any pathogen). Thus, to understand the transcriptional changes caused by HIV-1 infection in these cell populations, it is not appropriate to compare bulk RNA sequencing data from HIV-1/host cell co-cultures to sequencing data from control cultures not exposed to pathogen. The correct comparisons to make are between cells transcribing from provirus and those transcribing PIC HIV-1 cDNA, or Bystander cells.

As reported by Marsh and Wu and colleagues [10], “transcription in the absence of integration is selective and skewed towards certain viral early genes such as nef and tat, with highly diminished rev and vif”. In general, our single cell analysis agrees with this conclusion, but provides a more nuanced picture. In our data, compared to cells in the same co-culture transcribing from provirus, many more PIC cells are producing tat, nef, gag-pol, and env transcripts. However, the level of a given transcript per cell is not detectably different from Provirus cluster cells. In contrast, the prevalence of PIC cells that make rev, gag, and accessory gene transcripts constitute a smaller fraction of the PIC cells. Nevertheless, those few PIC cells that are producing rev, gag and accessory gene transcripts are doing so at levels equivalent to Provirus cells. In Provirus cells, the numbers of cells detectably making fully spliced versus un-spliced transcripts are equal. The levels of these transcripts is on average similar to the levels of transcripts produced by PIC cells. Thus, it is difficult to make the generalization that high overall transcript levels are a trigger for virus production.

Only the Provirus cells appear to be making p24, mCherry or Vpu protein. Using Western blot analysis, we only see the production of p24, mCherry and Vpu in protein samples containing Provirus cells. It is not clear whether our failure to detect HIV proteins in preparations enriched for PIC /Bystander cells is because very few cells in this cluster were making the proteins (and thus below our level of detection), or if there was some restriction mechanism preventing protein production, or both. Single cell protein analysis technology might be able to address this point. The robust detection of mCherry, p24 and Vpu in protein samples enriched for Provirus cells confirms that late protein synthesis is an attribute of the Provirus cells. Nef protein is reported in the literature to be detectable in protein from both Provirus and PIC cluster cells [10]. Nef protein in not made in cells infected with the DHIV3-mCherry, due to the mCherry sequence replacing the 5’ portion of the nef gene, and we could not detect Nef protein from any DHIV3 infected cultures.

Integrated provirus transcription is required for virus production [12]. It is regulated by promoter elements in the HIV LTR. Thierry and co-workers have recently shown that PIC cDNA and provirus are differentially responsive to NFkB promotion [45]. As others have found, they report provirus transcription is enhanced by NFkB and AP1 binding. However, they find PIC cDNA transcription to be inhibited by NFkB activation. Our data adds an additional layer of complexity to this picture. We can show that early and late gene transcripts are detectable in Provirus cells, at levels equal to the respective transcripts detected in PIC cells. We also show that, in cells that have transitioned from PIC/Bystander to Provirus, the background transcriptome reflects an overall down regulation of NFkB and AP1 transcripts, and upregulation by E2F family promoted transcripts. E2F is not a promoter in the HIV-1 LTR, and we do not suggest that E2F regulation of provirus transcription is the key to the PIC to Provirus transition. However, we do propose that an E2F family promoter-dominated transcriptome is required for virus production.

This proposition appears counter to literature, in which E2F is thought to suppress viral transcription [25, 46]. Nevertheless, the pathways upregulated by E2F are those consistent with what one would intuitively anticipate a being required for viral production. The recognition of E2F promoter family proteins in virus production is not new and is consistent with literature citing a role for Rb phosphorylation and E2F activation in HIV-1 linked tumorigenesis [47]. Indeed, we show that levels of phospho-Rb are higher in Provirus cells. Consistent with this was our observation in our THP-1 system, and in primary cultures of human macrophage and T-cells, that cells that have already committed to making virus are more likely than PIC cells to make virus upon second infection. If this interpretation is correct, then study of genome-wide interactions that accompany provirus integration and amplify E2F signaling might be key to understanding the switch in transcriptome necessary for viral protein production. Several recent scRNA-seq studies using donor T-cell preparations have revealed complicated relationships between T-cell transcriptome and HIV-1 production. In general these studies point to heterogeneity in latent and active HIV-1 infected cells [48-50]. Here, we focused on the distinction between GFP positive cells (transcribing from provirus) compared to GFP negative cells (either Bystander cells or those transcribing from PIC HIV-1 cDNA), in a consistent cell culture model. We believe our approach added clarity to the interpretation of data that is complicated by many distinguishable, possibly stochastic, cell clusters of unknown origin or relevance.

## Materials and methods

### Reagents

THP-1 cells, a monocytic cell line,were obtained from ATCC (Cat#TIB-202). HyClone^™^ RPMI 1640, kanamycin sulfate, Corning^™^ Accutase^™^ detachment solution and phorbol 12-myristate 13-acetate (PMA) were obtained from Fisher Scientific (cat. # SH30011.03, BP906-5, MT25058CI, and BP685-1, respectively). Fetal bovine serum was purchased from Atlanta Biologicals (cat. # S11150). BD Horizon^™^ Fixable Viability Stain 450 was obtained from BD Biosciences (cat. # 562241).

### Cell culture

THP-1 cells were grown in RPMI 1640 supplemented with 10% FBS and KAN (50 μg/mL) at 37°C, 5% CO_2_.

### Generation of DHIV3-mCherry

DHIV3-mCherry virus was generated by calcium phosphate transfection (25). In brief, HEK293FT cells were grown to 80% confluence. DHIV3-mCherry plasmid and VSVg plasmid were mixed with calcium chloride (2.5 M) and HEPES buffered saline solution. The calcium phosphate-DNA suspension was added dropwise to the cells. Chloroquine (100 mM) was subsequently added. HEK293FT cells were incubated overnight with solution. The medium was replaced with fresh DMEM and incubated for an additional 48 hours. Supernatant was collected and filtered (0.45 μm). Optimal viral titers were determined by titrating the virus in THP-1 cells. A titer volume of 100 μL in 500 μL total (∼5 x 106 TU/mL in THP-1s) was chosen due to its high DHIV infection and minimal effects on viability. Titer volume (up to 400 μL of 500 μL) was increased as infectivity fell off in stocks over time.

### DHIV3-mCherry and THP-1 co-culture

For the THP-1s, cells were preincubated overnight in PMA (20 ng/mL) at 500,000 cells/well in order to generate differentiated macrophages. Medium was replaced for THP-1 cells 1 hour prior to the addition of DHIV3-mCherry. Cells were incubated at 37°C for 24 hours followed by preparation for analysis by flow cytometry and cell imaging.

### Flow cytometry

Adherent THP-1 cells were incubated with Accutase^™^ for 15 minutes at 37°C. THP-1 cells were transferred to 5-mL tubes and washed with PBS. Cells were resuspended in BD Horizon™ Fixable Viability Stain 450 (0.25 μg/mL) and incubated at 4°C for 30 minutes. Cells were then fixed in 2% formaldehyde for 30 minutes at 4°C. Cells were analyzed using a FACS Canto. Percent infection by DHIV was quantified as a subset of the live population (FSC/V450/50-). Gates for infection were set according to the uninfected “mock” THP-1 cell controls. Population analysis was then done using FlowJoTM v10.7.

### 10X Genomics library construction and sequencing

Two biological replicate cultures of HIV(+) THP-1 cells (HIVreplicate1 and HIVreplicate2) and HIV(-) THP-1 cells (here after referred to as Control, or WT1 and WT2 for wild type) were processed through the 10X Genomics Chromium Single Cell Controller with Single Cell Gene Expression 3’ Solution (v2 chemistry). Sequencing was done on an Illumina HiSeq 2500 instrument.

Cell suspensions were partitioned into an emulsion of nanoliter-sized droplets using a 10X Genomics Chromium Single Cell Controller and RNA sequencing libraries were constructed using the Chromium Single Cell 3’ Reagent Kit v2 (10X Genomics Cat#PN-120237). Briefly, droplets contained individual cells, reverse transcription reagents and a gel bead loaded with poly(dT) primers that include a 16 base cell barcode and a 10 base unique molecular index. Lysis of the cells and gel bead enables priming and reverse transcription of poly-A RNA to generate barcoded cDNA molecules. Libraries were constructed by End Repair, A-Tailing, Adapter Ligation and PCR amplification of the cDNA molecules. Purified cDNA libraries were qualified on an Agilent Technologies 2200 TapeStation using a D1000 ScreenTape assay (Agilent Cat#5067-5582 and Cat#5067-5583). The molarity of adapter-modified molecules was defined by quantitative PCR using the Kapa Biosystems Kapa Library Quant Kit (Kapa Biosystems Cat#KK4824).

HiSeq 125 Cycle Paired-End Sequencing v4: Sequencing libraries (25 pM) were chemically denatured and applied to an Illumina HiSeq v4 paired end flow cell using an Illumina cBot. Hybridized molecules were clonally amplified and annealed to sequencing primers with reagents from an Illumina HiSeq PE Cluster Kit v4-cBot (PE-401-4001). Following transfer of the flowcell to an Illumina HiSeq 2500 instrument (HCS v2.2.38 and RTA v1.18.61), either a 26x100 cycle or 125 cycle paired-end sequence run was performed using HiSeq SBS Kit v4 sequencing reagents (FC-401-4003). Basic html (notebook) files describing coding and QC data are provided in Appendix IV.

### Data analysis for UMAP (Fig. 2) of HIVrepeat1

Raw FASTQ files from 10x Genomics from were processed by 10x Genomics’ Cell Ranger software. Each library was processed with ‘cellranger count’ pipeline with a common genomic reference made up of human and HIV genomes as well as mCherry. The human genomic reference was GRCh38 with gene annotation from Ensembl release 91, where only features with gene_biotype:protein_coding were kept. The HIV genome and annotation was acquired from NCBI genome (RefSeq ID NC_001802.1). No warnings were issued by 10x Genomics regarding sequencing, alignment, or cell-based QC metrics; however, the samples could have been sequenced deeper as reflected in the sequencing saturation statistics.

In attempt to recover those (perhaps lower quality) GEM partitions, the raw gene-barcode matrices from ‘cellranger count’ (located in ‘outs/raw_gene_bc_matrices’) was processed with the EmptyDrops algorithm (R package DropletUtils v1.2.2) to discriminate cells from background GEM partitions at a false discovery rate (FDR) of 0.001% [51]. GEM partitions with 2000 UMI counts or less were considered to be devoid of viable cells, while those with at least 10,000 UMI counts were automatically considered to be cells.

For each technical replicate, additional quality control measures were taken to filter out low-quality cells. Cell-based QC metrics were calculated with R package scater (v1.16.2) using the perCellQCMetrics function. Cells with extremely low UMI counts, extremely low gene counts or extremely high percentage of expression attributed to mitochondrial genes were also flagged as low quality. Extremeness in any of these three measures was determined by 3 median absolute deviations from the median with the scater isOutlier function. These cells suspected of being low quality were removed from downstream analysis with one exception in the HIV replicates; if cells exhibited above median HIV gene expression, they were not discarded as these were thought to hold potential value as examples of cells in which viral replication suppressed other gene expression (not observed). Further analysis of the HIV biological replicates showed there were remaining low quality cells as marked by unusual mitochondrial gene expression or low library size, which were removed to improve the signal to noise ratio. Specifically, HIV-infected cells with mitochondrial expression of 7.23% and above were removed as well as cells that have less than 3242 UMI counts. To ensure no cell type was discarded due to filtering, average gene expression was compared gene-wise between discarded and kept cells in scatter plots. There were no genes of interest that exhibited markedly different average gene expression between the discarded and kept cells, suggesting the filtering did not remove interesting sub- populations. After filtering low quality cells, technical replicates were combined into biological replicates HIVreplicate1, HIVreplicate2, wt1 and wt2.

For each biological replicate, cells were normalized [51] and scored for a number of important attributes. Each cell from HIV biological replicates was assigned a HIV activity score with Seurat’s (v3.2.2) AddModuleScore function [53], where a high score indicates HIV gene expression was stronger in the cell relative to randomly selected genes of similar expression strength in the biological replicate. Cell cycle phases and scores were assigned cells with the cyclone method [54]; Cells were scored against a simulated doublet population of cells with scran’s (v1.10.2) doubletCells function; those cells with extremely high doublet scores (5 median absolute deviations) were removed and remaining cells were re-normalized again with scran’s quickCluster and scater’s normalize methodology [51] for differences in sequencing depth between libraries.

### Data analysis for t-Sne insert (Fig. S-3)

10x Cell Ranger raw sequencing data was processed into UMI counts with; using the ‘mkfastq’, ‘count’, bioinformatic modules. Cell Ranger de-multiplexed cDNA libraries into FASTQ files with Illumina’s bcl2fastq and aligned reads to a hybrid genomic reference composed of human (Ensemble GRCh38), HIV (NCBI ID: NC_001802.1), and mCherry genomic references with STAR aligner [51, 57]. CellRanger filtered cell barcodes and unique molecular identifiers (UMIs) in estimation of gene-cell UMI counts using only reads that mapped uniquely within the transcriptome. We specified an ‘expected cell number’ of 3000 per library based on reported cell recovery rates.

The QC metrics reported by Cell Ranger indicated that our library construction was a success; the libraries averaged 97.9% valid cell barcodes, 60.6% of reads mapping to the transcriptome, and reported a median of 2402 genes detected per cell (mean of 15262.2 genes per library). Only in HIV-infected samples did reads map to the HIV genome. Cell Ranger also evaluated dimension reduction, clustering, and differential gene expression analysis under default parameters. For further details of the Cell Ranger data processing and analysis pipeline, see https://support.10xgenomics.com/single-cell-gene-expression/software/pipelines/latest/algorithms/overview. Some interactive data analysis, i.e. t-Sne visualization, was conducted with 10x Genomics’ Cell Loupe Browser.

Due to cost, this study was limited by low sequencing depth (mean of 41,433 reads per cell per library). This limitation was mitigated by removing genes with low sequencing coverage; specifically, a gene was filtered out if it did not have 1 percent of cells reporting at least 3 UMI; cells were filtered out if they did not have at least 200 genes with a UMI count. Cells with exceedingly high (top 2%) ribosomal and/or mitochondrial content were filtered out. To reduce mutliplets contaminating analysis, cells with the top 2.3% total UMI were removed (see 10x Genomics benchmarks).

### Dimension reduction, clustering, and differential expression

Highly variable genes were identified with scran’s trendVar and decomposeVar. Specifically, loess smoothing was applied to the gene variance (dependent variable) and the mean gene expression (independent variable) after having corrected for the % mitochondrial expression and cell cycle effects on the cells. Genes with average expression below the first quartile were filtered out of consideration. Gene variance was decomposed into biological and technical components, where genes with variance above the mean trend (loess fit) were assumed to possess biological variation [55]. This process was repeated for the HIV biological replicates separately followed by scran’s combineVar function applied to the combined HIV replicates to identify genes estimated to have positive biological variation and controlled with a false discovery rate of 0.05.

### Gene set enrichment analysis

A gene set enrichment analysis (GSEA) was conducted with R package fgsea (v1.8.0) [56]. The log2 fold change vector of strong-HIV vs. weak-HIV was evaluated for enrichment against 3 different collections of MSigDB gene sets; namely, Hallmark, REACTOME, and Transcription Factor Targets. The Benjamini-Hochberg false discovery rate (FDR) was controlled at 10%.

A DGE and GSEA analysis was conducted on HIV biological replicates in comparison to Control biological replicates. The wild type replicate expression data was subject to (nearly) identical quality control and identical pre-processing steps. As described above, quality control on HIV cells was subject to a greater degree of scrutiny.

To mitigate biases due to potential batch-specific variation, DGE and GSEA analyses contrasting Provirus -HIV and PIC/Bystander-HIV populations to Control cells leveraged consensus between various pairwise contrasts. For instance in the HIV- Provirus vs. Control (WT) contrast, t-tests were evaluated in 4 distinct contrasts: (i) HIVreplicate1-Active vs. WT1; HIVreplicate1-Act Provirus ive vs. WT2; (iii) HIVreplicate2- Provirus vs. WT1; (iv) and HIVreplicate2- Provirus vs. WT2. The scran function combine Markers performed a meta-analysis across the 4 contrasts with the Simes method. The Simes meta-analysis tested whether any of the 4 contrasts manifest either a change a gene-wise expression for DGE analysis; that is to say the meta-analysis p-value encodes the evidence against the null hypothesis, which assumes the gene is not changed in any of the 4 comparisons. Similarly in the GSEA analysis, the log2 fold change statistics from the 4 comparisons were tested for enrichment of the 3 previoulsy mentioned MSigDB gene set collections (Hallmark, REACTOME, Transcript Factor Targets). The results were also merged with the Simes meta-analysis. This same strategy for the HIV- Provirus vs. Control contrast was repeated in the HIV- PIC/Bystander vs. Control comparison.

There was interest in modeling the progression of infection from wild type to PIC/Bystander-HIV to Provirus -HIV clusters. Specifically, there was interest in identifying what genes exhibit a variable expression profile or non-constant trend when ordered from Control to PIC/Bystander-HIV to Provirus -HIV. Macnair and Claassen [41] developed a supervised psuedotime R package, called psupertime, that is tailored to this express purpose. In particular, a penalized logistic ordinal regression model was fit to the combined HIV and Control data. The input data was a subset of highly variable genes identified in the same manner as described above, but including Control data as well. The gene expression data had been normalized, log2 transformed and followed by linear correction of effects due to percent mitochondrial expression and cell cycle phase. The model was able to clearly order cells that reflects the expected order of Control then PIC/Bystander-HIV then Provirus -HIV. The psupertime method also reports a small set of genes which strongly associate with the expected progression, which is based on the magnitude of the penalized coefficients in the logistic ordinal regression.

### Western protocol

The stored THP-1 (uninfected, HIV-1 infected, HIV-1 infected with MK-2048 treatment) cells were pelleted by centrifugation for 5 min at 10,000 x g. Precipitates were then washed twice in cold PBS. Afterwards, cells were lysed using 50 mM Tris pH 7.4, 100 mM NaCl and 2 mM EDTA, complete® protease inhibitor, 2 mM NaF, 2 mM sodium orthovanadate and 10% SDS. Protein concentrations were determined using bovine serum albumin standard and Coomassie Plus Protein Reagent from Pierce Biotechnology (Rockford, IL). 10 µg of the whole cell lysate was separated using NuPAGE 4−12% Bis-Tris gradient gels (Invitrogen Life Sciences, Carlsbad, CA) and transferred to a PVDF membrane (Millipore, Billerica, MA). These were then blocked is 2% BSA in TBST for 20 min at room temperature, incubated overnight with primary antibodies at 4°C in blocking buffer solution and with the secondary antibody for 45 minutes at room temperature. Protein was detected using chemiluminescence and blots were visualized using a Protein Simple FluoroChem M system.

### Antibodies and fluors

P24 and Gag protein precursor production was detected with monoclonal mouse IgG- AG3.0, (NIH AIDS Res. Reagent Prog., Germantown, MD; Cat. # 4121), 1:500, and using an AlexaFlour 633nm or 700nm goat anti-mouse IgG (H+L) secondary anti-body (Fisher Scientific, Pittsburgh, PA) for flow cytometry. Anti-HIV-1 NL4-3 VPU rabbit polyclonal, also from the NIH AIDS Res. Reagent Prog. (Cat. # 969), was used at 1:20 dilution for Western Blots. Goat anti-mCherry, OriGene (TR150126), was used at a 1:5000 dilution for Western Blots. ThermoFisher Scientific Anti-Rb (LF-MA0173, 32C8) was used at a 1:1000 dilution. Invitrogen anitbodies: anti-pRB (T821, 710314) was used at a 1:500 dilution, anti-pIκB (S32, 701271) was used at a 1:500 dilution, and anti-B-actin (PA5-85291) was used at a 1:5000 dilution.

Other antibodies obtained from the NIH AIDS Res. Reagent Prog., Germantown, MD include: anti-HIV-1 IIIB gp120 Polyclonal (Cat. # 57), anti-HIV-1 RF gp160 Polyclonal (HT7) (Cat. # 189), anti-HIV-1 Tat Polyclonal (Cat. # 705), anti-Nef Monoclonal (EH1) (Cat. # 3689), anti-HIV-1 Nef Polyclonal (Cat. # 2949), anti-HIV-1 Vpr 1-50 aa Polyclonal (Cat. # 11836); anti-HIV-1 HXB2 Vif Polyclonal (Cat. #12256), anti-HIV-1 HXB2 IN Polyclonal (Antigen 2) (Cat. # 12877), anti-HIV-1 RT Monoclonal (MAb 21) (Cat. # 3483), anti-HIV-1 HXB2 RT Polyclonal (Antigen 2) (Cat. # 12881), anti-HIV-1 Protease Polyclonal (Cat. # 13564), anti-HIV-1 Rev Monoclonal (1G7) (Cat. # 7376), which were tested at various dilutions.

HRP-conjugated secondary antibodies were used to develop Western Blots, anti-rabbit A0545 (1:5000) and anti-mouse A9044 (1:5000) from Sigman Chem. Co., and anti-goat 401515 (1:10,000) from CalBiochem. Propidium Iodide was obtained from Molecular Probes, Eugene, OR. DAPI blue was obtained from Acros Organics, Geel, Belgium.

### Real-time PCR methods

For detection of integrated proviral DNA, DNA from control, DHIV3-mCherry infected, DHIV3-mCHerry infected plus integrase inhibitor treated THP1 cells was purified using Qiagen Blood and Tissue DNeasy kits. The PCR evaluation of integrated HIV was performed using the primers and PCR conditions described from Chun et al. [34]. Briefly, the primers were:

Alu->LTR 5, 5’-TCCCAGCTACTCGGGAGGCTGAGG-3’

LTR->Alu 3’, 5’-AGGCAAGCTTTATTGAGGCTTAAGC-3’

Nested secondary PCR primers (generating a 352 bp ampilicon):

5’, 5’-CACACACAAGGCTACTTCCCT-3’

3’, 5’-GCCACTCCCCIGTCCCGCCC-3’

We used Ranger polymerase and buffer conditions (Meridian Bioscience, Thomas Scientific, Swedesboro, NJ) for the long-range PCR with Alu-LTR primer sets, and BioTaq polymerase and buffer conditions (Meridian Bioscience, Thomas Scientific, Swedesboro, NJ) for the nested PCR. The integrated HIV PCR was performed on an MJ PTC-200 with an MJR 2X48 and a Chromo-4 alpha unit for the long-range and nested PCR, respectively. The nested PCR was performed as a real-time assay using SYBR Green I to detect the amplicon progression curves and evaluate the melting curve.

For detection of total HIV DNA, in order to determine if comparable total HIV DNA was present in the samples the same samples described above, we utilized the 5’ nested primer (5’- CACACACAAGGCTACTTCCCT-3’) along with the LTR->Alu 3’ primer (5’- AGGCAAGCTTTATTGAGGCTTAAGC-3’) using PCR conditions similar to the nested PCR described above but with a 30 sec extension time for the 484 bp amplicon. This PCR was performed on a Roche LightCycler 480 instrument using BioTaq polymerase and buffer conditions with SYBR Green I detection of the 484 bp amplicon.

Detection of circular 2-LTR DHIV3-mCherry PIC DNA was performed as described in Brussel and Sonigo [33]. Briefly, the primers used were:

HIV F, 5’ GTGCCCGTCTGTTGTGTGACT 3’

HIV R1, 5’ ACTGGTACTAGCTTGTAGCACCATCCA 3’.

Initially we performed the PCR conditions used by Brussel and Sonigo [33], using BioTaq polymerase and buffer conditions with a 25 sec extension time and SYBR Green I detection, but we were unable to detect any amplicon 2LTR circle PIC product. Following the detection of the total PIC HIV data, we hypothesized that the 2-LTR content in these samples could be considerably lower at this 24 hr time point. Therefore, we reran the PCR again a second time and were able to detect an expected 231 bp amplicon. To verify this approach, we synthesized new PCR primers:

RU5 forward: 5’-GCTTAAGCCTCAATAAAGCTTGCCT-3’ (this is the compliment of the LTR->Alu primer described above from Chun et al. [34]).

U3 reverse: 5’-ACAAGCTGGTGTTCTCTCCT-3’.

This primer set also did not generate the 2-LTR circular PIC amplicon within 50 PCR cycles but was designed to encompass the amplicon generated by Brussel and Sonigo primers above. Therefore, we used the HIV F and R1 primers as a nested set and ran a 1:20 dilution of this amplification for another 50 PCR cycles to obtain the expected 231 bp amplicon. This demonstrates the 2-LTR circular form of PIC cDNA was present at low levels in all the conditions where DHIV3-mCherry was used, but not in the control samples.

### Sequential DHIV3-mCherry, DHIV3-GFP infection

PBMCs from healthy human donors were isolated using lymphocyte separation medium (Biocoll separating solution; Biochrom) or lymphoprep (Stemcell). CD4+ T cells were negatively isolated using the RosetteSep™Human CD4+ T Cell Enrichment Cocktail (Stem Cell Technologies) or the EasySep™ Human Naïve CD4+ T Cell Isolation Kit (Stem Cell Technologies) according to the manufacturer’s instructions. Primary rCD4s were cultured to a density of 5 x 106/mL in RPMI-1640 medium containing 10% FCS, glutamine (2 mM), streptomycin (100 mg/mL), penicillin (100 U/mL) and interleukin 2 (IL-2) (10 ng/mL).

Monocyte-derived macrophages (MDMs) were obtained by stimulation of PBMC cultures with 15 ng/mL recombinant human M-CSF (R&D systems) and 10% human AB serum (Sigma Aldrich) in DMEM supplemented with glutamine (2 mM), streptomycin (100 mg/mL) and penicillin (100 U/mL) for 6 days.

### Statistical analysis

The pairwise TTests function from Scran was used to determine statistically significant differential expression of genes between groups. This was performed for all comparison sets. Only those genes which were significantly different were included in Hallmark, REACTOME, pseudotime, psupertime and TFT analyses. Other default statistical standards were adopted from the various software recommendations during data analyses unless otherwise specified.

## Acknowledgments

This work was supported by the University of Utah HCI Flow Cytometry Facility and Metabolomics core facilities, and the HSC High Throughput Genomics core facility. The help of Christopher J. Conley and Brian Lohman in bio-statistical analysis is gratefully acknowledged. Mr. A.L. Lim was partially supported during this work by the grant U01TW008163 (P.I., Margo Haygood, University of Utah). This work was also supported in part by R21 AI124823 (LRB & VP) and a University of Utah seed grant (LRB).

## Supporting Information Figure Captions (S figures)

**S-1 DHIV3-mCherry map.** Snapgene [58] map of DHIV3-mCherry plasmid.

**S-2 Seurat analysis of biological repeats HIVreplicate1 and HIVreplicate2.** HIVreplicate1a and HIVreplicate1b, and HIVreplicate2a and HIVreplicate2b, are technical repeat data. Technical repeats were conducted with each experiment. Figure shows PC analysis of biological repeat experiments. Technical duplicates were not different and so were combined for each repeat

**S-3 tSne projection of scRNA seq data from experiment HIVreplicate1**. Seurat analysis and t-SNE projection of data shown in Figure 2. Viral transcript numbers (**h**) were determined for cells containing any detected HIV-1transcript, as described in methods. Orange dots represent high-level transcript load per cell, greater than 8 transcripts mapping to HIV-1 genes per cell, green dots indicate cells with lower transcript loads detected per cells, and blue dots indicate cells with no detectable HIV-1 transcripts. Barcodes of cells in Provirus Cluster (A) tracked to Provirus Cluster cells in UMAP analysis (Fig. 2).

**S-4 Unsupervised clustering of UMAP shown in** Fig 4. Left panels shows unsupervised clustering obtained at K values from 10 to 130. Right panels show Violin plots of HIV-1 transctipts/cell in the clusters identified at the specified K values (Scran’s buildSSNGraph using the PCA as input). Clusters 6 and 8 at K equal to 10 contained most of the cells in the semi supervised Provirus cluster (circled in red) and were used to define Provirus transcriptome, versus the remaining cells making up the semi supervised PIC/Bystander cluster. Stipulation of lower K values means that during analysis any one given cell is clustered with a smaller number of cells with similar transcriptomes.

**S-5 Real-time PCR analysis of DNA samples from Control, DHIV3-mCherry infected, and DHIV3-mCherry infected, integrase inhibitor treated THP-1 cells.** MW, molecular weight markers. Lanes 1, Control cell DNA; Lane 2, DNA from DHIV3-mCherry infected culture cells; Lane 3, DNA from DHIV3-mCherry infected cultures treated with integrase- inhibitor (25nM MK-2048). 100 ng of DNA was tested in each amplification unless noted. Primers used are described in Methods. Panels **A**, **B** and **C**) PCR demonstration of integrated proviral HIV DNA. Panel **A**) examples of the progression curves; upper curve represent lanes 2 at 200 and 100 ng DNA respectively, middle curves reflect lanes 3, respectively, bottom 2 curves were generated by Control DNA. Panel **B**) melting curves; the upper curves represent lanes 2 at 200 and 100 ng and lane 3 at 200 ng DNA respectively. Panel **C**) shows the amplicons generated from the integrated DNA using the nested PCR strategy described by Chun et al.[34] on a 1% agarose gel. The amplicon product sizes matched the predicted product size of 352 bp. These examples were from two biological replicates, one using 200 ng and one starting with 100 ng of starting DNA purified using Qiagen Blood and Tissue DNeasy kits. The agarose gel shows the integrated proviral DNA, assessed using an MJ PTC-200 thermal cycler and the nested PCR was evaluated using a Chromo-4 alpha unit. Note that the 200 ng samples with integrase inhibitor (Lane 3) show a small amount of integrated provirus DHIV3-mCherry DNA, demonstrating that the inhibitor did not completely inhibit the DHIV-mCherry integration. This is consistent with the 48 hr gag/p24 protein production seen in the immunoblot analysis (Fig. 7), and flow cytometry analysis (Fig. 6). Panels **D**, **E**, and **F**) show the same DNA samples used to detect 2- LTR circle PIC cDNA from the second or two consecutive PCR runs. Panel **D**) Lower 3 curves show progression curves with lack of 2LTR primer products in Control DNA, while all 6 biological repeats, 3 from DHIV3-mCherry infected cultures and 3 from DHIV3-mCherry infected cultures treated with integrase inhibitor show amplification of PIC cDNA p2LTR products. Panel **E**) shows the melting curves for these amplification products with the lower 3 curves representing Control DNA, the lowest curve representing Control DNA from the nested PCR approach (see Methods). Panel **F**) shows the amplicons generated run out on a 1% agarose gel. In these experiments the PCR was assessed using a Roche LightCyler 480. Biological replicates of 100 ng starting DNA are represented as “rep 1” and “rep 2”, using the HIV F and R1 primers of Brussels and Sonigo [33]. In the confirmation experiment lanes, under the “nested” label in the agarose gel, wider bracketing primers were used in the first amplification followed by the HIV F and R1 primers “nested” in the second run. Perhaps not surprising after 100 cycles, there are contaminating PCR products in the Control lanes; however, the expected 231 bp amplicon is not detectable in the Control cell DNA, while it is the predominant product in DNA from either infected, or infected and integrase inhibitor treated cell DNAs. Panels **G, H** and **I**) show the same DNA samples used to detect total DHIV-mCherry DNA, also assessed using the Roche LightCyler 480. Primers used are described in Methods. Panel **G**) shows the progression curves generated in this experiment, the lowest 2 curves represent Control cell DNA. Panel **H**) shows the melting curves for these products, again the lowest 2 curves are from Control cell DNA. Panel **I**) shows the amplicons generated, of predicted size, run out on a 1% agarose gel. The real-time PCR results show roughly equivalent total amounts of DHIV3-mCherry in infected culture DNAs, whether in the presence of integrase inhibitor or not. This indicates that overall, the total amounts of PIC cDNA are similar in integrase inhibitor treated and untreated DHIV3-mCherry infected cultures.

**S-6 Unsupervised clustering of integrase inhibitor treated DHIV3 infected cells from** Figure 8. No cluster corresponding to the Provirus cluster identified in HIVreplicate1 or HIVreplicate2 could be identified, regardless of K value specified. Data were analyzed as in Figure 4. Left panels shows unsupervised clustering obtained at K values from 10 to 130. Right panels show Violin plots of HIV-1 transctipts/cell in the clusters identified at the specified K10 values (Scran’s buildSSNGraph using the PCA as input). Stipulation of lower K values means that during analysis any one given cell is clustered with a smaller number of cells with similar transcriptomes.

**S-7 Feature plots of integrase inhibitor treated cultures.** Transcripts of DHIV3-mCherry infection readily detectable in the presence of integrase inhibitor Panel **A**. Effects of cell cycle (Panel **B)**, mitochondrial gene expression (Panel **C**) and number of genes detected per cell (Panel **D**) shown for comparison to Figure 3.

**S-8 Biological repeat experiments HIVreplicate1 and HIVreplicate2.** UMAP projections of Seurat analysis of biological repeat experiments HIVreplicate1 (Panel **A**) and HIVreplicate2 (Panel **B**). Seurat analysis yielded 8.1% of cells in Provirus cluster from experiment HIVreplicate1, 6% of cells in Provirus cluster in repeat HIVreplicate2, in agreement with percentages of mCherry positive percentages obtained for duplicate cultures analyzed by flow cytometry.

**S-9 Unsupervised clustering of HIVrepeat2 UMAP projection.** Left panels shows unsupervised clustering obtained at K nearest neighbor values from 10 to 130. Right panels show Violin plots of HIV-1 transctipts/cell in the clusters identified at the specified K values (Scran’s buildSSNGraph using the PCA as input). Cluster 3 at K equal to 10 contained most of the cells in the semi supervised Provirus cluster (circled in red) and was used to define Provirus transcriptome, versus the remaining cells making up the semi supervised PIC/Bystander cluster. Stipulation of lower K values means that during analysis any one given cell is clustered with a smaller number of cells with similar transcriptomes.

**S-10. The Distribution of HIV-1 transcripts throughout Provirus and PIC/Bystander clusters of HIVreplicate2.** Feature plot showing distribution of cells from UMAP in Fig. S-8 containing detectable DHIV3-mCherry transcripts. As describe above, these UMAP projections were made with Seurat’s FeaturePlot function. They are colored by expression of individual genes (UMAP projection colored by walktrap, normalized log2 values). ASP is a negative control, bacterial gene transcript sequence. The distribution repeats the results obtianed in HIVrepeat1 (Fig. 10A).

**S-11 Differential gene expression comparison of Provirus and PIC cluster gene transcripts from biological repeat experiments.** Consistent positive correlation of common DGE in HIVreplicate1 (abscissa) versus HIVreplicate2 (ordinate) repeat experiments (Spearman’s rank correlation coefficient of all common genes 0.384), agreed with Hallmark and REACTOME analyses that showed similar pathways up- or down- regulated in the Provirus versus PIC/Bystander clusters of the biological repeat experiments.

**S-12 Sequential infections of primary human lymphocyte and macrophage cultures.** Active, mCherry positive, cells were 2 to 5 times more likely to make HIV-1 encoded GFP protein upon second infection than PIC/Bystander cells upon second infection. Panels **A**, **H**) primary cultures of T-lymphocytes and macrophages at time equals 0 hrs, respectively. Panels **B**-**D**) primary lymphocytes infected with low titer DHIV3. Panels **E**-**G**) primary lymphocytes infected with high titer. Panels **H**-**K**) primary macrophages. Percentage of mCherry cells also producing GFP, compared to cells producing mCherry only, is always 2 to 5 times higher than the percentage of cells making only GFP, compared to those cells not producing mCherry. Panel **A** and **H**) Time equal 0 hrs; addition of DHIV3-mCherry. Panel **B**, **E** and **I**) time equal 24 hrs; addition of DHIV3-GFP. Panel **C**, **F** and **J**) time equals 48 hrs after DHIV3-mCherry addition, 24 hrs after DHIV3-GFP addition. Panel **D**, **G** and **K**) time equals 72 hrs after DHIV3- mCherry addition, 48 hrs after DHIV3-GFP addition.

## References

1. Finzi D, Hermankova M, Pierson T, Carruth LM, Buck C, Chaisson RE, Quinn T C, Chadwick K, Margolick J, Brookmeyer R, Gallant J, Markowitz M, Ho DD, Richman DD, Siliciano RF. Identification of a reservoir for HIV-1 in patients on highly active antiretroviral therapy. Science 1997 278:1295–300.

2. Chun TW, Carruth L, Finzi D, Shen X, DiGiuseppe JA, Taylor H, Hermankova M, Chadwick K, Margolick J, Quinn TC, Kuo YH, Brookmeyer R, Zeiger MA, Barditch-Crovo P, Siliciano RF. Quantification of latent tissue reservoirs and total body viral load in HIV-1 infection. Nature 1997 387:183–8.

3. Wong JK, Hezareh M, Günthard HF, Havlir DV, Ignacio CC, Spina CA, Richman DD. Recovery of replication-competent HIV despite prolonged suppression of plasma viremia. Science. 1997 Nov 14;278(5341):1291–5. doi: 10.1126/science.278.5341.1291. PMID: 9360926.

4. Szaniawski MA, Spivak AM, Bosque A, Planelles V. Sex Influences SAMHD1 Activity and Susceptibility to Human Immunodeficiency Virus-1 in Primary Human Macrophages. J Infect Dis. 2019 Feb 15;219(5):777–785. doi: 10.1093/infdis/jiy583. PMID: 30299483; PMCID: PMC6376916.

5. Pang S, Koyanagi Y, Miles S, Wiley C, Vinters HV, Chen IS. High levels of unintegrated HIV-1 DNA in brain tissue of AIDS dementia patients. Nature. 1990 Jan 4;343(6253):85–9. doi: 10.1038/343085a0. PMID: 2296295.

6. Zhou Y, Zhang H, Siliciano JD, Siliciano RF. Kinetics of human immunodeficiency virus type 1 decay following entry into resting CD4+ T cells. J Virol. 2005 Feb;79(4):2199–210. doi: 10.1128/JVI.79.4.2199-2210.2005. PMID: 15681422; PMCID: PMC546571.

7. Hamid FB, Kim J, Shin CG. Distribution and fate of HIV-1 unintegrated DNA species: a comprehensive update. AIDS Res Ther. 2017 Feb 16;14(1):9. doi: 10.1186/s12981-016-0127-6. PMID: 28209198; PMCID: PMC5314604.

8. Chan CN, Trinité B, Lee CS, Mahajan S, Anand A, Wodarz D, Sabbaj S, Bansal A, Goepfert PA, Levy DN. HIV-1 latency and virus production from unintegrated genomes following direct infection of resting CD4 T cells. Retrovirology. 2016 Jan 5;13:1. doi: 10.1186/s12977-015-0234-9. PMID: 26728316; PMCID: PMC4700562.

9. Wu Y, Marsh JW. Early transcription from nonintegrated DNA in human immunodeficiency virus infection. J Virol. 2003 Oct;77(19):10376–82. doi: 10.1128/jvi.77.19.10376-10382.2003. PMID: 12970422; PMCID: PMC228496.

10. Kelly J, Beddall MH, Yu D, Iyer SR, Marsh JW, Wu Y. Human macrophages support persistent transcription from unintegrated HIV-1 DNA. Virology. 2008 Mar 15;372(2):300–12. doi: 10.1016/j.virol.2007.11.007. Epub 2007 Dec 4. PMID: 18054979; PMCID: PMC2276161.

11. Kruize Z, Kootstra NA. The Role of Macrophages in HIV-1 Persistence and Pathogenesis. Front Microbiol. 2019 Dec 5;10:2828. doi: 10.3389/fmicb.2019.02828. PMID: 31866988; PMCID: PMC6906147.

12. Englund G, Theodore TS, Freed EO, Engelman A, Martin MA. Integration is required for productive infection of monocyte-derived macrophages by human immunodeficiency virus type 1. J Virol. 1995 May;69(5):3216–9. doi: 10.1128/JVI.69.5.3216-3219.1995. PMID: 7707554; PMCID: PMC189028.

13. Hwang B, Lee JH, Bang D. Single-cell RNA sequencing technologies and bioinformatics pipelines. Exp Mol Med. 2018 Aug 7;50(8):96. doi: 10.1038/s12276-018-0071-8. PMID: 30089861; PMCID: PMC6082860.

14. Chen G, Ning B, Shi T. Single-Cell RNA-Seq Technologies and Related Computational Data Analysis. Front Genet. 2019 Apr 5;10:317. doi: 10.3389/fgene.2019.00317. PMID: 31024627; PMCID: PMC6460256.

15. 10X Genomics Home Page https://www.10xgenomics.com/?utm_medium%5B0%5D=search&utm_medium%5B1%5D=search&utm_source%5B0%5D=google&utm_source%5B1%5D=google&utm_campaign=sem-goog-2021-website-page-ra_g-brand-amer&useroffertype=website-page&userresearcharea=ra_g&userregion=amer&userrecipient=customer&usercampaignid=7011P000001PCfa&utm_content=Pure%20Brand&utm_term=10x%20genomics&gclid=Cj0KCQiA3smABhCjARIsAKtrg6L5CIu8nsc24EDzVxlSXMQ1IjYrYk8KP9bBVXEva4QP15dnxLtfwjUaAteHEALw_wcB&gclsrc=aw.ds

16. Sloan RD, Donahue DA, Kuhl BD, Bar-Magen T, Wainberg MA. Expression of Nef from unintegrated HIV-1 DNA downregulates cell surface CXCR4 and CCR5 on T- lymphocytes. Retrovirology. 2010 May 13;7:44. doi: 10.1186/1742-4690-7-44. PMID: 20465832; PMCID: PMC2881062.

17. Wu Y, Marsh JW. Selective transcription and modulation of resting T cell activity by preintegrated HIV DNA. Science. 2001 Aug 24;293(5534):1503–6. doi: 10.1126/science.1061548. PMID: 11520990.

18. Poon B, Chang MA, Chen IS. Vpr is required for efficient Nef expression from unintegrated human immunodeficiency virus type 1 DNA. J Virol. 2007 Oct;81(19):10515–23. doi: 10.1128/JVI.00947-07. Epub 2007 Jul 25. PMID: 17652391; PMCID: PMC2045493.

19. Gelderblom HC, Vatakis DN, Burke SA, Lawrie SD, Bristol GC, Levy DN. Viral complementation allows HIV-1 replication without integration. Retrovirology. 2008 Jul 9;5:60. doi: 10.1186/1742-4690-5-60. PMID: 18613957; PMCID: PMC2474848.

20. Essaghir A, Toffalini F, Knoops L, Kallin A, van Helden J, Demoulin JB. Transcription factor regulation can be accurately predicted from the presence of target gene signatures in microarray gene expression data. Nucleic Acids Res. 2010 Jun;38(11):e120. doi: 10.1093/nar/gkq149. Epub 2010 Mar 9. PMID: 20215436; PMCID: PMC2887972.

21. Banks CJ, Joshi A, Michoel T. Functional transcription factor target discovery via compendia of binding and expression profiles. Sci Rep. 2016 Feb 9;6:20649. doi: 10.1038/srep20649. PMID: 26857150; PMCID: PMC4746627.

22. ENCODE Transcription Factor Targets Dataset https://maayanlab.cloud/Harmonizome/dataset/ENCODE+Transcription+Factor+Targets

23. Gaynor R. Cellular transcription factors involved in the regulation of HIV-1 gene expression. AIDS. 1992 Apr;6(4):347–63. doi: 10.1097/00002030-199204000-00001. Erratum in: AIDS 1992 Jun;6(6):following 606. PMID: 1616633.

24. Schiralli Lester GM, Henderson AJ. Mechanisms of HIV Transcriptional Regulation and Their Contribution to Latency. Mol Biol Int. 2012;2012:614120. doi: 10.1155/2012/614120. Epub 2012 Jun 3. PMID: 22701796; PMCID: PMC3371693.

25. Kundu M, Guermah M, Roeder RG, Amini S, Khalili K. Interaction between cell cycle regulator, E2F-1, and NF-kappaB mediates repression of HIV-1 gene transcription. J Biol Chem. 1997 Nov 21;272(47):29468–74. doi: 10.1074/jbc.272.47.29468. PMID: 9368006.

26. Mbonye U, Karn J. Transcriptional control of HIV latency: cellular signaling pathways, epigenetics, happenstance and the hope for a cure. Virology. 2014 Apr;454–455:328-39. doi: 10.1016/j.virol.2014.02.008. Epub 2014 Feb 22. PMID: 24565118; PMCID: PMC4010583.

27. Frolov MV, Dyson NJ. Molecular mechanisms of E2F-dependent activation and pRB- mediated repression. J Cell Sci. 2004 May 1;117(Pt 11):2173–81. doi: 10.1242/jcs.01227. PMID: 15126619.

28. Bracken AP, Ciro M, Cocito A, Helin K. E2F target genes: unraveling the biology. Trends Biochem Sci. 2004 Aug;29(8):409–17. doi: 10.1016/j.tibs.2004.06.006. PMID: 15362224.

29. Bosque A, Planelles V. Induction of HIV-1 latency and reactivation in primary memory CD4+ T cells. Blood. 2009 Jan 1;113(1):58–65. doi: 10.1182/blood-2008-07-168393. Epub 2008 Oct 10. PMID: 18849485; PMCID: PMC2614643.

30. Hotter D, Bosso M, Jønsson KL, Krapp C, Stürzel CM, Das A, Littwitz-Salomon E, Berkhout B, Russ A, Wittmann S, Gramberg T, Zheng Y, Martins LJ, Planelles V, Jakobsen MR, Hahn BH, Dittmer U, Sauter D, Kirchhoff F. IFI16 Targets the Transcription Factor Sp1 to Suppress HIV-1 Transcription and Latency Reactivation. Cell Host Microbe. 2019 Jun 12;25(6):858–872.e13. doi: 10.1016/j.chom.2019.05.002. Epub 2019 Jun 4. PMID: 31175045; PMCID: PMC6681451.

31. Mascolini M. Merck Offers Unique Perspective on Second-Generation Integrase Inhibitor 10th International Workshop on Clinical Pharmacology of HIV Therapy, Amsterdam, April 15–17, 2009

32. NIH AIDS Reagent Program, Reagent Datasheet Detail: monoclonal mouse IgG-AG3.0, (NIH AIDS Res. Reagent Prog., Germantown, MD; Cat. # 4121) https://www.hivreagentprogram.org/Catalog/HRPMonoclonalAntibodies/ARP-4121.aspx

33. Brussel A, Sonigo P. Evidence for gene expression by unintegrated human immunodeficiency virus type 1 DNA species. J Virol. 2004 Oct;78(20):11263–71. doi: 10.1128/JVI.78.20.11263-11271.2004. PMID: 1 5452245; PMCID: PMC521838.

34. Chun TW, Stuyver L, Mizell SB, Ehler LA, Mican JA, Baseler M, Lloyd AL, Nowak MA, Fauci AS. Presence of an inducible HIV-1 latent reservoir during highly active antiretroviral therapy. Proc Natl Acad Sci U S A. 1997 Nov 25;94(24):13193–7. doi: 10.1073/pnas.94.24.13193. PMID: 9371822; PMCID: PMC24285.

35. Sauter D, Hotter D, Van Driessche B, Stürzel CM, Kluge SF, Wildum S, Yu H, Baumann B, Wirth T, Plantier JC, Leoz M, Hahn BH, Van Lint C, Kirchhoff F. Differential regulation of NF-κB-mediated proviral and antiviral host gene expression by primate lentiviral Nef and Vpu proteins. Cell Rep. 2015 Feb 3;10(4):586–99. doi: 10.1016/j.celrep.2014.12.047. Epub 2015 Jan 22. PMID: 25620704; PMCID: PMC4682570.

36. Langer S, Hammer C, Hopfensperger K, Klein L, Hotter D, De Jesus PD, Herbert KM, Pache L, Smith N, van der Merwe JA, Chanda SK, Fellay J, Kirchhoff F, Sauter D. HIV-1 Vpu is a potent transcriptional suppressor of NF-κB-elicited antiviral immune responses. Elife. 2019 Feb 5;8:e41930. doi: 10.7554/eLife.41930. PMID: 30717826; PMCID: PMC6372280.

37. Schwartz S, Felber BK, Benko DM, Fenyö EM, Pavlakis GN. Cloning and functional analysis of multiply spliced mRNA species of human immunodeficiency virus type 1. J Virol. 1990 Jun;64(6):2519–29. doi: 10.1128/JVI.64.6.2519-2529.1990. PMID: 2335812; PMCID: PMC249427.

38. Sheldon M, Ratnasabapathy R, Hernandez N. Characterization of the inducer of short transcripts, a human immunodeficiency virus type 1 transcriptional element that activates the synthesis of short RNAs. Mol Cell Biol. 1993 Feb;13(2):1251–63. doi: 10.1128/mcb.13.2.1251. PMID: 8423790; PMCID: PMC359010.

39. Svensson V, Natarajan KN, Ly LH, Miragaia RJ, Labalette C, Macaulay IC, Cvejic A, Teichmann SA. Power analysis of single-cell RNA-sequencing experiments. Nat Methods. 2017 Apr;14(4):381–387. doi: 10.1038/nmeth.4220. Epub 2017 Mar 6. PMID: 28263961; PMCID: PMC5376499.

40. Qiu P. Embracing the dropouts in single-cell RNA-seq analysis. Nat Commun. 2020 Mar 3;11(1):1169. doi: 10.1038/s41467-020-14976-9. PMID: 32127540; PMCID: PMC7054558.

41. Psupertime: supervised pseudotime inference for single cell 2 RNA-seq data with sequential labels Will Macnair, Manfred Claassen 28 March 2019 https://www.biorxiv.org/content/10.1101/622001v1

42. Qiu X, Mao Q, Tang Y, Wang L, Chawla R, Pliner HA, Trapnell C. Reversed graph embedding resolves complex single-cell trajectories. Nat Methods. 2017 Oct;14(10):979–982. doi: 10.1038/nmeth.4402. Epub 2017 Aug 21. PMID: 28825705; PMCID: PMC5764547.

43. Street K, Risso D, Fletcher RB, Das D, Ngai J, Yosef N, Purdom E, Dudoit S. Slingshot: cell lineage and pseudotime inference for single-cell transcriptomics. BMC Genomics. 2018 Jun 19;19(1):477. doi: 10.1186/s12864-018-4772-0. PMID: 29914354; PMCID: PMC6007078.

44. Rubin SM. Deciphering the retinoblastoma protein phosphorylation code. Trends Biochem Sci. 2013 Jan;38(1):12–9. doi: 10.1016/j.tibs.2012.10.007. Epub 2012 Dec 3. PMID: 23218751; PMCID: PMC3529988.

45. Thierry S, Thierry E, Subra F, Deprez E, Leh H, Bury-Moné S, Delelis O. Opposite transcriptional regulation of integrated vs unintegrated HIV genomes by the NF-κB pathway. Sci Rep. 2016 May 11;6:25678. doi: 10.1038/srep25678. PMID: 27167871; PMCID: PMC4863372.

46. Kundu M, Srinivasan A, Pomerantz RJ, Khalili K. Evidence that a cell cycle regulator, E2F1, down-regulates transcriptional activity of the human immunodeficiency virus type 1 promoter. J Virol. 1995 Nov;69(11):6940–0. doi: 10.1128/JVI.69.11.6940-6946.1995. PMID: 7474112; PMCID: PMC189612.

47. Ambrosino C, Palmieri C, Puca A, Trimboli F, Schiavone M, Olimpico F, Ruocco MR, di Leva F, Toriello M, Quinto I, Venuta S, Scala G. Physical and functional interaction of HIV- 1 Tat with E2F-4, a transcriptional regulator of mammalian cell cycle. J Biol Chem. 2002 Aug 30;277(35):31448–58. doi: 10.1074/jbc.M112398200. Epub 2002 Jun 7. PMID: 12055184.

48. Bradley T, Ferrari G, Haynes BF, Margolis DM, Browne EP. Single-Cell Analysis of Quiescent HIV Infection Reveals Host Transcriptional Profiles that Regulate Proviral Latency., Cell Rep. 2018 Oct 2;25(1):107-117.e3 doi: 10.1016/j.celrep.2018.09.020. PMID: 30282021; PMCID: PMC6258175.

49. Cohn LB, da Silva IT, Valieris R, Huang AS, Lorenzi JCC, Cohen YZ, Pai JA, Butler AL, Caskey M, Jankovic M, Nussenzweig MC. Clonal CD4^+^ T cells in the HIV-1 latent reservoir display a distinct gene profile upon reactivation., Nat Med. 2018 May;24(5):604-609.e3 doi: 10.1038/s41591-018-0017-7. Epub 2018 Apr 23. PMID: 29686423; PMCID: PMC5972543.

50. Golumbeanu M, Cristinelli S, Rato S, Munoz M, Cavassini, M, Beerenwinkel N, Ciuffi A. Single-Cell RNA-Seq Reveals Transcriptional Heterogeneity in Latent and Reactivated HIV-Infected Cells., Cell Rep. 2018 Apr 24;23(4):942-950. doi: 10.1016/j.celrep.2018.03.102. PMID: 29694901.

51. Lun ATL, Riesenfeld S, Andrews T, Dao TP, Gomes T; participants in the 1st Human Cell Atlas Jamboree, Marioni JC. EmptyDrops: distinguishing cells from empty droplets in droplet-based single-cell RNA sequencing data., Genome Biol. 2019 Mar 22;20(1):63. doi: 10.1186/s13059-019-1662-y. PMID: 30902100; PMCID: PMC6431044.

52. McCarthy DJ, Campbell KR, Lun AT, Wills QF. Scater: pre-processing, quality control, normalization and visualization of single-cell RNA-seq data in R. Bioinformatics. 2017 Apr 15;33(8):1179–1186. doi: 10.1093/bioinformatics/btw777. PMID: 28088763; PMCID: PMC5408845.

53. Butler A, Hoffman P, Smibert P, Papalexi E, Satija R. Integrating single-cell transcriptomic data across different conditions, technologies, and species. Nat Biotechnol. 2018 Jun;36(5):411–420. doi: 10.1038/nbt.4096. Epub 2018 Apr 2. PMID: 29608179; PMCID: PMC6700744.

54. Scialdone A, Natarajan KN, Saraiva LR, Proserpio V, Teichmann SA, Stegle O, Marioni JC, Buettner F. Computational assignment of cell-cycle stage from single-cell transcriptome data. Methods. 2015 Sep 1;85:54–61. doi: 10.1016/j.ymeth.2015.06.021. Epub 2015 Jul 2. PMID: 26142758.

55. loess: Local Polynomial Regression Fitting. https://rdrr.io/r/stats/loess.html

56. Sergushichev AA. An algorithm for fast preranked gene set enrichment analysis using cumulative statistic calculation. 2016 bioRxiv 060012; doi: https://doi.org/10.1101/060012

57. Dobin A, Davis CA, Schlesinger F, Drenkow J, Zaleski C, Jha S, Batut P, Chaisson M, Gingeras TR. STAR: ultrafast universal RNA-seq aligner. Bioinformatics. 2013 Jan 1;29(1):15–21. doi: 10.1093/bioinformatics/bts635. Epub 2012 Oct 25. PMID: 23104886; PMCID: PMC3530905.

58. SnapGene software (from Insightful Science; available at snapgene.com)

