## Supplemental Figures for "HIV-1 provirus transcription and translation in macrophages differs from pre-integrated cDNA complexes and requires E2F transcriptional programs"

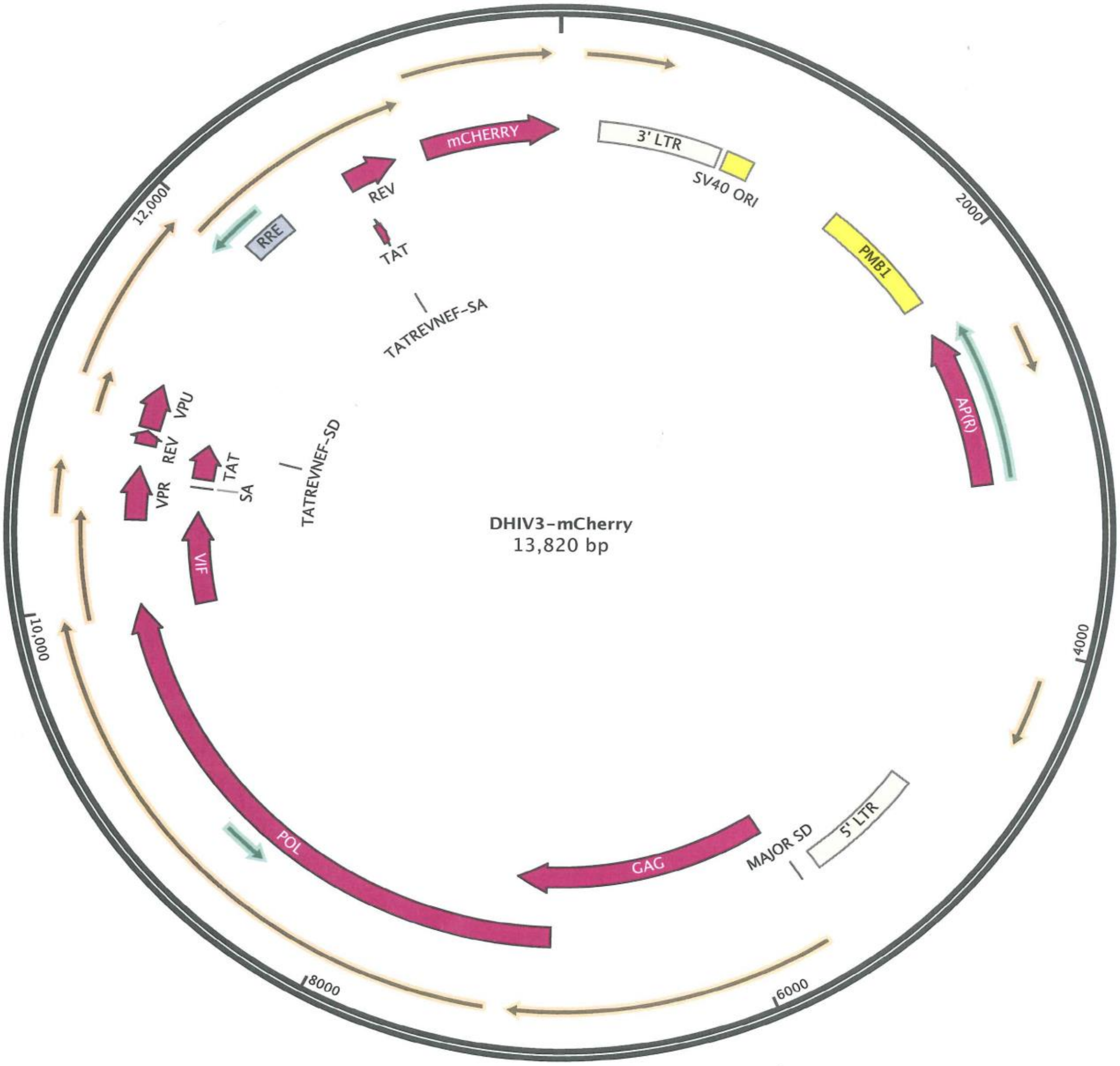


**S-1 DHIV3-mCherry map.** Snapgene [58] map of DHIV3-mCherry plasmid.

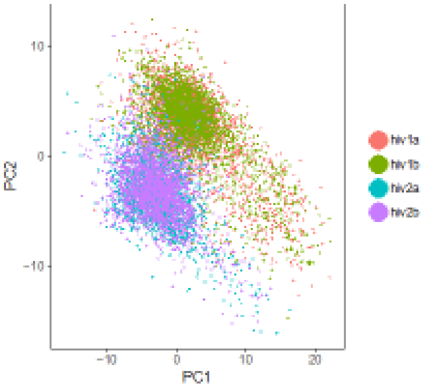


**S-2 Seurat analysis of biological repeats HIVreplicate1 and HIVreplicate2.**
HIVreplicate1a and HIVreplicate1b, and HIVreplicate2a and HIVreplicate2b, are technical repeat data. Technical repeats were conducted with each experiment. Figure shows PC analysis of biological repeat experiments. Technical duplicates were not different and so were combined for each repeat.

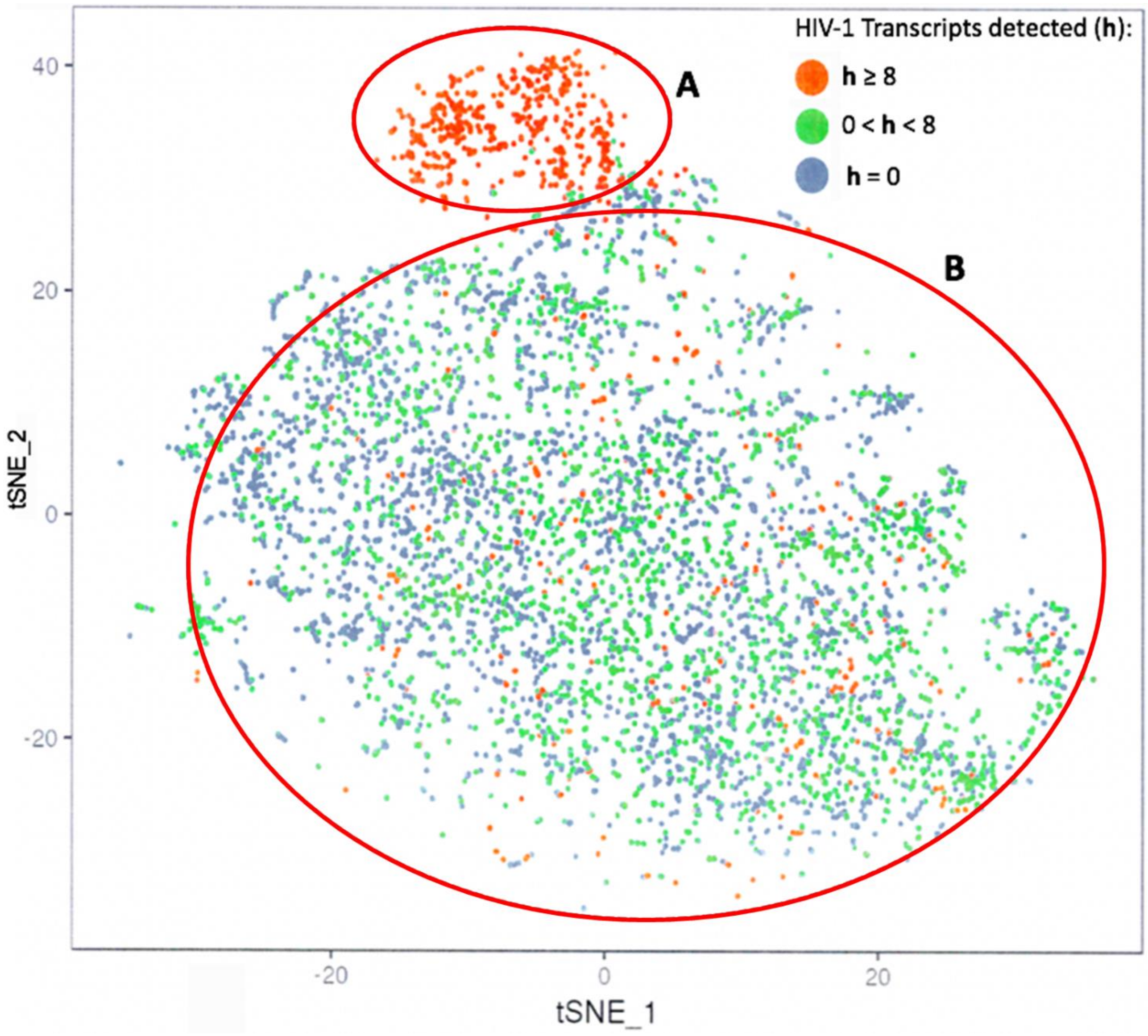


**S-3 tSne projection of scRNA seq data from experiment HIVreplicate1**. Seurat analysis and t-SNE projection of data shown in Figure 2. Viral transcript numbers (**h**) were determined for cells containing any detected HIV-1transcript, as described in methods. Orange dots represent high-level transcript load per cell, greater than 8 transcripts mapping to HIV-1 genes per cell, green dots indicate cells with lower transcript loads detected per cells, and blue dots indicate cells with no detectable HIV-1 transcripts. Barcodes of cells in Provirus Cluster (A) tracked to Provirus Cluster cells in UMAP analysis (Fig. 2).

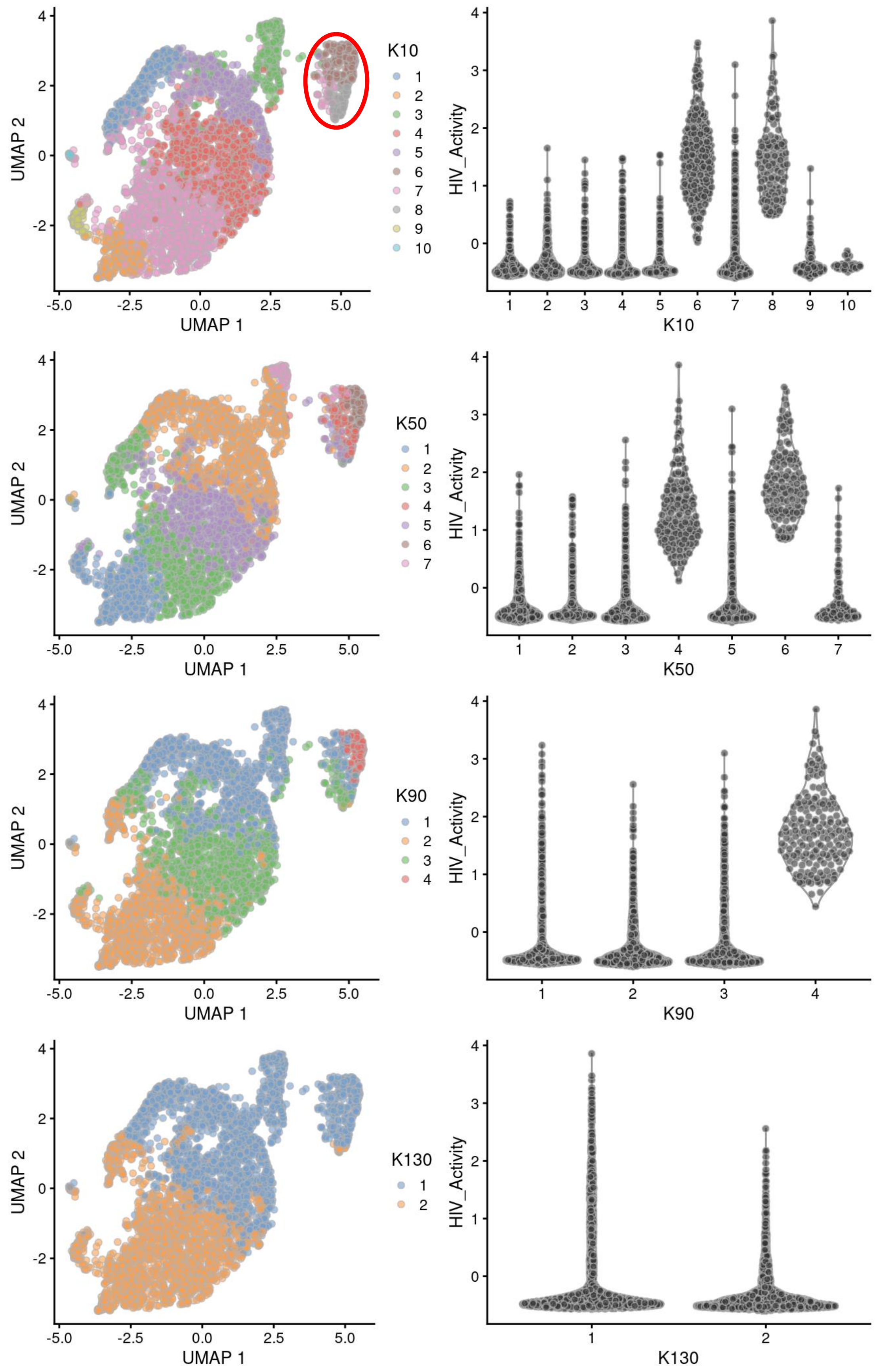
UMAP 1
UMAP 2
K10
1
2
3
4
5
6
7
8
9
10
HIV_Activity
K50
K90
K130

**S-4 Unsupervised clustering of UMAP shown in Fig 4.** Left panels show unsupervised clustering obtained at K values from 10 to 130. Right panels show Violin plots of HIV-1 transcripts/cell in the clusters identified at the specified K values (Scran's buildSSNGraph using the PCA as input). Clusters 6 and 8 at K equal to 10 contained most of the cells in the semi-supervised Provirus cluster (circled in red) and were used to define Provirus transcriptome, versus the remaining cells making up the semi-supervised PIC/Bystander cluster. Stipulation of lower K values means that during analysis any one given cell is clustered with a smaller number of cells with similar transcriptomes.

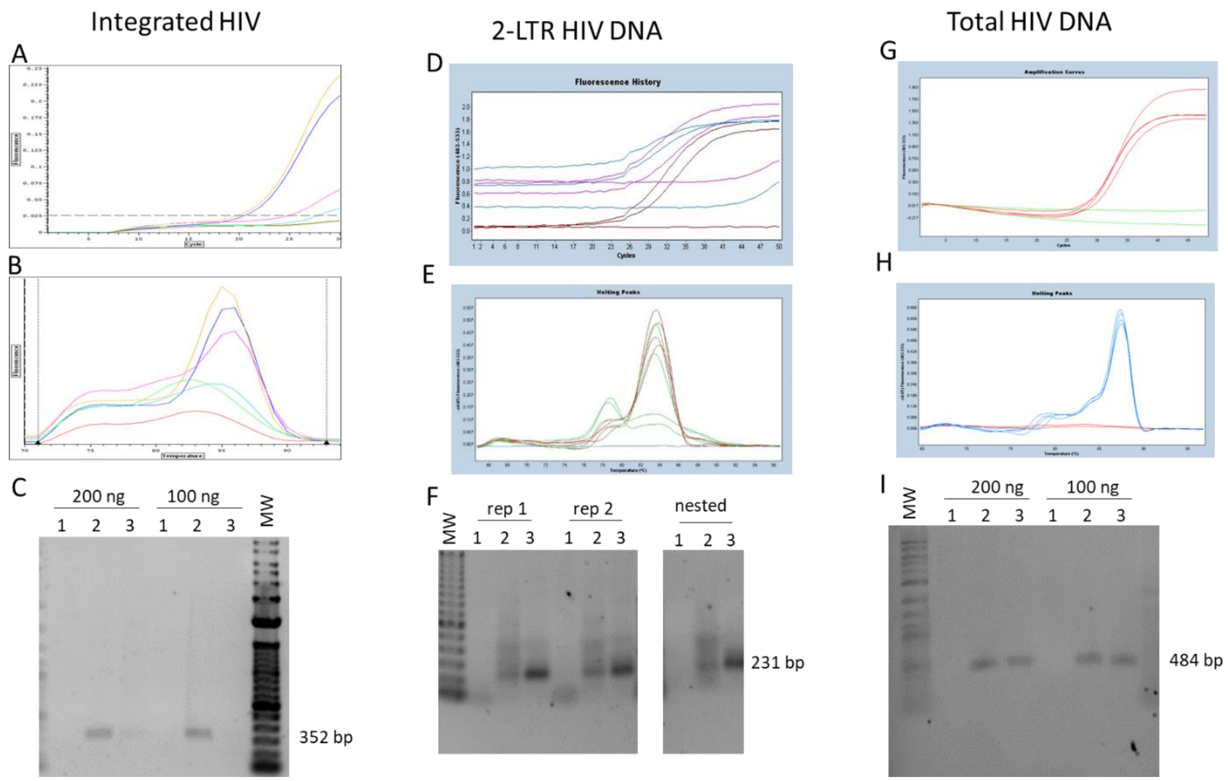


**S-5 Real-time PCR analysis of DNA samples from Control, DHIV3-mCherry infected, and DHIV3-mCherry infected, integrase inhibitor treated THP-1 cells.** MW, molecular weight markers. Lanes 1, Control cell DNA; Lane 2, DNA from DHIV3-mCherry infected culture cells; Lane 3, DNA from DHIV3-mCherry infected cultures treated with integrase inhibitor (25nM MK-2048). 100 ng of DNA was tested in each amplification unless noted. Primers used are described in Methods. Panels **A**, **B,** and **C**) PCR demonstration of integrated proviral HIV DNA. Panel **A**) examples of the progression curves; upper curve represent lanes 2 at 200 and 100 ng DNA respectively, middle curves reflect lanes 3, respectively, bottom 2 curves were generated by Control DNA . Panel **B**) melting curves; the upper curves represent lanes 2 at 200 and 100 ng and lane 3 at 200 ng DNA respectively. Panel **C**) shows the amplicons generated from the integrated DNA using the nested PCR strategy described by Chun et al. [34] on a 1% agarose gel. The amplicon product sizes matched the predicted product size of 352 bp. These examples were from two biological replicates, one using 200 ng and one starting with 100 ng of starting DNA purified using Qiagen Blood and Tissue DNeasy kits. The agarose gel shows the integrated proviral DNA, assessed using an MJ PTC-200 thermal cycler, and the nested PCR was evaluated using a Chromo-4 alpha unit. Note that the 200 ng samples with integrase inhibitor (Lane 3) show a small amount of integrated provirus DHIV3-mCherry DNA, demonstrating that the inhibitor did not completely inhibit the DHIV-mCherry integration. This is consistent with the 48 hr gag/p24 protein production seen in the immunoblot analysis (Fig. 7), and flow cytometry analysis (Fig. 6). Panels **D**, **E**, and **F**) show the same DNA samples used to detect 2-LTR circle PIC cDNA from the second or two consecutive PCR runs. Panel **D**) Lower 3 curves show progression curves with lack of 2LTR primer products in Control DNA, while all 6 biological repeats, 3 from DHIV3-mCherry infected cultures and 3 from DHIV3-mCherry infected cultures treated with integrase inhibitor show amplification of PIC cDNA p2LTR products. Panel **E**) shows the melting curves for these amplification products with the lower 3 curves representing Control DNA, the lowest curve representing Control DNA from the nested PCR approach (see Methods). Panel **F**) shows the amplicons generated run out on a 1% agarose gel. In these experiments, the PCR was assessed using a Roche LightCyler 480. Biological replicates of 100 ng starting DNA are represented as “rep 1” and “rep 2”, using the HIV F and R1 primers of Brussels and Sonigo [33]. In the confirmation experiment lanes, under the “nested” label in the agarose gel, wider bracketing primers were used in the first amplification followed by the HIV F and R1 primers “nested” in the second run. Perhaps not surprising after 100 cycles, there are contaminating PCR products in the Control lanes; however, the expected 231 bp amplicon is not detectable in the Control cell DNA, while it is the predominant product in DNA from either infected, or infected and integrase inhibitor treated cell DNAs. Panels **G, H,** and **I**) show the same DNA samples used to detect total DHIV-mCherry DNA, also assessed using the Roche LightCyler 480. Primers used are described in Methods. Panel **G**) shows the progression curves generated in this experiment, the lowest 2 curves represent Control cell DNA. Panel **H**) shows the melting curves for these products, again the lowest 2 curves are from Control cell DNA. Panel **I**) shows the amplicons generated, of predicted size, run out on a 1% agarose gel. The real-time PCR results show roughly equivalent total amounts of DHIV3-mCherry in infected culture DNAs, whether in the presence of integrase inhibitor or not. This indicates that overall, the total amounts of PIC cDNA are similar in integrase inhibitor treated and untreated DHIV3-mCherry infected cultures.

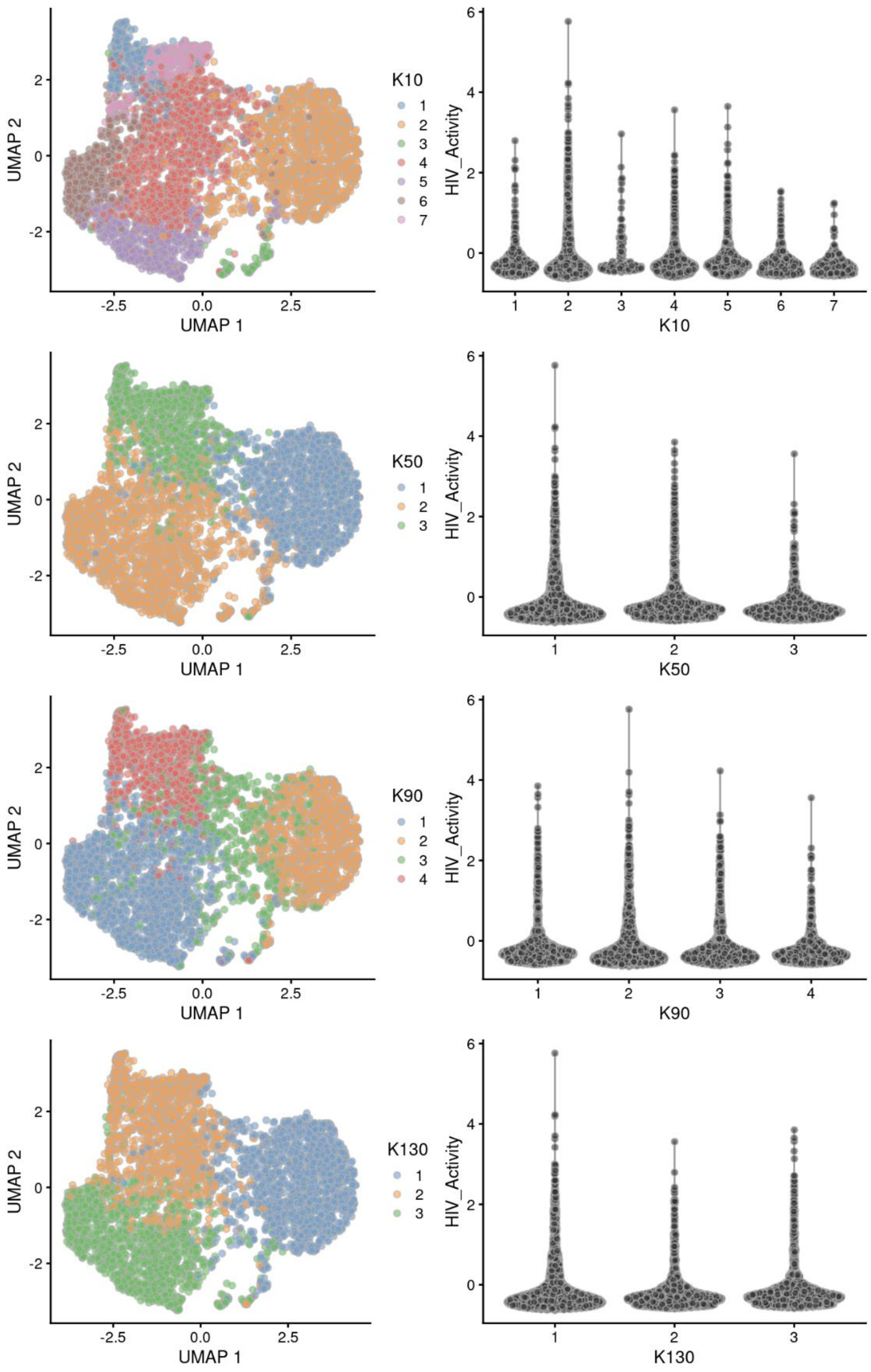

UMAP 2
UMAP 1
K10
1
2
3
4
5
6
7
HIV_Activity
K50
K90
K130
-2.5
0.0
2.5

**S-6 Unsupervised clustering of integrase-inhibitor-treated DHIV3 infected cells from Figure 8.** No cluster corresponding to the Provirus cluster identified in HIVreplicate1 or HIVreplicate2 could be identified, regardless of K value specified. Data were analyzed as in Figure 4. Left panels show unsupervised clustering obtained at K values from 10 to 130. Right panels show Violin plots of HIV-1 transcripts/cell in the clusters identified at the specified K10 values (Scran's buildSSNGraph using the PCA as input). Stipulation of lower K values means that during analysis any one given cell is clustered with a smaller number of cells with similar transcriptomes.

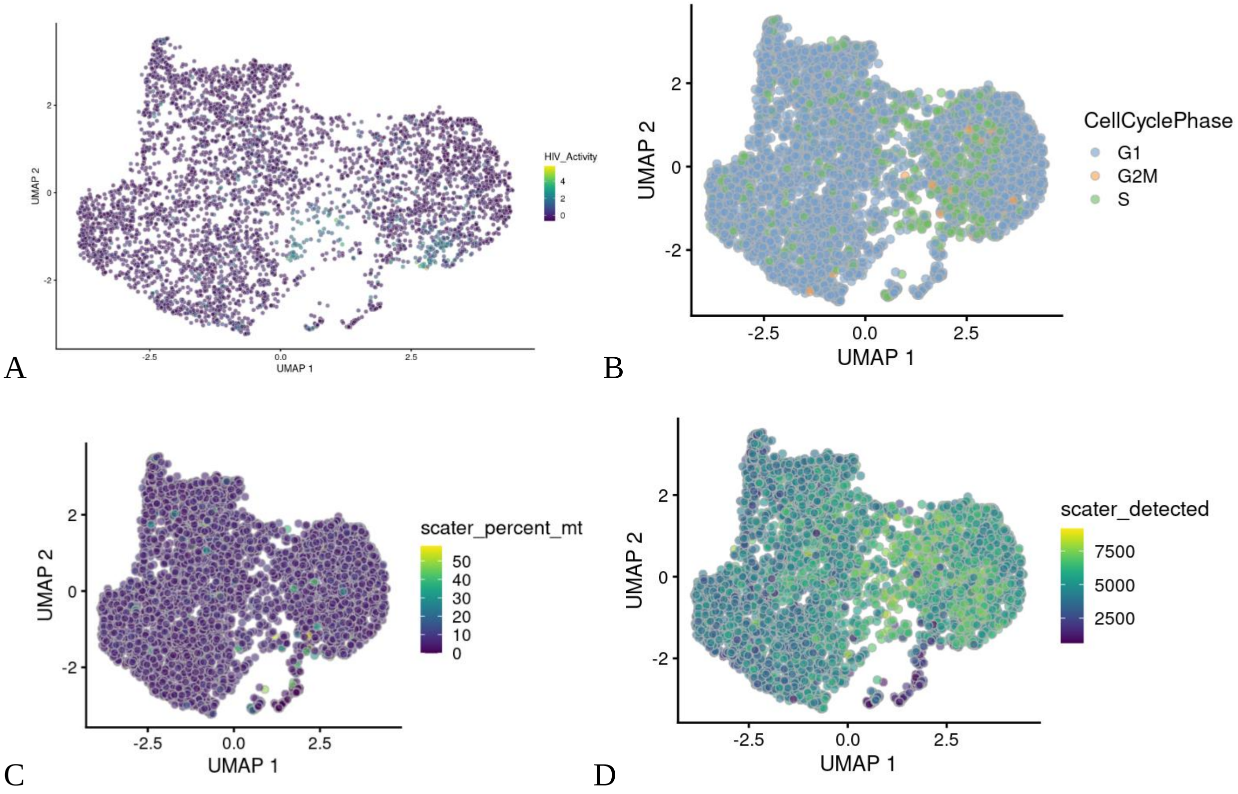


**S-7 Feature plots of integrase inhibitor-treated cultures.** Transcripts of DHIV3-mCherry infection readily detectable in the presence of integrase inhibitor Panel **A**. Effects of cell cycle (Panel **B)**, mitochondrial gene expression (Panel **C**), and number of genes detected per cell (Panel **D**) shown for comparison to Figure 3.

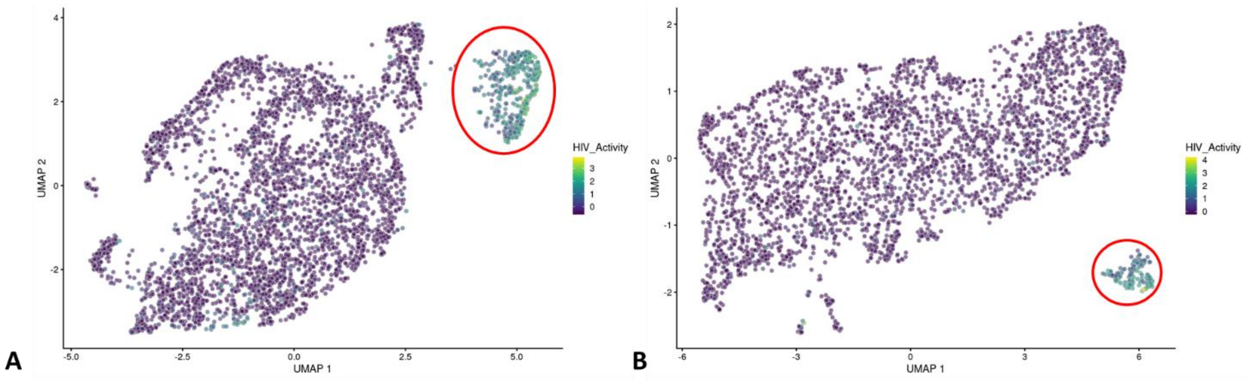


**S-8 Biological repeat experiments HIVreplicate1 and HIVreplicate2.** UMAP projections of Seurat analysis of biological repeat experiments HIVreplicate1 (Panel **A**) and HIVreplicate2 (Panel **B**). Seurat analysis yielded 8.1% of cells in Provirus cluster from experiment HIVreplicate1, 6% of cells in Provirus cluster in repeat HIVreplicate2, in agreement with percentages of mCherry positive percentages obtained for duplicate cultures analyzed by Flow Cytometry.

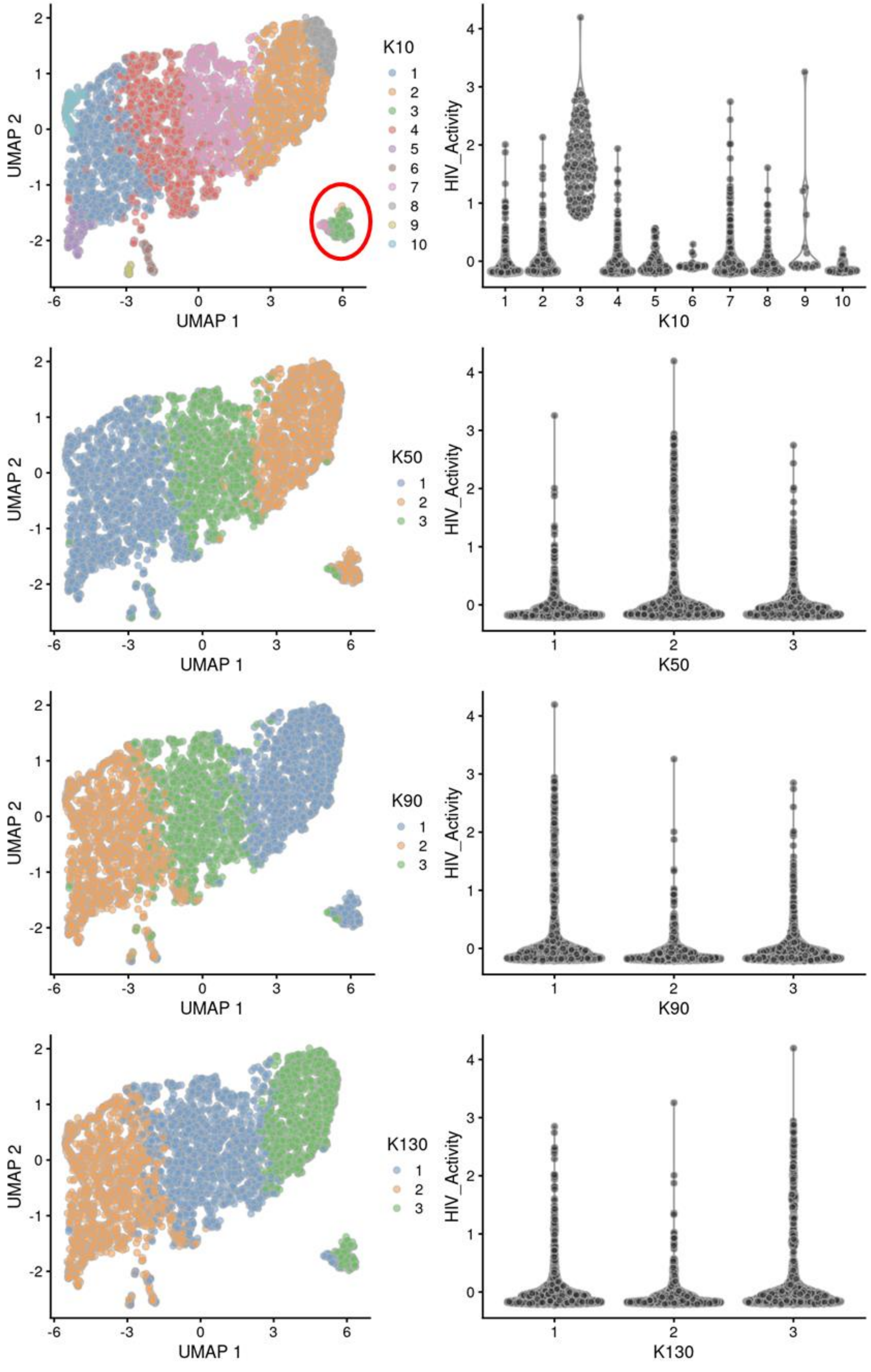
UMAP 1
UMAP 2
K10
1 2 3 4 5 6 7 8 9 10
HIV_Activity
K50
K90
K130
1 2 3

**S-9 Unsupervised clustering of HIVrepeat2 UMAP projection.** Left panels show unsupervised clustering obtained at K nearest neighbor values from 10 to 130. Right panels show Violin plots of HIV-1 transcripts/cell in the clusters identified at the specified K values (Scran's buildSSNGraph using the PCA as input). Cluster 3 at K equal to 10 contained most of the cells in the semi-supervised Provirus cluster (circled in red) and was used to define Provirus transcriptome, versus the remaining cells making up the semi-supervised PIC/Bystander cluster. Stipulation of lower K values means that during analysis any one given cell is clustered with a smaller number of cells with similar transcriptomes.

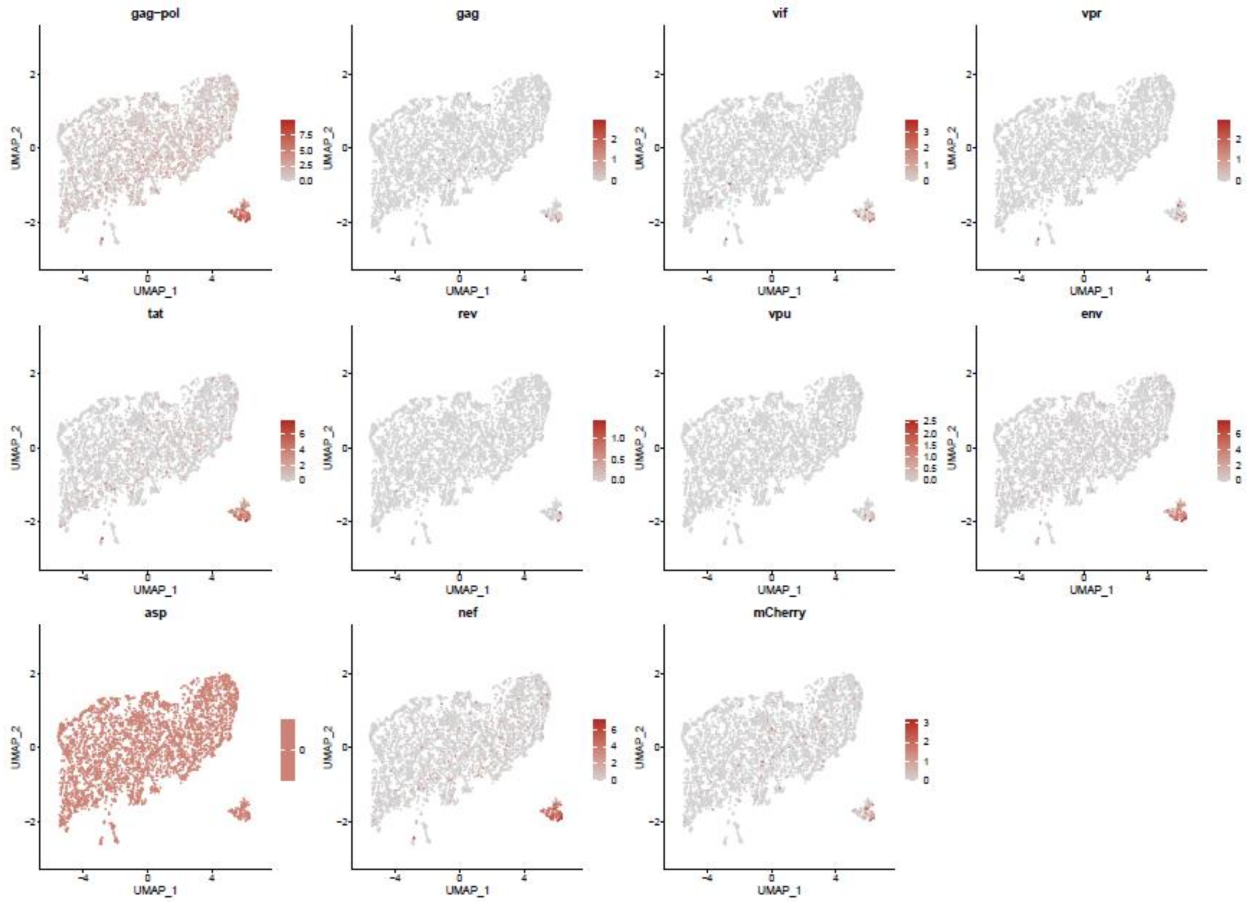


**S-10. The Distribution of HIV-1 transcripts throughout Provirus and PIC/Bystander clusters of HIVreplicate2.** Feature plot showing the distribution of cells from UMAP in Fig. S-9 containing detectable DHIV3-mCherry transcripts. As described above, these UMAP projections were made with Seurat's FeaturePlot function. They are colored by expression of individual genes (UMAP projection colored by walktrap, normalized log2 values). ASP is a negative control, bacterial gene transcript sequence. The distribution repeats the results obtained in HIVrepeat1 (Fig. 10 A).

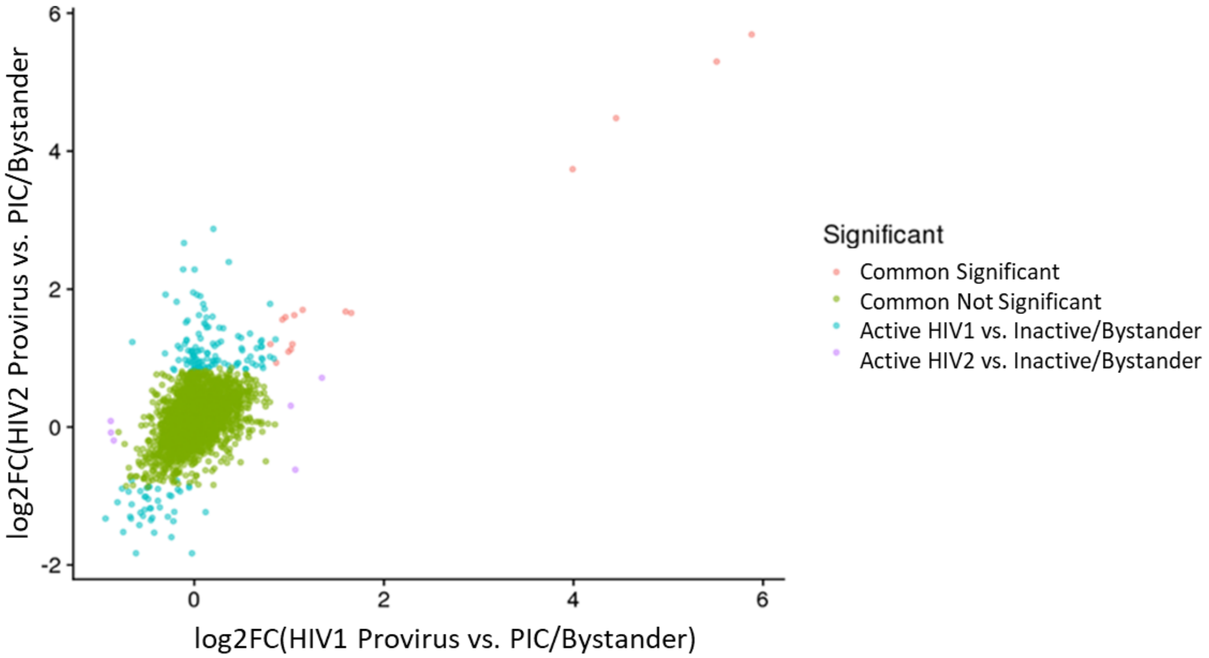


**S-11 Differential gene expression comparison of Provirus and PIC cluster gene transcripts from biological repeat experiments.** Consistent positive correlation of common DGE in HIVreplicate1 (abscissa) versus HIVreplicate2 (ordinate) repeat experiments (Spearman's rank correlation coefficient of all common genes 0.384), agreed with Hallmark and REACTOME analyses that showed similar pathways up- or down-regulated in the Provirus versus PIC/Bystander clusters of the biological repeat experiments.

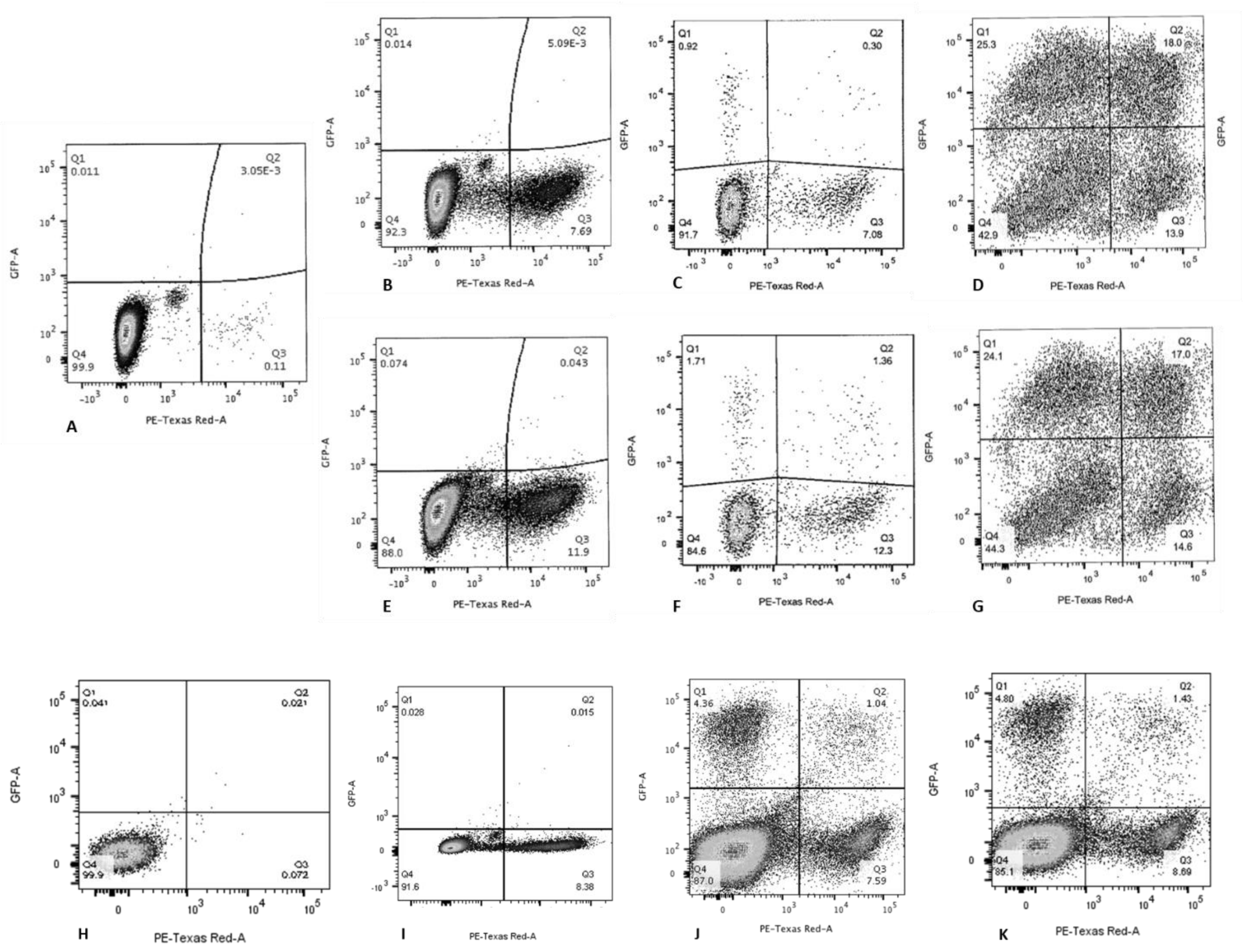


**S-12 Sequential infections of primary human lymphocyte and macrophage cultures.** Provirus, mCherry positive, cells were 2 to 5 times more likely to make HIV-1 encoded GFP protein upon second infection than PIC/Bystander cells upon second infection. Panels **A**, **H**) primary cultures of T-lymphocytes and macrophages at time equals 0 hrs, respectively. Panels **B-D**) primary lymphocytes infected with low titer DHIV3. Panels **E-G**) primary lymphocytes infected with high titer. Panels **H-K**) primary macrophages. Percentage of mCherry cells also producing GFP, compared to cells producing mCherry only, is always 2 to 5 times higher than the percentage of cells making only GFP, compared to those cells not producing mCherry. Panel **A** and **H**) Time equal 0 hrs; addition of DHIV3-mCherry. Panel **B**, **E** and **I**) time equal 24 hrs; addition of DHIV3-GFP. Panel **C**, **F,** and **J**) time equals 48 hrs after DHIV3-mCherry addition, 24 hrs after DHIV3-GFP addition. Panel **D**, **G,**and **K**) time equals 72 hrs after DHIV3-mCherry addition, 48 hrs after DHIV3-GFP addition.
